# Androgen receptor imprints satellite cells stemness and preserves their reservoir for lifelong regeneration and optimal repair

**DOI:** 10.1101/2025.10.30.685031

**Authors:** Joe G. Rizk, Kamar C. Ghaibour, Sirine Souali-Crespo, Emilia Calvano, Aurore Bilger, Hugues Jacobs, Nadia Messaddeq, Qingshuang Cai, Erwan Grangirard, Rajesh Sahu, Arnaud Ferry, Valentin Fourcade, Joffrey Zoll, Gianni Zanardelli, Nacho Molina, Coralie Fontaine, Jean-François Arnal, Daniel Metzger, Delphine Duteil

**Affiliations:** Université de Strasbourg, CNRS, Inserm, IGBMC UMR 7104-UMR-S 1258, F-67400 Illkirch, France; Centre de Recherche en Myologie, UMRS974-Sorbonne Université-INSERM U974-Association Institut de Myologie, France; Université de Strasbourg, Centre de recherche en biomédecine de Strasbourg, UMR 3072, Strasbourg, France; Institut National de la Santé et de la Recherche Médicale (INSERM), U1297, Institut des Maladies Métaboliques et Cardiovasculaires (I2MC), Université Toulouse III Paul Sabatier, CHU Rangueil (University Hospital) de Toulouse, F-31432 Toulouse, France

**Keywords:** Satellite Cells, Androgen Receptor, Androgens, Skeletal Muscle, Regeneration, Ageing, Symmetric differentiation

## Abstract

Skeletal muscle stem cells (MuSC) are the guardians of muscle regeneration, sustaining tissue integrity through a delicate balance of quiescence, activation, and lineage commitment. While numerous molecular cues have been implicated in regulating these processes, the influence of androgen receptor (AR) signaling, an essential hormonal pathway for male muscle physiology, has remained largely unexplored. Here, we show that AR expression defines quiescent MuSC and acts as a safeguard of their dormancy. Integrated multi-omic analyses reveal a redistribution of AR binding from quiescence-maintenance loci to regulatory elements driving activation and metabolic reprogramming during repair. Loss of AR in young adult mice disrupts this balance, precipitating premature cell-cycle entry, skewed division modalities, depletion of the stem cell reservoir, and destabilization of the niche. These defects converge with hallmarks of aging-associated androgen decline, while androgen supplementation restores regenerative competence. Together, our findings establish AR signaling as a pivotal determinant of MuSC fate and a cornerstone of skeletal muscle homeostasis.

## INTRODUCTION

Skeletal muscles exhibit a remarkable regeneration capacity upon injury, resulting from the fusion of newly formed myocytes to generate myofibers that replace the damaged ones. This process is initiated by myofiber degeneration, causing inflammatory cell infiltration followed by the activation of satellite cells, which are the muscle stem cells (MuSC) ^1^. In the intact muscle, MuSC reside in a quiescent state within a specialized niche between the myofiber sarcolemma and the basal lamina ^2, 3^. They are defined by the expression of the myogenic transcription factor paired box 7 (*Pax7*), a key marker of their identity and function ^4^.

Upon muscle injury, MuSC exhibit a dual capacity. They can either self-renew to replenish the stem cell pool and preserve long-term regenerative competence, or enter a proliferative phase followed by activation of the myogenic differentiation program, culminating in fusion and formation of new myofibers ^5, 6^. These key myogenic features, are intrinsically dependent on MuSC microenvironment, comprising immune and stromal cells. This microenvironment is tightly regulated by interactions with extracellular matrix components, inter-cell signaling and paracrine or endocrine stimuli ^7–9^. With age, skeletal muscle regenerative capacities progressively decline, which has been recently attributed to a reduction in MuSC number and function ^10–12^. These processes are thought to be exacerbated by hormonal changes, including decrease in circulating androgens ^13–15^.

Androgens are steroid hormones mainly produced by male gonads. They play a crucial role in modulating muscle homeostasis and MuSC fate, both during postnatal development and at adult stages ^16, 17^. In young adult mice, androgens promote MuSC quiescence through a Notch-dependent pathway ^18^, and influence their activation, proliferation, and differentiation ^18–20^. In addition, androgen supplementation increases MuSC numbers in both animal models and humans ^21^. Oppositely, castration-induced androgen deprivation in rodents leads to perturbed MuSC homeostasis ^22^. Androgens exert their effects through the androgen receptor (AR), a ligand-activated transcription factor and member of the nuclear receptor superfamily, which upon ligand binding translocates to the nucleus and binds specific DNA sequences known as androgen response elements (AREs), to regulate target genes expression ^23–25^. We previously showed that AR in myofibers is instrumental for the musculoskeletal system by controlling gene networks required for structural and metabolic demands ^25, 26^. Nevertheless, the role of androgens *via* AR in MuSC during muscle regeneration, the broader consequences of androgen-dependent decline on the regenerative machinery, the underlying molecular mechanisms, and its potential causal link to age-associated muscle degeneration remain to be fully elucidated.

By integrating genome-wide profiling and histological analysis, with the study of an inducible MuSC-specific AR knockout mouse model and stem-like myogenic cell lines, we delineate the molecular mechanisms through which androgens *via* their receptor regulate MuSC regenerative functions. Our findings highlight the pivotal role of MuSC-AR in muscle repair, identifying AR-dependent stem cell profiles and supportive MuSC subtypes involved in postnatal myogenesis. We further reveal that AR orchestrates stemness transitions by sequential chromatin repositioning following injury, thereby controlling gene networks governing MuSC self-renewal, proliferation, and differentiation. Additionally, we demonstrate that androgen signaling contributes to the age-related decline in muscle healing capacities, and validate the translational relevance of murine findings in human myoblasts.

## RESULTS

### AR deficiency in MuSC in young individuals compromises regenerative capacity and alters metabolic and structural transcriptional programs in skeletal muscle

To dissect the age-dependent role of AR signaling within MuSC, we generated mice with a conditional AR deletion in this cell lineage (hereafter AR^(i)sat-/Y^ mice). Nine-week-old mice harboring floxed *Ar* alleles and expressing the *Pax7*-CreER^T2^ transgene were treated with tamoxifen (TAM) for 5 days, enabling selective deletion of *Ar* in MuSC (Figures S1A-S1D). TAM-treated littermates lacking the *Pax7*-CreER^T2^ transgene served as controls. To evaluate the functional consequences of AR ablation on skeletal muscle regenerative capacities, we subjected young (3-month-old) control and AR^(i)sat-/Y^ mice to acute Tibialis Anterior (TA) muscle injury *via* cardiotoxin (CTX) injection one month after gene ablation, and assessed histological features of repair at 5-, 7-, 14-, 28-and 42-days post-injury (dpi). While no gross phenotypical abnormalities were noticed in the absence of injury (Figures S1E-S1G), At 5 dpi, a timepoint at which MuSC activation and early differentiation culminate ^27^, both genotypes showed similar hallmarks of effective regeneration, including centronucleated fibers, necrosis, and immune cell infiltration (Figure S1H). However, at 7 dpi, AR^(i)sat-/Y^ mice exhibited regenerative defects, including persistent inflammation, and aberrant fiber morphology that persisted until 42 dpi, that were not due to the CreER^T^^2^ knock-in (Figures 1A and S1H). Ultrastructural analyses performed at 7 dpi, a timepoint at which mononuclear cells fuse in order to form myosyncytia and primary regenerative outcomes are captured ^28^, revealed that AR^(i)sat-/Y^ muscles displayed disorganized sarcomeres, with misaligned triads characterized by dilated T-tubules, disrupted Z-lines, and poorly defined or absent M-lines (Figure 1B). These structural muscle defects correlated with a significant reduction in myofiber cross-sectional area (Figure 1C), which translated into a significant reduction in contractile force at 28 dpi (Figure 1D). Muscle dysfunction was accompanied by a marked reduction in the MuSC pool at 7 dpi (Figures 1E and 1F), which persisted at 28 dpi (Figures S1I and S1J), showing that loss of AR in MuSC severely impairs regenerative capacity in young male mice.

**Figure 1:**
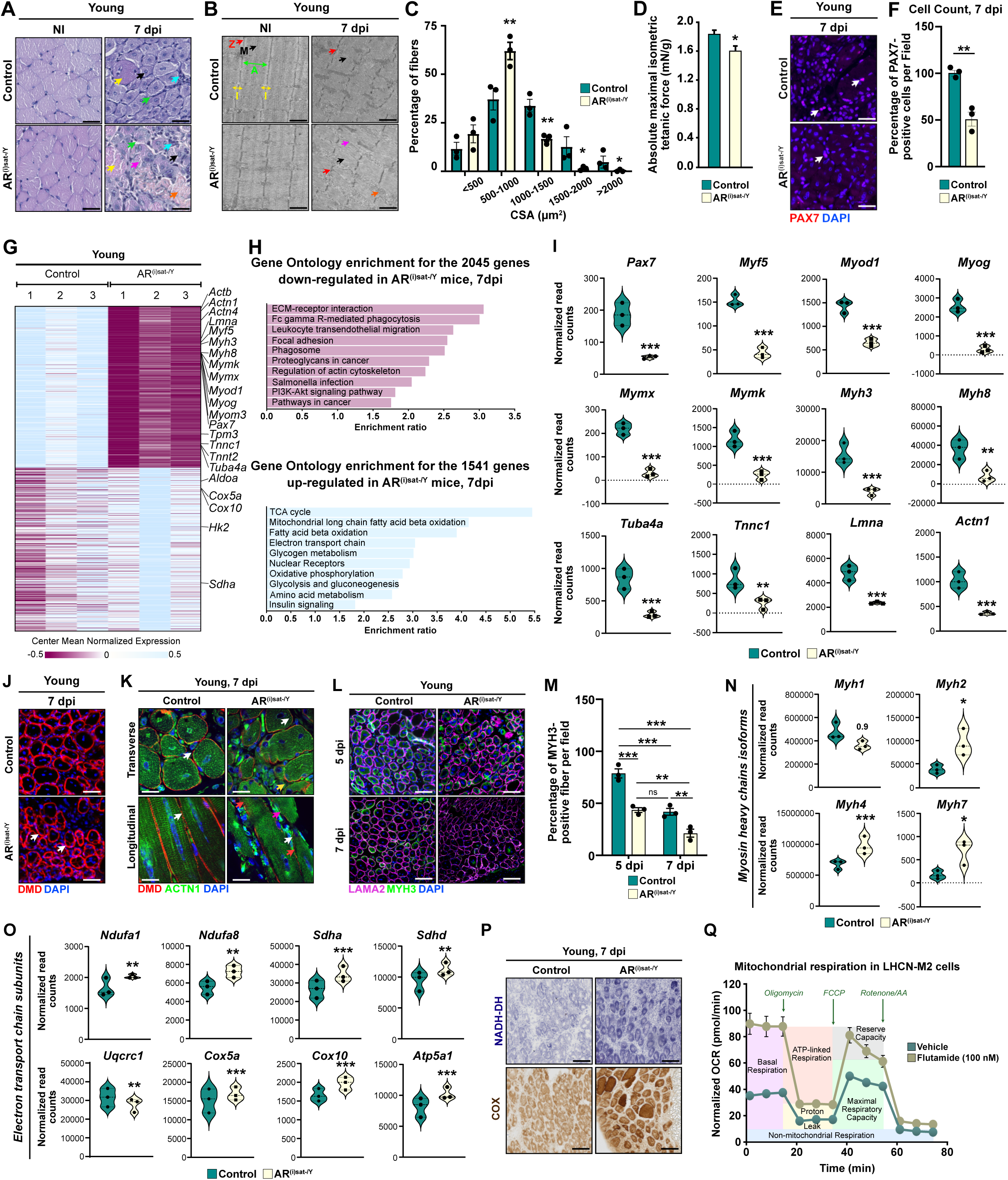
Absence of MuSC-AR impairs muscle regenerative capacities at young age. **(A)** Representative hematoxylin and eosin (H&E) staining of tibialis anterior (TA) muscles of young control and AR^(i)sat-/Y^ male mice under non-injured (NI) condition, and at 7 days post-injury (7 dpi). Black arrows indicate centrally nucleated fibers. Cyan arrows point to immune infiltration. Yellow arrows denote necrotic fibers. Green arrows show interstitial space. Pink arrows label misshaped regenerating myofibers. Orange arrows mark fibrosis. Scale bars, 75 µm. **(B)** Ultrastructure analysis of NI and 7-days-injured TA of young control and AR^(i)sat-/Y^ male mice. Z: Z-line, M: M-line, I: I-bands, A: A-band. Black arrows indicate M-line disturbances. Red arrows point to Z-line disruption. Pink arrows depict T-tubules misalignments. Orange arrows denote fibrosis. Green arrows show the A-band placement. Yellow arrows label the I-bands placements. Scale bars, 1 µm. **(C)** Distribution of myofibers cross-section area (CSA) in TA muscles of young control and AR^(i)sat-/Y^ male mice 7 dpi. Data are presented as mean ± SEM. Statistical test used Two-way ANOVA with Sidak’s correction; * = p < 0.05; ** = p < 0.01. **(D)** *In situ* single-fiber isometric force of 28-days-injured TA of young control and AR^(i)sat-/Y^ male mice. Data are presented as mean ± SEM. Statistical test used was two-tailed Mann-Whitney test; * = p < 0.05. **(E-F)** Representative immunofluorescent labeling of PAX7 (in red) **(E)**, and corresponding quantification of the number of PAX7-positive cells **(F)**, in 7-days-injured TA of young control and AR^(i)sat-/Y^ male mice. White arrows point to PAX7-positive cells. Nuclei were stained with DAPI. Data are presented as mean ± SEM. Statistical test used is two-tailed Mann-Whitney test; ** = p < 0.01. **(G-H)** Heatmap depicting the mean centered normalized expression of indicated genes selected from bulk RNA-sequencing analysis performed in 7-days-injured TA of young control and AR^(i)sat-/Y^ male mice **(G)**, and corresponding pathway analysis **(H)**. **(I)** Violin plots representing the normalized read counts of the indicated genes, selected from bulk RNA-sequencing analysis performed in 7-days-injured TA of young control and AR^(i)sat-/Y^ male mice. Statistical test used is Wald test; ** = p < 0.01; *** = p < 0.001. **(J)** Representative immunofluorescent detection of DMD (in red), in 7-days-injured TA of young control and AR^(i)sat-/Y^ male mice. Nuclei were stained with DAPI. White arrows depict fibers emerging within other fibers. Scale bars, 100 µm. **(K)** Representative immunofluorescent detection of DMD (in red), and sarcomeric alpha actinin (ACTN1) (in green), in 7-days-injured transversal and longitudinal sections of TA of young control and AR^(i)sat-/Y^ male mice. Nuclei were stained with DAPI. White arrows point to nuclei. Yellow arrows denote sarcolemma discontinuation. Red arrows show ACTN1 aggregates. Pink arrows denote stairs-shape in regenerating fibers. Scale bars, 50 μm. **(L-M)** Representative immunofluorescent detection of embryonic myosin (MYH3) (in green), and LAMA2 (in magenta) **(L)**, and corresponding quantification of the number of MYH3-positive cells **(M)** in 5-and 7-days-injured TA of young control and AR^(i)sat-/Y^ male mice. Nuclei were stained with DAPI. Scale bars, 85 μm. Data are presented as mean ± SEM. Statistical test used is Two-way ANOVA with Tukey correction; ns, non-significant; ** = p < 0.01; *** = p < 0.001. **(N-O)** Violin plots representing the normalized read counts of the indicated genes encoding myosin heavy chain (*Myh*s) isoforms **(N)**, or subunits of the electron transport chain complexes **(O)** selected from bulk RNA-sequencing analysis performed in 7-days-injured TA of young control and AR^(i)sat-/Y^ male mice. Statistical test used is Wald test; * = p < 0.05; ** = p < 0.01; *** = p < 0.001. **(P)** Representative histochemical staining of NADH dehydrogenase (NADH-DH) and COX activities in 7-days-injured TA of young control and AR^(i)sat-/Y^ male mice. Oxidative and intermediate fibers are darkly and moderately stained, respectively; glycolytic fibers are lightly stained. Scale bars, 200 μm. **(Q)** Mitochondrial respiration profile of vehicle and 100 nM flutamide-treated LHCN-M2 cells, assessed using the Seahorse Mito Stress Test, following sequential injection of oligomycin, carbonyl cyanide-p-trifluoromethoxyphenylhydrazone (FCCP), and rotenone/AA, and expressed as oxygen consumption rate (OCR). Data are normalized to total nuclear counts, and presented as mean ± SEM. Statistical test used was two-tailed Mann-Whitney test; * = p < 0.05.

To elucidate AR transcriptional landscape, we conducted bulk RNA sequencing (RNA-seq) on TA muscles harvested at 7 dpi from young AR^(i)sat-/Y^ and control mice. Our analysis revealed a differential expression of 3,586 genes, comprising 2,045 downregulated and 1,541 upregulated genes in AR-deficient mice (Figures 1G and S1K). Pathway analyses of the downregulated genes highlighted a strong association with biological processes involved in actin cytoskeleton organization, focal adhesion, and developmental signaling cascades such as the PI3K-Akt pathway (Figure 1H). Strikingly, the expression of key myogenic regulators, including *Pax7*, *Myf5*, *Myod1*, *Myog*, fusion-mediating factors such as *Myomixer* (*Mymx*) and *Myomaker* (*Mymk*) was markedly reduced in AR^(i)sat-/Y^ muscles (Figure 1I), in agreement with the decreased MuSC number. Similarly, transcripts encoding contractile cytoskeletal components such as actin, actinin, troponin and tropomyosin isoforms (e.g. *Actb*, *Actn1*, *Actn4*, *Tnnc1*, *Tnnt2*, *Tpm3*), and embryonic myosin heavy chains isoforms (*Myh3*, embryonic and *Myh8*, perinatal), as well as non-contractile components, including laminins (*Lmna*), *Myomesin 3* (*Myom3*), and *Dystrophin* (*Dmd*), were significantly diminished (Figures 1G and 1I), providing a molecular rationale for the morphological abnormalities observed at this timepoint. In agreement, immunofluorescence analysis of DMD at 7 dpi revealed profound structural abnormalities in AR^(i)sat-/Y^ mice compared to control littermates (Figure 1J). Moreover, co-staining for sarcomeric alpha actinin (ACTN1) and DMD on transverse sections showed disrupted sarcomere alignment and uneven ACTN1 distribution in mutant fibers, while longitudinal sections confirmed major striation defaults (Figure 1K, pink arrows). Notably, whereas healing control fibers had centrally positioned nuclei, AR^(i)sat-/Y^ fibers showed disorganized central nuclear localization (Figure 1K, white arrows), reflecting a disrupted cytoskeletal organization. Consistent with our transcriptomic data, immunostaining for embryonic myosin heavy chain (MYH3) at 5 and 7 dpi showed a ∼50% reduction in the number of MYH3-positive fibers in AR^(i)sat-/Y^ mice compared to controls (Figures 1L and 1M), signifying delayed or incomplete myogenic progression. To further investigate this impairment, we examined the expression profile of myosin heavy chain isoforms (Figure 1N). In contrast to the downregulation of *Myh3* and *Myh8*, transcripts of *Myh2*, *Myh4*, and *Myh7* (encoding MyHC IIa, IIb, and I, respectively) were upregulated (Figure 1N), reflecting a shift toward increased expression of adult myosin heavy chain isoforms at the expense of fetal ones, although *Myh1* (MyHC IIx) did not reach statistical significance (Figure 1N). Interestingly, transcriptomic analysis of the 1,541 genes upregulated in 7 dpi-injured AR^(i)sat-/Y^ TA muscles revealed a predominant enrichment in metabolic pathways, particularly those governing glycolysis, oxidative phosphorylation, mitochondrial function, and the metabolism of glycogen, fatty acids, and amino acids (Figure 1H). Notably, expression of several key regulators of cellular metabolism, including aconitase 2 (*Aco2*), citrate synthase (*Cs*), and multiple isoforms of isocitrate dehydrogenase (e.g. *Idh2* and *Idh3a*), was markedly elevated in AR^(i)sat-/Y^ muscles. This was paralleled by increased expression of genes and components of the oxidative phosphorylation machinery, including most of the members of mitochondrial complex I (e.g., *Ndufa1*-*13*), II (e.g., *Sdha*-*d*), III (e.g., *Uqcrc1*-*2*), IV (e.g., *Cox5a*, and *Cox10a*), and V (e.g., *Atp5a1* and *Atp5b*) (Figure 1O). Of note, the expression of these genes was not affected in the absence of injury (Figures S1L and S1M).

To investigate whether the absence of MuSC-AR during regeneration promotes a metabolic shift toward a more oxidative muscle fiber phenotype, we stained NADH dehydrogenase activity (NADH-DH) 7 dpi. In regenerating TA muscles from control mice, we observed lightly stained fibers, indicative of glycolytic metabolism. In contrast, AR^(i)sat-/Y^ muscles displayed a higher presence of darkly stained fibers (Figure 1P), reflecting increased NADH-DH activity. These findings were corroborated by cytochrome c oxidase (COX) activity staining, indicative of a glycolytic-to-oxidative shift in fiber type composition (Figure 1P). These findings prompted us to investigate whether such metabolic alterations are intrinsically rooted in MuSC. Seahorse analysis performed on LHCN-M2 human myoblasts, treated with 100 nM of the AR antagonist flutamide, unveiled an increased oxidative phosphorylation activity when compared to vehicle-treated cells (Figures 1Q and S1N) indicating that the metabolic abnormalities observed in regenerating myofibers may originate from intrinsic defects in MuSC. In parallel, basal ECAR was measured for glycolysis analyses (Figures S1O and S1P).

Altogether, our data show that AR loss in young individuals leads to MuSC depletion and important structural defects, associated with a fiber-type switch towards oxidative metabolism.

### Androgen signaling in MuSC coordinates gene programs essential for homeostasis and regeneration

To elucidate the cell-specific AR-dependent molecular mechanisms governing MuSC homeostasis during skeletal muscle regeneration, we performed a SMART-seq on FACS-isolated MuSC from 7 dpi-injured TA of young (15-week-old) control and AR^(i)sat-/Y^ male mice. Transcriptomic analysis revealed differential expression of 821 genes in AR-deficient MuSC, including 327 down-and 494 upregulated genes (Figures 2A and S2A). Consistent with the Seahorse assay findings and the histological evidence of a metabolic shift toward oxidative metabolism at 7 dpi, analysis of the upregulated genes highlighted a strong enrichment of pathways related to metabolic processes, particularly cellular respiration and oxidative phosphorylation (Figure 2B). In particular, the expression of *Cox5b* and *Cox6b1*, in addition to numerous genes implicated in mitochondrial homeostasis, was significantly elevated in AR^(i)sat-/Y^ MuSC (Figure 2C). Conversely, pathway analysis of the downregulated gene set revealed a significant association with cell cycle regulation (e.g., DNA replication and cell division) (Figures 2B and 2D).

**Figure 2:**
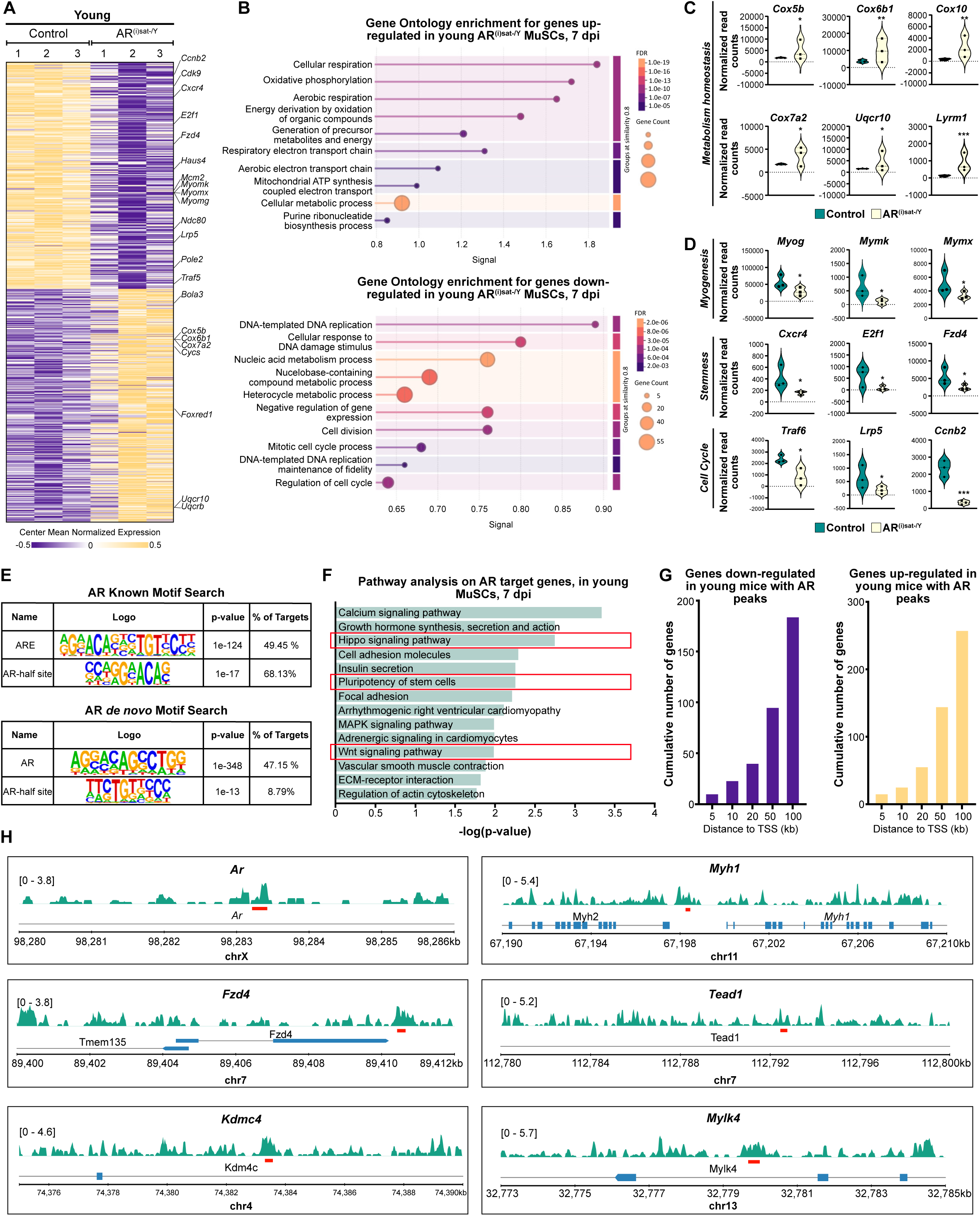
Characterization of AR cistrome and transcriptome in MuSC at young age. **(A)** Heatmap depicting the mean centered normalized expression of indicated genes selected from bulk RNA-sequencing analysis performed in 7-days-injured TA of young control and AR^(i)sat-/Y^ male mice. **(B)** Pathway analysis of down-and up-regulated genes in 7-days-injured TA of young control and AR^(i)sat-/Y^ male mice. **(C-D)** Violin plots representing the normalized read counts of the indicated genes, selected from bulk RNA-sequencing analysis performed in 7-days-injured TA of young control and AR^(i)sat-/Y^ male mice. Statistical test used is Wald test; * = p <0.05; ** = p < 0.01; *** = p < 0.001. **(E)** HOMER known and *de novo* motif analyses of AR-bound DNA sequences from 7-day-injured TA of young male mice. **(F)** Pathway analysis performed on AR target genes identified by CUT&RUN in MuSC of 7-day-injured TA of young male mice. Pathways of interest are squared in red. **(G)** Cumulative number of down-regulated genes, and up-regulated genes in MuSC of 7-day-injured TA of young male mice, with an AR peak located at the indicated distances from their TSS. **(H)** AR localization at the *Ar*, *Fzd4*, *Kdmc4*, *Myh1*, *Tead1*, and *Mylk4* loci, with binding sites shown in red.

To identify the direct transcriptional targets of AR, we isolated MuSC from TA muscles of control mice by FACS at 7 dpi, and performed CUT&RUN assays using AR-specific antibodies, alongside profiling of demethylated histone H3 (H3K4me2), a well-established marker of active promoters and enhancers. Bioinformatic analysis identified 7,346 AR-binding peaks, predominantly located within intronic and intergenic regions encompassing the canonical androgen response elements (AREs) (Figures 2E and S2B), as well as 39,900 peaks associated with the H3K4me2 mark, evenly distributed between promoter, intron and intergenic regions (Figure S2C). Integrative analysis of the two datasets revealed that AR is directly recruited to 3,104 transcribed genes (Figure S2D). Gene ontology and pathway enrichment analyses of AR-bound genes uncovered a strong association with key developmental and regenerative pathways, including the HIPPO (e.g. *Tead1*, *Wwc1*, *Fat4*) and WNT signaling cascades (e.g. *Wnt5a*, *Fzd4*), as well as signaling modules related to tissue remodeling and regeneration, such as cell adhesion, focal adhesion, extracellular matrix (ECM)-receptor interactions, and actin cytoskeleton regulation (Figure 2F). Notably, AR occupancy was also observed at genes essential for stem cell homeostasis (e.g. *Foxo1*, *Bcl2*), muscle repair (e.g. *Mstn*, *Cxcl2*, *Cdh15*), and sarcomere assembly or function (e.g. *Myh1*, *Mylk4*, *Myom2*). A transcription start site (TSS) gene-centric analysis demonstrated that AR binds more than the two-thirds of both down-and upregulated genes in AR^(i)sat-/Y^ mice within a 100-kb window (Figure 2G), showing that AR predominantly regulates gene expression through distal elements. Representative AR binding loci within key genes involved in the aforementioned pathways, including *Ar* itself, *Fzd4*, *Kdmc4*, *Myh1, Tead1*, and *Mylk4* are illustrated in Figure 2H. Gene ontology analysis revealed that genes downregulated in association with AR peaks at 7 dpi were predominantly involved in cellular structural and organelle-related processes (Figure S2E), whereas upregulated genes were enriched in metabolic pathways (Figure S2F), thereby providing a molecular basis for the impaired phenotype observed at the histological level. Altogether, these findings support a model in which AR orchestrates regenerative transcriptional programs in MuSC via enhancer-driven mechanisms.

### Single-cell RNA sequencing reveals early molecular and cellular alterations in the AR-deficient MuSC microenvironment

To decipher the primary events underlying defective MuSC homeostasis caused by AR loss, and the consequent impact on their microenvironment during impaired regeneration, we performed single-cell RNA sequencing (scRNA-seq) in 5-dpi-injured TA muscles of young (15-week-old) control and AR^(i)sat-/Y^ mice (Figure S3A). Of note, AR^(i)sat-/Y^ mice harbored the YFP transgene selectively in MuSC to efficiently follow those cells. We identified 12 unsupervised clusters, annotated to the literature ^29, 30^, based on the expression of specific marker genes (Figures 3A, S3A and S3B), with no evidence of cell population loss or emergence in mutants (Figures 3A, 3B and Table S1 and S2). However, we observed a slight reduction in the proportion of fibro-adipogenic progenitors (FAPs) and MuSC, as well as an increase in that of both macrophages in AR^(i)sat-/Y^ mice compared to controls (Figure 3B). Flow cytometry analysis of injured TA muscles at 5 dpi, furtherly confirmed a ∼30% increase in macrophages number among CD45^+^ immune cells (Figure 3C), and immunolabeling for F4/80 revealed an upward trend in macrophage abundance (Figure S3C and S3D). In parallel, flow cytometry analysis indicated an ∼8% decrease in MuSC, while total immune cells and FAPs were not significantly altered (Figure 3D). Notably, *Ar* expression was not ubiquitous across all populations; rather, it was predominantly detected in cell types exhibiting stem-like properties or mesenchymal profiles, including FAPs, fibroblasts, and MuSC (Figure 3E). *Ar* expression was completely abolished in mutant MuSC, and not in other cell types (Figure 3E), while *Yfp* was exclusively expressed in MuSC of AR^(i)sat-/Y^ mice, consistent with the genetic engineering of our model (Figure S3E). Moreover, AR deficiency in MuSC strongly altered the transcriptional profile of their cellular microenvironment (Figure S3F).

**Figure 3:**
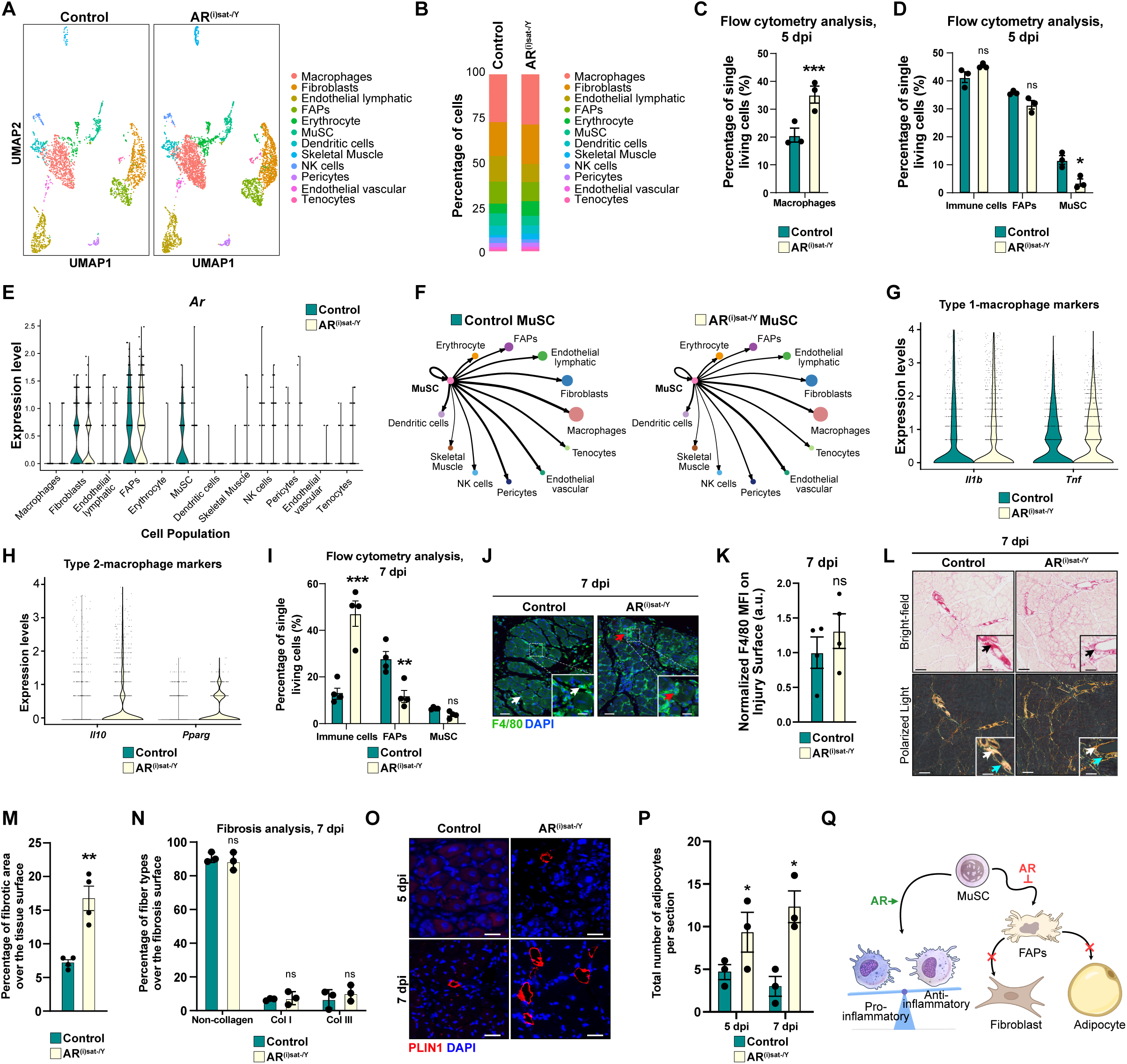
MuSC microenvironment is altered in the absence of AR. **(A-B)** Uniform manifold approximation and projection (UMAP) visualization of single cells **(A)**, and respective percentage of cell types **(B)**, from 5-days-injured control and AR^(i)sat-/Y^ TA of young mice, colored by cluster identity. **(C-D)** Percentage of macrophages **(C)**, immune cells, FAPs and MuSC **(D)**, in 5-days-injured TA of young control and AR^(i)sat-/Y^ mice, analyzed from flow-cytometry data conducted at the indicated time-point. Data are presented as mean ± SEM. Statistical test used was two-tailed Mann-Whitney test; ns, non-significant; * = p < 0.05, *** = p < 0.001. **(E)** Violin plot of *Ar* transcript levels in the indicated cell populations identified from 5-days-injured TA of young control and AR^(i)sat-/Y^ mice. **(F)** Schematic diagram of cell-cell communication between MuSC and other TA constitutive cells types in 5-days-injured TA of young control and AR^(i)sat-/Y^ mice. Arrow thickness indicates the relative interaction strength between populations, as inferred from ligand-receptor expression analysis. **(G-H)** Violin plots representing the transcript levels of macrophage type-1 **(G)**, and macrophage type-2 markers **(H)**, selected from scRNA-seq analysis performed in 5-days-injured TA of young control and AR^(i)sat-/Y^ mice. **(I)** Percentage of immune cells, FAPs and MuSC, in 7-days-injured TA of young control and AR^(i)sat-/Y^ mice, analyzed from flow-cytometry data conducted at the indicated time-point. Data are presented as mean ± SEM. Statistical test used was two-tailed Mann-Whitney test; ns, non-significant; ** = p < 0.01; *** = p <0.001. **(J-K)** Representative immunofluorescent detection of F4/80 (in green) **(J)**, and corresponding quantification of mean fluorescence intensity (MFI) of F4/80 signal **(K)**, in 7-days-injured TA of young control and AR^(i)sat-/Y^ mice. White arrows depict inter-fiber macrophages. Red arrows point to intra-fibers macrophages. Nuclei were stained with DAPI. Scale bars, 100 μm for the main, 30 μm for magnified inset. Statistical test used was two-tailed Mann-Whitney test; ns, non-significant. **(L)** Representative Sirius Red histochemical staining visualized under brightfield (upper panel), and polarized light (lower panel), in 7-days-injured TA of young control and AR^(i)sat-/Y^ mice. Black arrows point to collagen fibers deposition. White arrows depict type I collagen fibers. Cyan blue arrows indicate type III collagen fibers. Scale bars, 250 μm for the main, 90 μm for magnified inset. **(M-N)** Quantification of Sirius Red area from brightfield analysis **(M)**, and fibrosis composition **(N)** of 7-days-injured TA of young control and AR^(i)sat-/Y^ mice. Data are presented as mean ± SEM. Statistical test used was two-tailed Mann-Whitney test; ns, non-significant; ** = p < 0.01. **(O-P)** Representative immunofluorescent detection of perilipin (PLIN1) (red) **(O)**, and corresponding quantification **(P)**, in 5-and 7-days-injured TA of young control and AR^(i)sat-/Y^ mice. White arrows point to adipocytes. Nuclei were stained with DAPI. Scale bars, 100 μm. Data are presented as mean ± SEM. Statistical test used Two-way ANOVA with Sidak’s correction; * = p < 0.05. **(Q)** Schematic overview of the influence of AR signaling in MuSC within the regenerative niche at young age. MuSC-AR contributes to the orchestration of the immune response during muscle regeneration by ensuring the timely coordination of pro-and anti-inflammatory phases. In parallel, MuSC-AR signaling restrains FAPs from excessive extracellular matrix deposition and adipogenic differentiation. Green arrows indicate activation, red arrows indicate inhibition, and black arrows represent undefined interactions.

To assess intercellular communication within injured muscle tissue at 5 dpi, we employed CellChat analysis on scRNA-seq data. In control mice, this analysis revealed a highly interconnected signaling network encompassing all cellular populations, indicative of active and coordinated crosstalk during the regenerative process (Figures S3G and S3H). Interestingly, communication within the MuSC population was diminished in AR^(i)sat-/Y^ mice, as indicated by the decrease of the arrow thickness (Figure 3F), pointing to a disruption in intra-population signaling.

Given the critical interplay between MuSC, macrophages, and FAPs, we next examined the gene networks affected by MuSC-AR loss within these two cell populations. Functional enrichment analysis of the 1,707 differentially expressed genes upon AR loss in MuSC within macrophages uncovered a transcriptional landscape indicative of macrophages homeostasis (Figure S3I). In particular, transcripts encoding core components of the inflammasome pathway, including caspase 1 (*Casp1*) and its adaptor *Pycard*, were elevated, alongside the chemokines C-C motif chemokine ligand 2 (*Ccl2*) and C-X-C motif chemokine ligand 10 (*Cxcl10*), both of which are known to promote monocyte recruitment. Furthermore, we observed enrichment in metabolic and proteasome pathways intimately linked with immune regulation and tissue remodeling, including oxidative phosphorylation (e.g., *Cox6b1, Cox17*), and ubiquitin-proteasome mediated degradation (e.g., *Uba3*) (Figure S3I). Moreover, macrophage polarization analysis revealed robust expression of canonical pro-inflammatory markers including *Il1b* and *Tnf*, in both control and mutant macrophages at 5 dpi (Figure 3G). In contrast, macrophages from AR^(i)sat-/Y^ mice expressed anti-inflammatory markers such as interleukin-10 (*Il10*) and peroxisome proliferator-activated receptor gamma (*Pparg*) (Figure 3H), suggesting a precocious transition toward an anti-inflammatory phenotype in mutant individuals. Further subclustering of the macrophage population revealed that they did not constitute a homogeneous cluster, but instead, segregate into five transcriptionally distinct subpopulations with specialized functional programs (Figures S3J-S3M). Cluster 0 was characterized by high expression of pro-inflammatory markers such as *Ccr2*, *Il1b*, and *Ccl5*, consistent with recently recruited, monocyte-derived macrophages (Figures S3J-S3L). Cluster 1 displayed a strong pro-inflammatory profile, marked by elevated expression of *Tnf*, *Ccl2*, *Ccl7*, *Cxcl1/2/3*, interferon-stimulated genes (*Ifit1/2/3*), indicative of highly activated macrophages involved in leukocyte recruitment and inflammatory amplification (Figures S3J-S3L). Cluster 2 corresponded to phagocytic and lipid-associated macrophages, expressing canonical LAM markers including *Gpnmb*, *Ctsd*, *Spp1*, *Fabp5*, *Lgals3*, and *Cd36*, suggesting a central role in debris clearance and lipid handling during tissue remodeling (Figures S3J-S3L). Cluster 3 was enriched for genes involved in iron metabolism and oxidative stress responses, such as *Slc40a1*, *Hmox1*, *Fth1*, and *Ftl1*, together with *Marco*, *Spic*, and *Arg1*, consistent with specialized macrophages implicated in iron recycling and stress adaptation (Figures S3J-S3L). Finally, Cluster 4 exhibited a tissue-resident, reparative phenotype characterized by *Cd163*, *Lyve1*, *Mrc1*, *Retnla*, and *Folr2* expression, alongside extracellular matrix and remodeling genes including *Col1a1*/2, *Sparc*, and *Gas6*, indicating anti-inflammatory, pro-reparative functions (Figures S3J-S3L). Differential expression analysis across different populations further revealed cluster-specific functional alterations, with altered antigen-presenting capacity, impaired oxidative metabolism, and shifts toward migratory or scavenger states depending on the subpopulation (Figure S3L). scRNA-seq analysis of macrophage populations revealed an increased representation of an anti-inflammatory macrophage subcluster in AR^(i)sat-/Y^ muscles (Figure S3M). To further examine this observation at the injury site, we performed immunofluorescence labeling for CD163, a marker associated with anti-inflammatory, pro-reparative macrophages. Quantification revealed an increase in CD163 mean fluorescence intensity (MFI) in AR^(i)sat-/Y^ TA muscles at 5 dpi compared to controls (Figures S3N and S3O). These observations are consistent with our single-cell RNA sequencing analysis, which identified a macrophage cluster enriched for *Cd163*, *Mrc1*, and *Folr2* expression, indicative of tissue-resident, anti-inflammatory, and remodeling-associated macrophages (Figures S3J and S3K).

To determine whether this aberrant immune environment persists over time, we extended our analysis to 7 dpi. Flow cytometry revealed a three-fold increase in immune cells in AR^(i)sat-/Y^ TA relative to controls (Figure 3I), and immunofluorescent labeling of the macrophage-specific marker F4/80 exhibited an increasing trend of the signal (Figures 3J, white arrows, and 3K). Remarkably, macrophages were frequently localized inside degenerating muscle fibers of AR^(i)sat-/Y^ mice (Figure 3J, red arrows, and 3K), a phenomenon typically restricted to early regenerative stages, thus indicating delayed resolution of inflammation or prolonged clearance of necrotic tissue. CD163 levels were comparable between AR^(i)sat-/Y^ and control muscles, in contrast to the increased signal observed in mutants at 5 dpi (Figures S3N and S3O). This temporal pattern may suggest that macrophages in AR^(i)sat-/Y^ muscles transiently acquire reparative features at earlier stages of regeneration.

Together, these findings reveal that AR signaling in MuSC plays a pivotal role in modulating the inflammatory response within the injured muscle by impacting the macrophage transcriptomic repertoire.

To unravel the impact of AR loss in MuSC on FAPs transcriptional signature, we performed further functional enrichment analysis of their 976 differentially expressed genes (Figure S3F). Pathway analysis revealed upregulation of fibrosis-related transcripts, including *Tgfb*, alongside extracellular matrix (ECM) remodeling enzymes, such as *Adamts* isoforms, *Mmp19*, *Serpine1*, and *Cthrc1*. Moreover, we observed enhanced expression of key adipogenic regulators (*Zeb1*, *Cebpd*, *Socs1*, and *Socs3*). Functional enrichment also highlighted disruptions in metabolic pathways, notably oxidative phosphorylation (*Cox4i1*, *Cox5a*, *Cox6a1*) and aerobic respiration (*Aco2*, *Hadhb*, *Gls*, *Lipa*) (Figure S3P), pointing toward a shift in FAPs identity, from pro-regenerative to fibrotic and adipogenic. Interestingly, while FAPs numbers were comparable between genotypes at 5 dpi (Figure 3D), a two-fold reduction was observed in AR^(i)sat-/Y^ muscles by 7 dpi (Figure 3I). We therefore postulated that decline might be attributed to differentiation of FAPs towards the fibrogenic or adipogenic lineages. To this aim, we performed Sirius Red staining at 7 dpi on injured TA from control and mutant mice. Mutant muscles displayed a marked increase in collagen deposition, with fibrotic area nearly doubled compared to controls (Figures 3L and 3M, black arrows). These findings were corroborated by Masson’s Trichrome staining, which revealed a significant enrichment of collagen within the ECM at 7 dpi, with signs of early deposition already evident at 5 dpi in young AR^(i)sat-/Y^ mice (Figure S3Q). Polarized light microscopy revealed that the qualitative composition of the ECM, including collagenous components such as collagens type I (Figures 3L, white arrows and 3N) and III (Figures 3L, cyan blue arrows and 3N), and non-collagenous cones (e.g., fibronectin, laminins, elastin, tenascin-C, and proteoglycans) remained comparable between genotypes, indicating a quantitative rather than compositional remodeling. We further evaluated adipogenic conversion via perilipin-1 (PLIN1) immunostaining. At 5 dpi, AR^(i)sat-/Y^ muscles already showed early signs of adipocyte emergence, which further increased by 7 dpi, whereas control muscles remained largely devoid of PLIN1-positive cells (Figures 3O and 3P). Thus, our data demonstrate that AR loss in MuSC impacts FAPs differentiation capacities promoting fibrosis and adipogenesis.

Altogether, these results indicate that AR signaling in MuSC plays a central role in coordinating the regenerative niche by regulating the balance between pro-and anti-inflammatory immune responses and controlling FAP fate (Figure 3Q).

### AR deficiency in MuSC at young age perturbs their stemness signatures, disrupts their commitment, and impairs their differentiation potential

Strikingly, only few genes were differentially expressed in MuSC, suggesting that different cell populations might lay in this cluster (Figure S3F). To further explore the transcriptional changes between control and AR^(i)sat-/Y^ MuSC, we performed subclustering of MuSC following additional stringent quality control, yielding a final dataset of 445 MuSC (Figures S4A and S4B). Unsupervised clustering revealed six distinct MuSC subpopulations (Figure S4B), of which two clusters exhibited an immune signature characterized by the expression of *C1qc*, *Lyz2*, and *Cd68* (Figures S4B and S4C). Given recent findings describing these subsets as immuno-myoblasts in regenerating muscle ^29, 31^, we hypothesized that these clusters represent MuSC targeted for immune-mediated clearance.

Consequently, we excluded these populations from our analysis and re-clustered the remaining four subtypes based on their transcriptional profiles (Figures 4A, S4D, S4E and Table S3).

**Figure 4:**
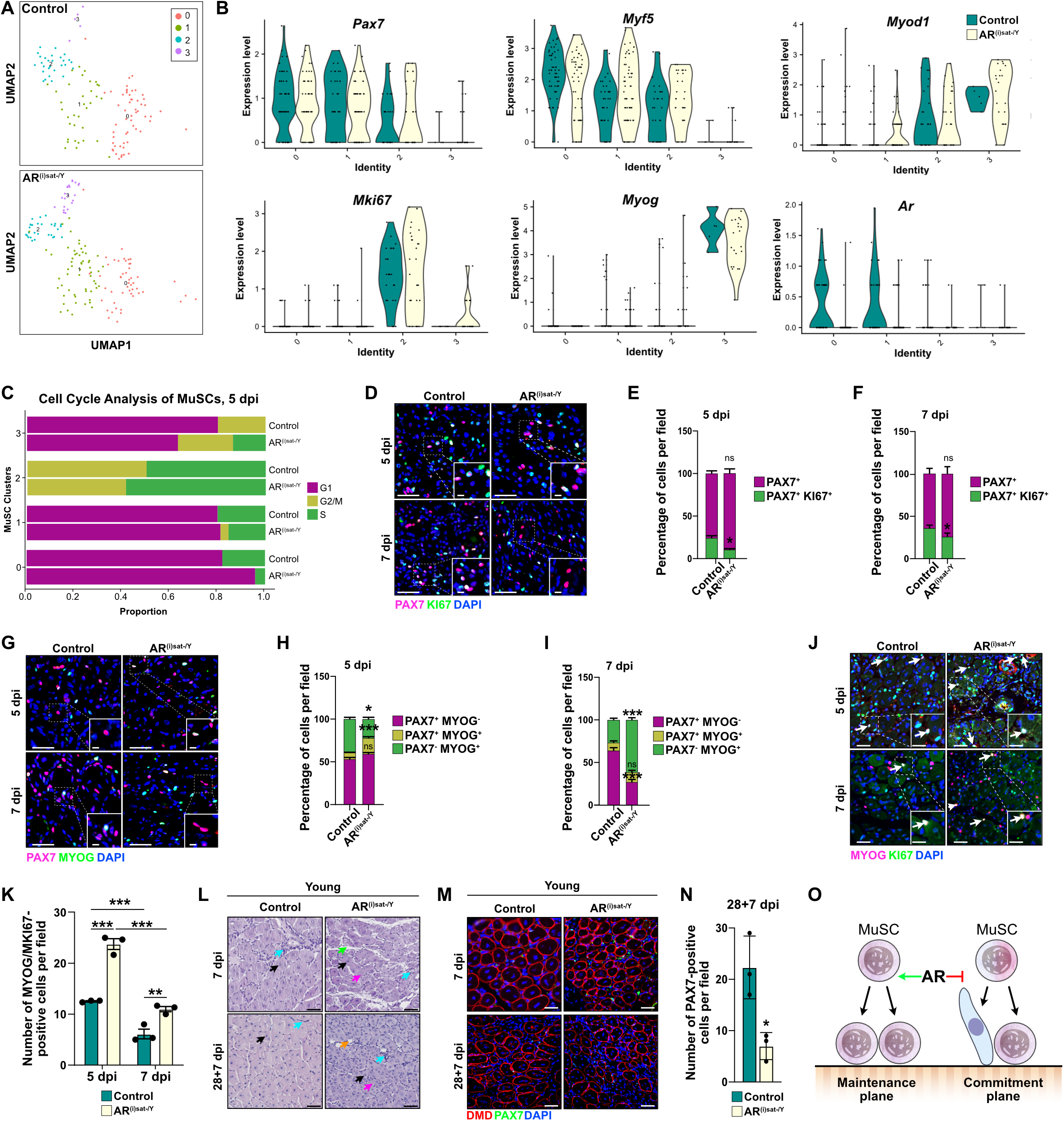
AR deficiency in MuSC favors symmetric differentiation over asymmetric division or symmetric self-renewal **(A)** UMAP visualization of MuSC reclusters from 5-days-injured control and AR^(i)sat-/Y^ TA of young mice, colored by cluster identity. **(B)** Violin plots representing the expression level of indicated genes in MuSC reclusters, selected from scRNA-seq analysis performed in 5-days-injured TA of young control and AR^(i)sat-/Y^ mice. **(C)** Cell cycle distribution across MuSC reclusters in 5-days-injured TA of young control and AR^(i)sat-/Y^ mice, showing the proportion of cells in G1, S, and G2/M phases. **(D-F)** Representative immunofluorescent detection of PAX7 (magenta), and KI67 (green) **(D)**, and corresponding quantifications in 5-**(E)** and 7-days **(F)** injured TA of young control and AR^(i)sat-/Y^ mice. Nuclei were stained with DAPI. Scale bars, 100 μm for the main, 30 μm for magnified inset. Statistical test used is Two-way ANOVA with Tukey correction; ns, non-significant; * = p < 0.05; *** = p < 0.001. **(G-I)** Representative immunofluorescent detection of PAX7 (magenta) and MYOG (green), and corresponding quantifications in 5-**(H)** and 7-days **(I)** injured TA of young control and AR^(i)sat-/Y^ mice. Nuclei were stained with DAPI. Scale bars, 100 μm for the main, 30 μm for magnified inset. Data are presented as mean ± SEM. Statistical test used is Two-way ANOVA with Tukey correction; ns, non-significant; * = p < 0.05; *** = p < 0.001. **(J-K)** Representative immunofluorescent detection of MYOG (in magenta) and KI67 (in green) **(J)**, and corresponding quantification of the number of MYOG/KI67-double positive cells per field of injury **(K)**, in 5-and 7-days-injured TA of young control and AR^(i)sat-/Y^ mice. White arrows depict MYOG/KI67-double positive cells. Nuclei were stained with DAPI. Scale bars, 75 μm for the main, 30 μm for magnified inset. Data are presented as mean ± SEM. Statistical test used is Two-way ANOVA with Tukey correction; ** = p < 0.01; *** = p < 0.001. **(L)** Representative HE staining of TA muscles of young control and AR^(i)sat-/Y^ mice, 7 (single injury) and 28+7 (double injury) days after injury. Cyan blue arrows denote inflammatory cells. Green arrows point to interstitial space. Black arrows depict central nuclei. Pink arrows show misshaped fibers. Scale bars, 100 µm. **(M)** Representative immunofluorescent detection of DMD (red), and PAX7 (green), of 28+7-days-injured young control and AR^(i)sat-/Y^ mouse TA. Scale bars, 50 µm. **(N)** Quantification of the number of PAX7-positive cells in 28+7-days-injured young control and AR^(i)sat-/Y^ mouse TA. Data are presented as mean ± SEM. Statistical test used was two-tailed Mann-Whitney test; * = p < 0.05. **(O)** Schematic overview of MuSC-autonomous regulation by AR signaling at young age. AR activity in MuSC promotes self-renewing divisions (maintenance plane), leading to the preservation of the stem cell pool. Conversely, AR signaling inhibits commitment divisions (commitment plane). Green arrow indicate activation, and red arrows indicate inhibition.

Our analysis delineated two distinct clusters of quiescent MuSC (clusters 0 and 1), both characterized by the expression of *Pax7* and *Myf5*, while lacking *Myod1* and *Myog* expression (Figures 4A, 4B and S4E). Moreover, we identified an activated MuSC subcluster (cluster 2) distinguished by the additional expression of *Myod1* and the proliferation marker *Mki67* (Figures 4A, 4B and S4E). Importantly, a fully committed myogenic cell population (cluster 3), characterized by *Myog* expression as well as that of *Mymk* and *Mymx,* and originally absent in control individuals, emerged in mutant mice (Figures 4A, 4B, S4E, and S4F). In line with this observation, we found that *Ar* expression was confined to quiescent clusters 0 and 1 in control mice, whereas ectopic *Myf5* and *Myod1* expression was noted in cluster 1 in AR^(i)sat-/Y^ individuals (Figure 4B), indicating that loss of AR in MuSC promotes myogenic commitment. Notably, several genes associated with symmetric division, including *Bmi1*, *Msi2*, and *Zfx*, were upregulated in mutant mice within cluster 3 (Figure S4G), alongside a broader expression of the fusogenic markers *Mymk* and *Mymx* across a larger subset of cells (Figure S4F), indicative of symmetric differentiation.

To provide evidence of commitment defaults caused by loss of AR in MuSC, we assessed cell cycle distribution within each cluster (Figures 4C and S4H). Our analysis revealed various perturbations upon AR loss, including cells in G2/M in cluster 1 (Figure 4C). Moreover, in the fully committed cluster 3, the proportion of cells in G1 dropped from ∼80% in controls to ∼60% in mutants, whereas those in S phase, absent in control mice, increased by more than 15% in MuSC-AR deficient mice (Figure 4C), indicating that they are more prone to mitotic activity.

To investigate the division modalities of MuSC, we performed a series of immunofluorescence analyses. Double staining for PAX7 and KI67 in injured TA muscles from control and AR^(i)sat-/Y^ mice at 5 and 7 dpi revealed a significant reduction in proliferating MuSC in mutant mice (Figures 4D-4F), indicative of impaired symmetric stem cell divisions. Furthermore, double immunodetection of PAX7 and MYOG at 5 dpi showed a marked increase in PAX7⁺/MYOG⁺ cells, accompanied by a decrease in PAX7⁺/MYOG⁻ cells upon AR loss in MuSC (Figures 4G and 4H). By 7 dpi, both PAX7⁺/MYOG⁺ and PAX7⁻/MYOG⁺ cell populations exhibited a two-fold increase in AR^(i)sat-/Y^ mice (Figures 4G and 4I), supporting a shift from symmetric stem cell divisions toward an asymmetric division program.

To determine whether the absence of AR in MuSC alters their mode of division, we performed MYOG and KI67 double immunostaining on injured TA muscles at 5 and 7 dpi. Notably, MYOG⁺/KI67⁺ double-positive cells were detected within regenerating regions of mutant muscles (Figure 4J), consistent with our scRNA-seq analysis showing an increased proportion of the committed cluster 3, characterized by *Mki67* expression in mutants (Figure S4I). Quantification confirmed that MYOG⁺/KI67⁺ cells were more abundant in AR^(i)sat-/Y^ muscles at both 5 and 7 dpi (Figure 4K), indicating that MuSC prematurely engage in the differentiation program. To further examine MuSC division behavior, single myofibers were isolated from injured TA muscles at 7 dpi and analyzed *in vitro* to assess the orientation and organization of satellite cell divisions. PAX7, KI67, and MYOG immunostaining revealed abnormalities in AR^(i)sat-/Y^ myofibers. Analysis of mitotic orientation on muscle fibers isolated from control and AR^(i)sat-/Y^ mice showed perpendicular MuSC division in mutant fibers (Figure S4J), indicative of a shift toward asymmetric differentiation, and suggesting disruption of the spatial control of MuSC division planes. In addition, mutant fibers exhibited nuclear budding-like structures along the fiber surface, a feature consistently absent in control fibers (Figure S4K). Overall, our data show that loss of AR in MuSC promotes premature differentiation and alters division orientation toward asymmetric divisions.

We henceforth hypothesized that increased symmetric differentiation may deplete the stem cell pool, compromising its ability to support effective repair during subsequent regenerative challenges. To test this, we subjected control and AR^(i)sat-/Y^ muscles to a secondary CTX-induced injury 28 days after the primary injury, and assessed tissue morphology 7 days later (28+7 dpi). At this stage, control muscles exhibited an accelerated regenerative response compared to muscles analyzed at 7 dpi following a single injury (Figure 4L), consistent with previous reports ^1, 32^, reflecting a primed regenerative state. This response was characterized by reduced immune cell infiltration (Figure 4L, cyan arrows), a predominance of centrally nucleated myofibers (Figure 4L, black arrows), and minimal interstitial matrix deposition (Figure 4L, green arrows). In contrast, AR^(i)sat-/Y^ muscles displayed exacerbated pathological features, including persistent immune infiltration (Figure 4L, cyan arrows), multinucleated fibers with pronounced morphological abnormalities, and disorganized fibers with fusion defects and nuclear mislocalization throughout the sarcoplasm (Figure 4L, pink arrows), together with fat deposition (Figure 4L, red arrows). Immunolabeling for DMD further confirmed the structural heterogeneity and irregularities in AR^(i)sat-/Y^ muscles at 28+7 dpi (Figure 4M). These defects were accompanied a threefold reduction in MuSC numbers in AR^(i)sat-/Y^ mice compared to controls as revealed by quantification PAX7-positive MuSC at 28+7 dpi (Figure 4N).

Overall, these results indicate that AR signaling in MuSC is essential for maintaining the stem cell pool by promoting symmetric divisions and preventing premature differentiation (Figure 4O), thereby ensuring effective regenerative capacity during both primary and subsequent injuries.

### AR controls MuSC fate through early chromatin remodeling

To capture early AR-dependent chromatin events with the potential to drive the scRNA-seq-detected transcriptomic alterations and downstream regenerative defects, we performed CUT&RUN profiling of AR and the active chromatin mark H3K4me2 in young FACS-isolated MuSC at 5 dpi. Consistent with our observations at 7 dpi, CUT&RUN analysis at 5 dpi identified 7,612 AR binding peaks, primarily localized to intronic and intergenic regions (Figure S5A), associated with 7,040 genes. The majority of these AR-bound loci coincided with H3K4me2 enrichment at promoter regions (Figures S5B and S5C). Motif analysis using HOMER revealed that 50.6% of AR peaks contained canonical AR binding motifs (Figure 5A), confirming the specificity of AR-DNA interactions. Functional enrichment analysis of AR target genes demonstrated significant involvement in pathways critical for stem cell homeostasis and tissue remodeling, notably the HIPPO and PI3K-Akt signaling pathways (Figure 5B). Similar to what we described at 7 dpi, gene-centric analysis revealed that AR is mainly recruited within a 100-kb window of more than 50% of differentially expressed genes (Figures 5C and 5D). Representative AR binding loci at key genes involved in these regulatory pathways, including *Clock*, *Eifd2* and *Psma4*, are depicted in Figure 5E. Together, these findings demonstrate that AR engages early with the MuSC chromatin landscape, preferentially at distal regulatory elements enriched in active histone marks, to control the transcriptional programs governing stem cell fate and tissue remodeling.

**Figure 5:**
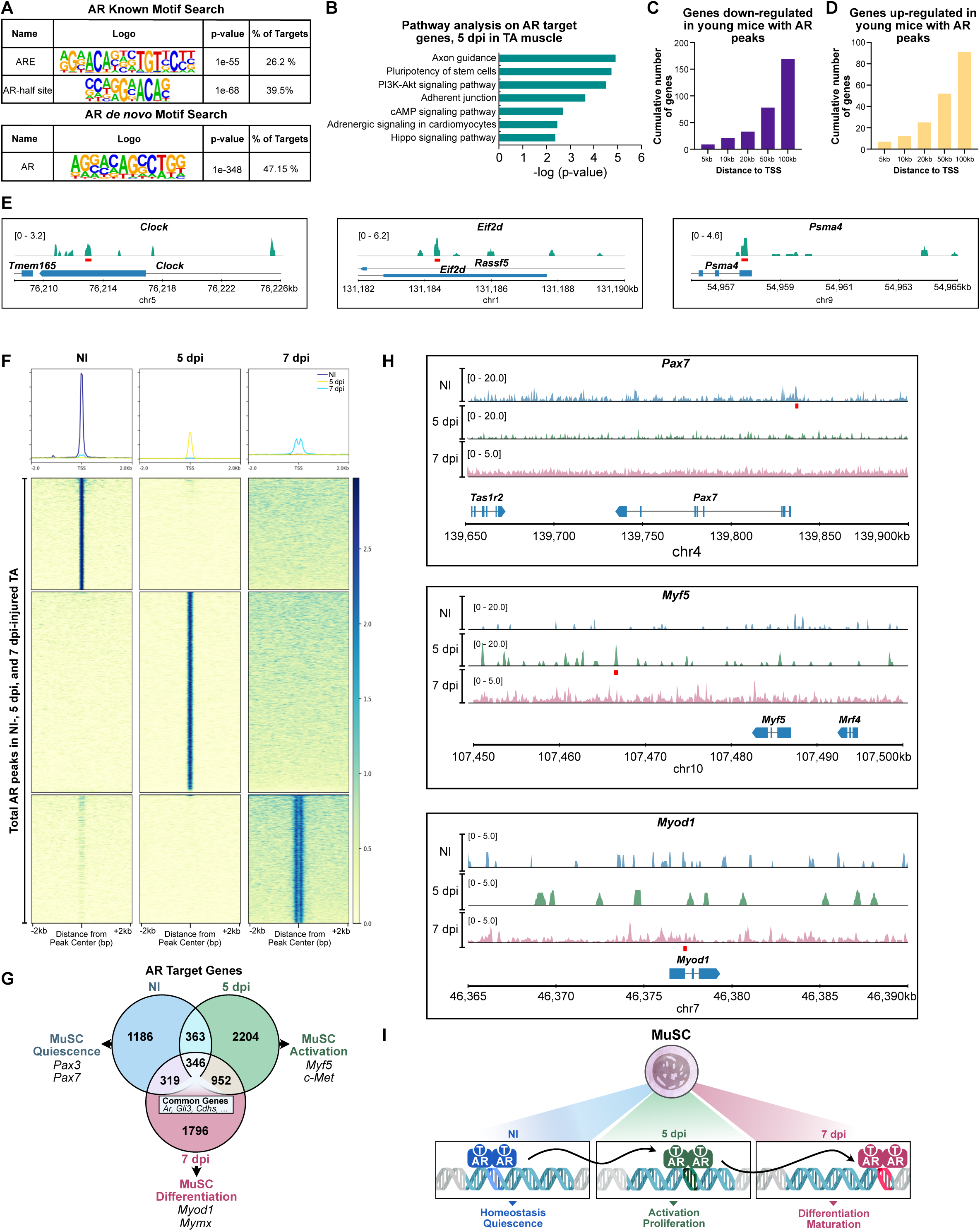
MuSC-AR exhibits time-dependent binding to target genes following injury **(A)** HOMER known and *de novo* motif analyses of AR-bound DNA sequences from 5-day-injured TA of young male mice. **(B)** Pathway analysis performed on AR common target genes identified in MuSC of 5-day-injured TA of young male mice. **(C-D)** Cumulative number of down-regulated genes **(C)**, and up-regulated genes **(D)** in MuSC of 5-day-injured TA of young male mice, with an AR peak located at the indicated distances from their TSS. **(E)** AR localization at the *Clock*, *Eif2d*, and *Psma4* loci. Binding sites are indicated in red. **(F)** Average tag density profiles, and corresponding tag density map of AR in NI, 5-and 7-days injured TA muscles of young male mice, +/− 2 kb from the AR peak center. **(G)** Overlap among genes bound by AR, in NI, 5-and 7-days injured TA muscles of young male mice. **(H)** AR localization in MuSC in NI, 5-and 7-days injured TA muscles of young male mice, at indicated loci. **(I)** Schematic overview of AR-mediated temporal regulation and chromatin binding dynamics in MuSC during regeneration at young age. AR displays stage-specific chromatin occupancy throughout muscle regeneration. In uninjured muscle, MuSC-AR supports homeostasis. Following injury, AR translocates to genomic regions that drive regenerative programs, favoring genes involved in MuSC activation and proliferation at 5 dpi, and shifting toward loci promoting differentiation and maturation by 7 dpi.

### AR dynamically repositions upon injury to govern time-dependent regenerative genomic programs

The transcriptomic discrepancies observed between 5 and 7 dpi led us to question AR positioning on the DNA during muscle repair. Deeptools analysis demonstrated minimal overlap and distinct peak profiles for each state between AR-bound chromatin landscapes 0-(Basal), 5-and 7 dpi (Figures 5F-5H). This temporal specificity was further confirmed by comparative analysis of the genes bound by AR in those specific clusters (Figure 5F). The genes commonly controlled by AR during the regeneration process were mainly implicated in cell-cell adhesion or in the connection with the synapse, essentially *via* cadherins (e.g. *Cdh2*, *Cdh6*, *Cdh7*), and included *Ar* itself, and *Gli3* that is essential for MuSC injury memory ^33^ (Figure 5G). More importantly, in the absence of injury, AR binding was enriched at genomic loci associated with MuSC stemness, including *Pax3*, *Pax7*, and *Kdm1a* (*Lsd1*) (Figures 5G and 5H), while AR-bound landscape profile markedly shifted at 5 dpi, targeting genes implicated in MuSC activation, such as *Myf5* and *Tead1*, and asymmetric division like *Aurkb*, or muscle commitment including *Myh4* and *Myh9* (Figures 5G and 5H). 7 dpi, AR was further reallocated to loci related to myogenic differentiation such as *Myod1*, *Mef2a* and *Mef2d*, and asymmetric division like *Numb* and *Pard3* (Figures 5G and 5H). Representative genes exhibiting AR binding loci in a temporal and stage-specific manner during regeneration such as *Pax7*, *Myf5*, and *Myod1* are depicted in Figure 5H. Collectively, these findings reveal that AR is dynamically repositioned in MuSC following injury, engaging stage-specific genomic loci to orchestrate transcriptional programs that govern activation and lineage commitment via MRFs (Figure 5I). This temporal chromatin remodeling ensures a balanced stem-to-committed transition, safeguarding the MuSC pool for future regenerative demands in young adults.

### Age-related regenerative decline in skeletal muscle is causally linked to diminished androgen signaling

Given the age-related decline in MuSC number and regenerative potential, we hypothesized that diminished androgen signaling may underlie this process. We thus quantified serum testosterone levels in mice at different ages and observed a progressive decrease from ∼25 nmol/L in young mice to a plateau of ∼5 nmol/L in old ones (Figure 6A). This was accompanied by a corresponding decrease in *Ar* expression at both the transcript and protein levels in non-injured (NI) TA muscle (Figures 6B-6D), as well as a decrease in the number of MuSC expressing AR, compared to young mice (Figure S6A). Notably, the transcript levels of the MuSC marker *Pax7* were diminished in NI TA of old control mice (Figure 6E).

**Figure 6:**
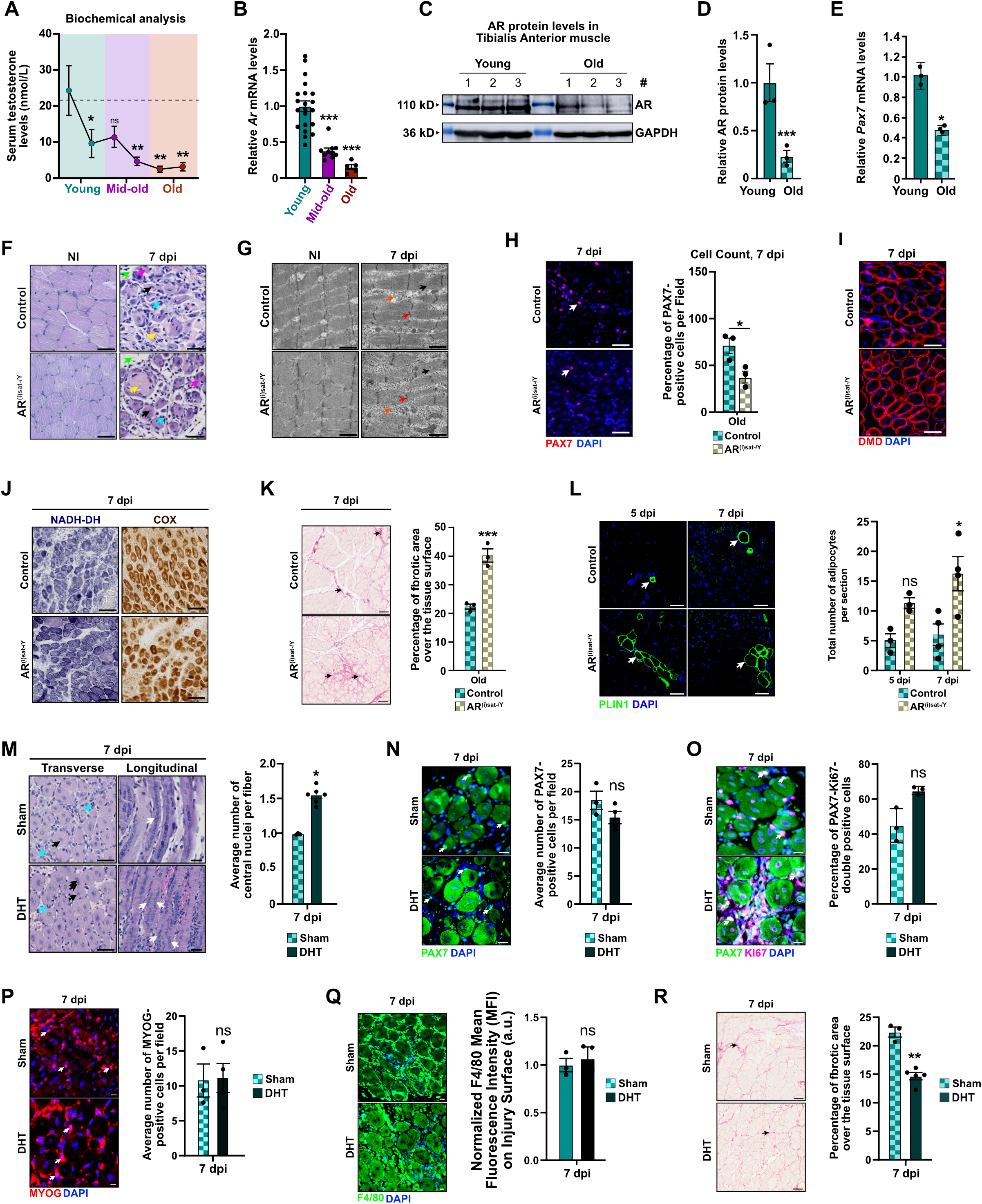
Age-related androgen decline alters skeletal muscle repair machinery **(A)** Serum testosterone levels of control male mice at indicated time-points (young, mid-old and old). Basal level is represented by a dashed line. Data are presented as mean ± SEM. Statistical test used is Ordinary One-way ANOVA relative to 2-month-old mice. ns, non-significant; * = p < 0.05; ** = p < 0.01. **(B)** Relative *Ar* transcript levels in TA muscles of control mice at indicated time-points (young, mid-old and old). Data are presented as mean ± SEM. Statistical test used is Ordinary One-way ANOVA. *** = p < 0.001. **(C-D)** Representative western blot analysis **(C)**, and corresponding quantification of the indicated proteins **(D)** in TA of young and old control male mice. GAPDH was used as a loading control. Data are presented as mean ± SEM. Statistical test used was two-tailed Mann-Whitney test; *** = p < 0.001. (E) Relative *Pax7* transcript levels in TA muscles from young and old control male mice. Data are presented as mean ± SEM. Statistical test used was two-tailed Mann-Whitney test; * = p < 0.05. (F) Representative hematoxylin and eosin (H&E) staining of TA muscles of control and AR^(i)sat-/Y^ old male mice under non-injured (NI) condition, and at 7 dpi. Black arrows indicate centrally nucleated fibers. Blue arrows point to immune infiltration. Yellow arrows denote necrotic fibers. Green arrows show interstitial space. Pink arrows label misshaped regenerating myofibers. Orange arrows mark fibrosis. Scale bars, 75 µm. (G) Ultrastructure analysis of NI and 7-days-injured TA of control and AR^(i)sat-/Y^ old male mice. Black arrows indicate M-line disturbances. Red arrows point to Z-line disruption. Pink arrows depict T-tubules misalignments. Orange arrows denote fibrosis. Green arrows show the A-band placement. Yellow arrows label the I-bands placements. Scale bars, 1 µm. (H) Representative immunofluorescent labeling of PAX7 (in red), and corresponding quantification of the number of PAX7-positive cells in 7-days-injured TA of control and AR^(i)sat-/Y^ old male mice. White arrows point to PAX7-positive cells. Nuclei were stained with DAPI. Data are presented as mean ± SEM. Statistical test used is two-tailed Mann-Whitney test; * = p < 0.05. (I) Representative immunofluorescent detection of DMD (in red), in 7-days-injured TA of control and AR^(i)sat-/Y^ old male mice. Nuclei were stained with DAPI. Scale bars, 100 µm. (J) Representative histochemical staining of NADH dehydrogenase (NADH-DH) and COX activities in 7-days-injured TA of control and AR^(i)sat-/Y^ old male mice. Oxidative and intermediate fibers are darkly and moderately stained, respectively; glycolytic fibers are lightly stained. Scale bars, 200 μm. (K) Representative Sirius Red histochemical staining visualized under brightfield, and corresponding quantification of Sirius Red area, in 7-days-injured TA of control and AR^(i)sat-/Y^ old male mice. Black arrows point to collagen fibers deposition. Scale bars, 250 μm. Data are presented as mean ± SEM. Statistical test used was two-tailed Mann-Whitney test; *** = p < 0.001. (L) Representative immunofluorescent detection of perilipin (PLIN1) (green), and corresponding quantification of total number of adipocytes, in 5-and 7-days-injured TA of control and AR^(i)sat-/Y^ old mice. White arrows point to adipocytes. Nuclei were stained with DAPI. Scale bars, 100 μm. Data are presented as mean ± SEM. Statistical test used is Two-way ANOVA with Tukey correction; ns = non-significant; * = p< 0.05. (M) Representative HE staining, and corresponding quantification of the average number of central nuclei of TA muscles of control Sham-operated or DHT-supplemented old male mice, 7 dpi. Cyan blue arrows point to immune infiltration. Black arrows denote central nuclei. White arrows show central or parallel nuclei alignments. Scale bars, 150 µm for transverse, and 100 µm for longitudinal sections. Data are presented as mean ± SEM. Statistical test used was two-tailed Mann-Whitney test; * = p < 0.05. **(N-Q)** Representative immunofluorescent detections of PAX7 (green) **(N)**, PAX7 (green) and KI67 (magenta) **(O)**, MYOG (red) **(P)**, and F4/80 (green) **(Q)**, and corresponding quantifications per field of injury in TA muscles of control Sham-operated or DHT-supplemented old male mice, 7 dpi. Nuclei were stained with DAPI. Scale bars, 50 µm (N), 20 µm (O), 20 µm (P), and 50 µm (Q). Data are presented as mean ± SEM. Statistical test used was two-tailed Mann-Whitney test; ns, non-significant.

To determine the functional consequences of these baseline deficits, CTX injury was induced in 1-year-old (hereafter old) control and mutant mice, derived from young mice treated with tamoxifen at 9 weeks and subsequently aged under natural conditions (Figure S6B). As expected, and consistent with previous reports ^10, 11^, old male mice exhibited markedly impaired regenerative capacity compared to young littermates (Figures S6C and S1K). To dissect the additional contribution of MuSC-specific AR loss to this decline, we directly compared aged mutants to age-matched controls. Of note, while PAX7-positive MuSC in control mice exhibited faint AR expression, it was entirely absent in AR^(i)sat-/Y^ mutants (Figures S6B and S6D). At 7 dpi, aged AR^(i)sat-/Y^ muscles displayed increased interstitial space, sustained immune infiltration, and misshaped regenerating fibers when compared to (Figure 6F). Ultrastructural analysis confirmed these impairments, showing disrupted sarcomere organization, fragmented Z-lines, and increased extracellular matrix deposition (Figure 6G). These structural impairments were accompanied by a significant decrease in MuSC numbers (Figure 6H), and deformed regenerating fibers (Figures 6I and S6E). In addition, recovery in aged muscles was accompanied by a pronounced shift toward oxidative metabolism (Figure 6J), increased fibrosis (Figure 6K) and adipocyte emergence (Figure 6L), highlighting the pivotal role of AR signaling in safeguarding the structural and functional fidelity of regenerating muscle. Notably, the regenerative phenotype observed in aged control muscles partially recapitulated features seen in young AR^(i)sat-/Y^ mutants, whereas the combination of aging and MuSC-specific AR loss further exacerbated these defects, highlighting the additive impact of hormonal decline and receptor deficiency on regenerative capacity (Figure S6C).

To determine whether the regenerative impairment is causally linked to androgenic insufficiency, we tested whether restoring androgen signaling could reverse aspects of the aged phenotype. We thereby supplemented old mice with 5-α-dihydrotestosterone (DHT) implants. DHT supplementation in old mice led to increased AR protein levels in gastrocnemius skeletal muscle 1 week after implantation (Figures S6F). At 7 dpi, DHT-treated mice exhibited significantly improved muscle regeneration, as evidenced by improved myofiber morphology, and an increased number of nuclei per regenerating fiber (Figure 6M). Although the total number of MuSC was not significantly altered (Figure 6N), we observed a slight increase in proliferating PAX7/KI67-double positive cells (Figure 6O), which likely contributes to the increased number of centrally located nuclei and enhanced myogenic fusion. Importantly, no significant differences were detected in differentiation capacity or immune infiltration, as assessed by MYOG and F4/80 staining, respectively (Figures 6P and 6Q). In addition, fibrosis analysis revealed a marked reduction in fibrotic area, with decreased collagen deposition within the extracellular matrix (Figure 6R). Together, these findings provide direct evidence that age-related androgen deficiency causally contributes to impaired aging skeletal muscle regeneration.

### Androgen signaling orchestrates distinct gene networks in young and old individuals leading to similar phenotypes

To understand how the age-related decline in androgen signaling affects MuSC regenerative programs, we first compared the transcriptomic profiles of old control MuSC to those of young counterparts at 7 dpi. This analysis identified 1,709 differentially expressed genes, including 798 downregulated and 911 upregulated transcripts (Figures 7A and S7A). Pathway enrichment analyses of the downregulated gene set highlighted a significant suppression of pathways related to protein synthesis and peptide/amide metabolism (Figure 7B), suggesting a reduced biosynthetic capacity in aging MuSC. Notably, several genes essential for myogenic differentiation and structural regeneration were significantly downregulated. These included *Myh4* and *Myod1* in addition to genes involved in sarcomere organization and contractile function, such as nebulin (*Neb*), *Tpm3*, and troponin C type 2 (*Tnnc2*) (Figures 7A and 7C). This transcriptional repression is consistent with the defective regenerative phenotype and histological and ultrastructural impairments observed in aged control mice at 7 dpi. Conversely, pathway analyses on the 911 genes upregulated in MuSC from old control mice compared to young control counterparts, revealed that these genes were predominantly associated with metabolic and biosynthetic processes, notably the regulation of RNA, microRNA, nitrogen metabolism, and nucleobase-containing compound metabolic pathways (Figures 7A and 7B), but also oxidative metabolism processes (e.g. *Acadsb*, *Coq8b, Coq10b* and *Pdk4*) (Figures 7A and 7C), collectively indicating a metabolic transition toward oxidative phosphorylation in old MuSC. These molecular findings align with the histological evidence of enhanced oxidative metabolism observed at 7 dpi in old individuals.

**Figure 7:**
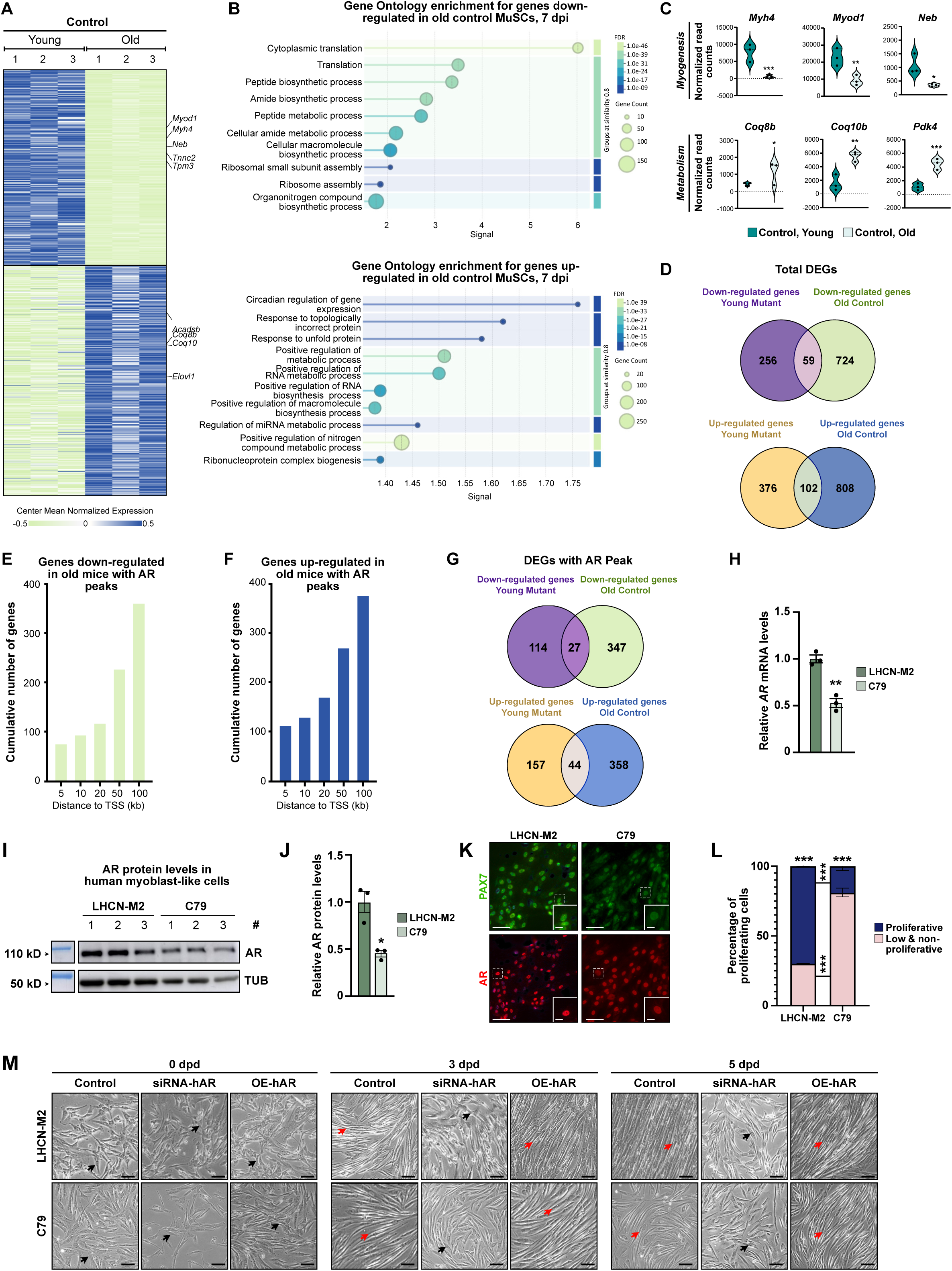
Androgen receptor governs MuSC function in aged muscle through mechanisms that differ from those operating in the young. **(A-B)** Heatmap depicting the mean centered normalized expression of down-and upregulated genes from SMART-seq analysis performed on control MuSC FACS-isolated from 7-day-injured TA of old and young mice **(A)**, and corresponding gene ontology enrichment **(B)**. **(C)** Violin plots representing the normalized read counts of the indicated genes, selected from SMART-seq analysis performed control MuSC FACS-isolated from 7-day-injured TA of old and young mice. Statistical test used is Wald test; * = p < 0.05; ** = p < 0.01; *** = p < 0.001. **(D)** Venn diagrams showing intersected genes among differentially expressed genes in MuSC of control old mice and AR^(i)sat-/Y^ young mice. **(E-F)** Cumulative number of down-regulated genes **(E)**, and up-regulated genes **(F)** in MuSC of 7-day-injured TA of old male mice, with an AR peak located at the indicated distances from their TSS. **(G)** Venn diagrams showing intersected genes with AR peaks among differentially expressed genes in MuSC of control old mice and AR^(i)sat-/Y^ young mice. **(H)** Relative *AR* transcript levels in control LHCN-M2 and C79 myoblasts. Data are presented as mean ± SEM. Statistical test used was two-tailed Mann-Whitney test; ** = p < 0.01. **(I-J)** Representative western blot analysis **(I)**, and corresponding quantification of the indicated proteins **(J)** in control LHCN-M2 and C79 myoblasts. α-TUBULIN (TUB) was used as a loading control. Data are presented as mean ± SEM. Statistical test used was two-tailed Mann-Whitney test; * = p < 0.05. **(K)** Representative immunofluorescent detection of PAX7 (in green) and AR (in red), in LHCN-M2 and C79 cells. Scale bars, 100 μm for the main, 10 μm for magnified inset. **(L)** Quantification of the percentage of high-and low/non-proliferative control LHCN-M2 and C79 myoblasts as determined by CellTrace analysis. Data are presented as mean ± SEM. Statistical test used is Two-way ANOVA with Tukey correction; *** = p< 0.001. **(M)** Representative brightfield images of control (CTR), AR-siRNA transfected (siRNA-hAR) and AR-plasmid transfected (OE-hAR) LHCN-M2 and C79 cells, 0, 3-and 5-days post-differentiation (dpd). Black arrows show fibroblast-like non-differentiated cells. Red arrows label spindle-like differentiated cells. Scale bars, 100 µm.

Since aged control muscles exhibit histological and structural features reminiscent of young AR^(i)sat-/Y^ ones, we next examined age-regulated genes to determine whether these shared changes involve AR-dependent regulatory programs. To this end, we compared age-regulated genes with those deregulated in young mutants and strickingly identified only 59 downregulated and 102 upregulated genes common between young and aged MuSC (Figure 7D). Remarkably, a substantial fraction of differentially expressed genes specific to aged MuSC exhibited AR binding within 100 kb of their TSSs (Figures 7E and 7F), suggesting potential direct AR regulation. To assess the extent to which age-associated transcriptional changes mimics AR loss in youth, we compared AR direct targets identified in young mutant MuSC with age-regulated ones in old controls. This analysis revealed only limited overlap with AR-bound genes (Figure 7G), indicating that the molecular mechanisms underlying AR regulation in young MuSC differ from those driving age-associated transcriptional changes, even though gene ontology analyses revealed similar functional enrichment (Figures S7D and S7F). This limited convergence highlights that, while AR deficiency in MuSC at young age partially recapitulates transcriptional features of aging, the molecular underpinnings differ, reflecting the distinct role of AR during development as opposed to the combined decline of hormone and receptor signaling observed in the elderly. Altogether, these findings support a model in which AR orchestrates regenerative transcriptional programs in MuSC *via* enhancer-driven mechanisms, a regulatory framework that persists with age but becomes functionally altered.

To extend our findings to a human model, we determined AR expression in LHCN-M2 and C79 myoblasts, isolated from young and aged male individuals, respectively. Notably, AR expression was reduced by approximately 50% in C79 compared to LHCN-M2 myoblasts at both the transcript (Figure 7H) and protein levels (Figures 7I and 7J), in addition to the PAX7 levels (Figure 7K). Moreover, LHCN-M2 cells displayed a higher propensity for proliferation (Figures 7L and S7I) and differentiation compared to C79 cells (Figure 7M), mirroring the differences observed between young and aged mice. In agreement, AR knockdown in both cell lines strongly impaired differentiation, as indicated by the conservation of a fibroblast-like morphology, and lack of the characteristic spindle-shaped structure observed in control cells, 5 days post-differentiation (dpd) (Figures 7M, S7J and S7K). Conversely, while AR overexpression did not have flagrant effects on LHCN-M2 cells differentiation, a premature more efficient myogenesis was observed in C79 cells, in the presence of elevated AR levels (Figures 7M, S7L-S7N). Collectively, these findings demonstrate that intact AR signaling, requiring both sufficient ligand and receptor, is essential for efficient myoblast differentiation, and by extension, effective skeletal muscle regeneration.

## DISCUSSION

Maintenance of the MuSC reservoir requires mechanisms that preserve quiescence, while ensuring that repair proceeds with structural and functional fidelity. In young male skeletal muscle, AR expression concentrates in the most quiescent, self-renewing MuSC states, and is largely extinguished as cells commit. Conditional loss of AR in MuSC disrupts dormancy, accelerates cell-cycle entry, and drives symmetric differentiation on the expense of stem cells renewal, leading to attrition of the regenerative pool. Notably, bulk RNA-seq at 7 dpi reveals significant impairments in *Pax7*, *Myf5*, *Myod1*, *Myog*, and *Mymx* expression in AR^(i)sat⁻/Y^ cells, whereas scRNA-seq at 5 dpi does not detect these changes. This discrepancy reflects differences in both regeneration stage and analytical resolution as bulk RNA-seq captures population-wide transcriptional shifts later in repair, whereas scRNA-seq interrogates cell-state-specific signatures at an earlier time point, without implying a biological inconsistency. On this basis, we here identify AR as lineage-intrinsic indicator of MuSC stemness in males, functionally distinct from its canonical anabolic role in myofibers ^16, 25, 26, 34^.

This AR-stemness relationship fits within a broader endocrine framework across the male life course. At young age, androgens promote the establishment of a long-lived quiescent MuSC pool from juvenile precursors through a Notch-dependent pathway ^18^. With aging, declining systemic androgens correlate with reduced MuSC number and impaired repair ^35^, and androgen supplementation restores regenerative capacity in low-androgen settings ^36, 37^. Consistent with this trajectory, our MuSC-specific AR knockout phenotypically recapitulates features of aging-associated stem cell exhaustion, including reduced responsiveness to repeated injuries, suggesting that androgen decline and diminished AR activity converge on shared mechanisms that erode the MuSC reservoir. It is worth noting, however, that transcriptomic differences observed between AR-deficient young MuSC and those from aged controls may also reflect distinct underlying causes, namely, the selective inactivation of AR in otherwise intact young MuSC versus the broader loss of stem cells with age, along with age-related alterations in the muscle microenvironment ^38^. These observations indicate that androgen decline and AR loss exert distinct yet complementary effects, the former reduces stem cell abundance and responsiveness, while the latter disrupts intrinsic programs of quiescence and fate. In aged AR-deficient mutants, these deficits appear to sum, producing an exacerbated phenotype that underscores the dual contribution of hormone and receptor to MuSC maintenance.

We also observe that AR expression in young individuals persists for weeks after injury in MuSC derived from self-renewal, reflecting early symmetric divisions driven by AR that maintain the quiescent stem pool for future regenerative demands. In contrast, differentiating MuSC gradually lose AR, showing that AR not only supports stem cell expansion, but also controls asymmetric divisions to produce committed cells for repair. Without AR, MuSC divisions favor committed daughters over self-renewal, depleting the reserve and impairing regeneration after repeated injury. This AR-dependent balance between renewal and differentiation in male MuSC has not been shown *in vivo* before. Earlier studies using female-derived C2C12 myoblasts showed DHT enhances proliferation, but these cells lack true quiescence and do not represent adult male muscle stem cells ^20, 42, 43^. In contrast, our findings reveal that physiological androgen levels *via* AR in male MuSC dynamically regulates their fate during regeneration.

MuSC-AR influences more than lineage fate in skeletal muscle. It promotes proliferation and differentiation in myogenic cells ^44, 45^, and its loss in our mouse model leads to fusion defects, fiber orientation defaults, and impaired ultrastructure, reflecting not only reduced progenitor output, but also a disrupted regenerative program in these progenitors. This contradicts earlier claims that MuSC-AR is dispensable for regeneration (Sakai et al., 2020), likely because their model did not disrupt AR in the early, true stem MuSC population. Our data clearly establish MuSC-intrinsic AR as a crucial regulator of androgen-driven muscle repair in males. This signaling axis also shapes metabolic identity. While myofiber-AR supports glycolytic metabolism, its absence in MuSC at young age shifts regenerated fibers toward oxidative pathways, which was recapitulated in aged individuals. Seahorse assays and histology confirm this shift, consistent with findings in AR-deficient myofibers ^25, 46^, AR-related disorders ^47^, and aging muscle ^48, 49^. Notably, myofiber-AR loss is linked to increased β-oxidation gene expression and reduced Complex IV components ^25, 50^, though this latter feature was absent in our model, possibly due to reactive oxygen species accumulation impairing electron transport and causing mitochondrial stress ^51^.

At the tissue level, MuSC-AR deletion in young individuals reveals increased inflammation, fibrosis, and adipocyte accumulation at the site of injury. Similar disturbances occur in androgen-deprived states, where impaired inflammatory resolution and accelerated fibro-adipogenic drift are observed ^52, 53^. These findings show that MuSC are not passive recipients, but active transmitters of androgen-sensitive signals shaping their microenvironment. This supports a bidirectional model in which AR in MuSC governs intrinsic fate decisions while orchestrating stromal and immune crosstalk during muscle repair. While our data indicate that MuSC-AR influences macrophages and FAPs, these effects are likely indirect. CellChat analysis did not reveal significant differences in secreted signaling molecules, and *in vitro* co-culture experiments were precluded by the poor adherence of FACS-isolated AR^(i)sat⁻/Y^ MuSC. Therefore, MuSC-AR appears to shape the regenerative niche predominantly through indirect intercellular mechanisms. Importantly, in AR-deficient muscles, macrophages prematurely express anti-inflammatory markers, yet the total inflammatory burden persists longer, reflecting an uncoupling between phenotypic transition and effective clearance. This aberrant trajectory likely contributes to impaired regeneration and highlights the nuanced role of MuSC-AR in coordinating the immune niche. Our CellChat analysis further suggests that AR-deficient MuSC perturb intercellular communication with macrophages and FAPs, although the precise signaling mediators remain to be defined. Identifying these factors represents a compelling avenue for future work, as it would illuminate the mechanisms by which MuSC orchestrate niche remodeling and coordinate regeneration.

To coordinate these diverse, stage-specific functions during muscle regeneration, we propose a dynamic chromatin occupancy model in which AR sequentially engages distinct transcriptional programs as MuSC progress from quiescence through activation to commitment. Such nuclear receptor mobility and partner switching are well documented in other systems ^54–57^. Emerging evidence in skeletal muscle shows AR collaborates with SMAD signaling to enhance BMP-driven transcriptional programs that regulate muscle mass and quality ^58^. In aging and pathological expansions of the AR-polyglutamine tract, altered chromatin binding and disrupted co-factor interactions have been observed ^59, 60^, suggesting that reduced AR availability with age may limit its genomic sampling and assembly of stage-specific complexes necessary to maintain stemness, and sequentially enable differentiation.

In sum, MuSC-AR preserves quiescence, sustains the stem cell pool, shapes glycolytic identity, and orchestrates niche remodeling. Its decline leads to reserve loss, metabolic drift, and regenerative failure (Figure S7O), highlighting MuSC-AR as a therapeutic target in aging and disease.

## METHODS

### Mouse Studies

#### Housing

Mice were maintained on a C57BL/6 background and housed in a temperature-and humidity-controlled facility under a 12-hour light/dark cycle, with *ad libitum* access to standard rodent chow (2800 kcal/kg, Usine d’Alimentation Rationelle, Villemoisson-sur-Orge, France) and water. Breeding and maintenance followed institutional guidelines, and all experiments were conducted in an accredited facility in compliance with French and EU regulations on laboratory animal use (Directive 2010/63/EU). Procedures were approved by the local ethical committee (Com’Eth, Strasbourg, France) and by the French Ministry of Research (MESR). Mice were euthanized *via* cervical dislocation, and muscles were harvested for histological and molecular analysis.

#### Generation of AR^(i)sat−/Y^ mutant mice

For satellite cell-specific androgen receptor (AR) ablation, AR^L2/L2^ female mice, carrying LoxP sites flanking chromosome X exon 1 encoding the AR N-terminal domain ^61^, were intercrossed with *Pax7*-CreER^T2^ male mice ^62^ to generate *Pax7*-CreER^T2^/AR^L2/Y^ mice, which express CreER^T2^ recombinase selectively in satellite cells. To induce recombination, 9-week-old AR^L2/Y^ control males and their *Pax7*-CreER^T2^/AR^L2/Y^ pre-mutant littermates received intraperitoneal tamoxifen injections (1 mg/mouse/day) for five consecutive days, generating control and AR^(i)sat−/Y^ mutant mice. Alternatively, mice were intercrossed with a *Rosa26*-YFP reporter line ^63^ to generate *Pax7*-CreER^T2^/AR^L2/Y^/*Rosa26*-YFP^Tg/0^ mice with one YFP allele, allowing selective labeling of satellite cells and lineage tracing of their progeny. *Pax7*-CreER^T2^/Rosa26-YFP^Tg/0^ mice lacking the floxed AR allele were used as additional controls. Mice were analyzed at 3 months of age (young adults) or at 1 year of age (aged cohort). Primer sequences required for genotyping PCR are listed in Table S4.

#### Mouse Treatments

For cardiotoxin-induced muscle injury model, 3 months young adults or at 1-year aged mice were anesthetized with 2.5% isoflurane using a SomnoSuite device. To induce muscle damage, 20 μl of 25 μM cardiotoxin (CTX) (Latoxan, Valence, France, L8102), dissolved in phosphate-buffered saline (PBS) were injected into the tibialis anterior (TA) muscle.

For androgen supplementation, 5-alpha-dehydrotestosterone (DHT) implants were inserted subcutaneously in the intrascapular region of 21-month-old wild-type male mice, as described ^64^. Muscle injury was induced by CTX injection three weeks post-implantation to avoid off-target effects caused by DHT-burst.

#### Testosterone serum levels measurement

Blood was collected from mice in 50 mM EDTA tubes (Sigma-Aldrich), and plasma was isolated by centrifugation at 3,000 g for 15 minutes at 4°C. The supernatant was collected, and serum testosterone levels were quantified using the Testosterone Rat/Mouse ELISA Kit (LDN, AR E-8000R), following the manufacturer’s instructions.

#### Contractile Measurements

*In situ* isometric TA muscle force in response to nerve stimulation was measured as described ^26^. In brief, specific maximal tetanic force (mN/mm^2^) was determined by dividing the measured force by the estimated CSA of the muscle. Assuming a cylindrical muscle shape with a density of 1.06 mg/mm^3^, the CSA was calculated as the volume of the muscle divided by its (muscle) length. The optimal muscle length (L0) was defined as the length at which maximal tetanic force was observed. All data collected from the muscle lever system were recorded and analyzed using a microcomputer with the PowerLab/4SP system (AD Instruments).

### Histological analyses

#### Muscle histology

Muscles were either snap-frozen in liquid nitrogen-cooled isopentane for cryosections, or fixed in 4% PFA overnight at 4 °C then washed in PBS and 70% ethanol and embedded in paraffin. 10 μm thick cryosections were prepared at −20 °C using a 2800 Frigocut cryostat (Leica, St Jouarre, France) and kept at −20 °C until further analysis. 5 μm thick paraffin sections were cut at room temperature (RT) using a microtome (Leica RM, 2145, 0737/09.1998) and stored at 4 °C until further analysis.

Hematoxylin and eosin, and Masson’s Trichrome staining were performed on paraffin sections as described, while NADH and COX staining were assessed on cryo-sections as described ^25, 63, 65^.

#### Quantification of Collagen Deposition

Collagen content assessed by Sirius Red staining (Picro-Sirius Red Stain, Pint, StatLab, STPSRPT) was imaged under a bright-field microscope, or under polarized light using the Axioscan 7 (Carl Zeiss) slide scanner. Birefringent collagen fibers were visualized based on their differential light refraction properties, allowing for the selective detection of collagen deposition. Quantification was performed by analyzing the polarized signal intensity and distribution across the tissue using standardized image analysis parameters established by the in-house anatomic pathology platform.

#### Immunofluorescence staining

Paraffin and cryo-sections were processed using distinct protocols tailored to antibody requirements. Paraffin sections underwent deparaffinization, and antigen retrieval was done under controlled temperature and pressure conditions for 20 min in either tris-hydroxymethyl-aminomethane (Tris buffer) (pH=9.0) or in Signal Stain ® Citrate Unmasking Solution (Cell Signalling 14746S). In contrast, frozen sections were fixed for 10 min in 4% PFA at RT, and permeabilized with PBST for 15 min at RT. Afterwards, paraffin or frozen sections were incubated overnight at 4 °C with the primary antibodies diluted in PBST directed against AR (Rabbit, Abcam, ab108341, 1:500), PAX7 (Rabbit; Thermo Fisher, PA1-117, 1:400, or Mouse; DSHB, AB_528428, 1:50), MYOG (Abcam, ab1835, 1:400), ACTN1 (Mouse,

Abcam, ab9465, 1:400), DMD (Mouse, IGBMC, 1Dystro1D2, 1:1000 or Rabbit sigma, D8168, 1:400), KI67 (Rat, R&D systems, 14-5698-82, 1:500), LAMA2 (Rat, Sigma Aldrich, L0663-.1ML, 1:500), eMyHC

(Mouse, DSHB, F1.652, 1:20), F4/80 (Rabbit, Cell Signaling, 70076, 1:400) and PLIN1 (Rabbit, Abcam, Ab3526. 1:200). Mouse (Santa Cruz Biotechnology, E3117, 1:500), rat (Vector Laboratories, BA-4000, 1:500) or rabbit (PeproTech, 500-P000-500UG, 1:500). IgGs were used as controls. Sections were rinsed with wash buffer, and incubated with appropriate Alexa-Fluor-conjugated secondary antibodies with different fluorescent wavelengths (488, 555 or 647 nm, 1:500, Invitrogen: a32732, a32723, a21206, a21424, a48265, a32733, a32728) in permeabilization buffer for 45 min at RT. After wash, slides were mounted with Fluoromount™ Aqueous Mounting medium containing DAPI (Invitrogen, 00-4959-52).

#### Fiber cross-sectional area measurements

Muscle cross-sections were stained with DMD (Abcam, ab15277, 1:500) mark the sarcolemma surrounding each fiber. CSAs were quantified using QuPath (open-source image analysis software) integrated with a customized Groovy macro (available via GitLab). In brief, individual fibers were identified based on the intensity and continuity of the stained sarcolemma surrounding each fiber by segmentation. Areas were measured after background subtraction and automated thresholding.

#### Quantification of F4/80 and CD163 mean fluorescence intensity (MFI)

Fluorescence images of injured TA muscles immunostained for F4/80 and CD163 were analyzed using QuPath software (QuPath-0.5.0). Regions of interest (ROIs) corresponding to the injured areas were annotated manually, and a pixel classifier was trained on these annotations to categorize pixels as background/negative, positive signal, or other non-positive signals. Fluorescence intensity thresholds were set manually for each marker to ensure accurate detection of positive staining. Mean fluorescence intensity within the annotated ROIs was then measured and exported for further analysis.

#### Muscle ultrastructural analysis

1 mm² TA muscle samples were fixed overnight at 4°C by immersion in a mix of 2.5% glutaraldehyde and 2.5% paraformaldehyde diluted in 0.1 M cacodylate buffer (pH 7.4), washed in the same buffer for 30 min, and stored at 4°C. Post-fixation was performed with 1% osmium tetroxide in 0.1 M cacodylate buffer for 1 h at 4°C, followed by dehydration through a graded ethanol series (50%, 70%, 90%, and 100%) and propylene oxide (30 min each). Samples were oriented longitudinally or transversally and embedded in Epon 812. 70 nm ultrathin sections were obtained and contrasted with uranyl acetate and lead citrate. Imaging was performed at 70 kV using a Morgagni 268D electron microscope, with digital acquisition via a Mega View III camera (Soft Imaging System).

### Cell culture analyses

#### Cell culture and treatments

LHCN-M2 and C79 cells were cultured in Dulbecco’s Modified Eagle Medium (DMEM 4,5 g/L of glucose, Invitrogen 4165-062)/Medium 199 (Invitrogen 22340-020) (4:1), supplemented with 20% fetal calf serum (FCS), 25 µg/mL fetuin, 5 µg/mL human insulin, 40 µg/mL gentamycin and 0.5 ng/mL basic fibroblast growth factor (bFGF) along with 5 ng/mL human epidermal growth factor (hEGF) and 0.2 µg/mL dexamethasone ^66^. Cells were maintained at 37°C in a humidified atmosphere with 5% CO₂. Cells were split at 70-80% confluence, and the medium was replaced every 48 hours.

To induce differentiation, LHCN-M2 and C79 cells were seeded at 90% confluence and switched at confluence to differentiation medium (DMEM supplemented with 2% horse serum and 40 µg/mL gentamycin) for up to 5 days, with medium changes every 48 hours.

For transfection, human AR siRNA (5’-GGGCACTTCGACCATTTCTGA-3’), or scramble siRNA (5’-AAGCTTCATAAGGCGCATAGC-3’), were introduced using Typical RNAiMAX Transfection kit (ThermoFischer, 13778075), following the manufacturer’s protocol. Cells were seeded in six-well plate and transfected with 50 nM siRNA diluted in Opti-MEM (Gibco, USA) at 50-60% confluence for 24h, after which the medium was replaced with fresh differentiation or growth medium. Cells were subsequently treated with 100 nM of DHT (Sigma, A8380-1G) diluted in ethanol 24h after transfections.

For AR overexpression, MuSC were transfected with the pEGFP-AR^WT^ plasmid using Lipofectamine 2000 DNA Transfection kit (ThermoFisher, 11668027), according to the manufacturer’s instructions. Cells were seeded in six-well plate and transfected with 14 µg pEGFP-AR^WT^ plasmid diluted in Opti-MEM (Gibco, USA) at 50-60% confluence for 24h, after which the medium was replaced with fresh differentiation or growth medium. Cells were subsequently treated with 100 nM of DHT (Sigma, A8380-1G) diluted in ethanol 24h after transfections.

#### Cell Proliferation Analysis

Cell proliferation was assessed using the CellTrace™ CFSE (Carboxyfluorescein Succinimidyl Ester) Assay (Thermo Fisher Scientific, USA, C34557) as described ^67^. Briefly, LHCN-M2 and C79 cells were incubated with 5 µM CFSE at 37°C for 20 min in a humidified atmosphere with 5% CO₂. The labeling reaction was quenched by adding an equal volume of complete growth medium, followed by two washes with PBS to remove excess dye. Labeled cells were kept in appropriate culture conditions and collected at 24-, 48-, and 72-hours post-labeling for flow cytometry analysis. CFSE fluorescence intensity was measured using a flow cytometer (BD LSRFortessa™, BD Biosciences, USA) with excitation at 488 nm and emission detection at 530 nm. Data were analyzed using FlowJo software (BD Biosciences, USA), and cell proliferation was determined by monitoring the progressive dilution of CFSE fluorescence across multiple cell generations. To identify non-and slow-proliferating cells, we defined the 30% of cells exhibiting the highest CellTrace Violet signal in control as the reference population, and calculated the proportion of cells with equal or greater signal intensity.

#### Immunocytochemistry

Immunocytofluorescence assay was performed by incubating fixed LHCN-M2 and C79 cells with AR (Rabbit, Abcam, ab108341, 1:500) and PAX7 (Mouse; DSHB, AB_528428, 1:50) antibodies. Observations were made under a Leica confocal microscope.

#### Seahorse analysis of LHCN-M2 myoblasts

Mitochondrial respiration was assessed using the Seahorse XF Cell Mito Stress Test Kit (Agilent Technologies, Part Number: 103015-100) according to the manufacturer’s protocol. Briefly, ∼1800 LHCN-M2 myoblasts were seeded in Seahorse XF cell culture microplates (Agilent), and treated with either vehicle or 100 nM flutamide (Abcam, ab141234) for 1 hour. The assay was performed on a Seahorse XFe96 Analyzer (Agilent), following the sequential injection of oligomycin (1 μM), FCCP (2.5 μM), and rotenone/antimycin A (1 μM). Basal respiration, ATP-linked respiration, maximal respiration, and spare respiratory capacity were calculated using Wave software (Agilent).

### Flow cytometry analysis

#### Muscle cell populations composition

Muscle stromal vascular fraction was isolated as described ^68^. Briefly, hind limb muscles were dissected, minced into small fragments, and enzymatically digested with 1.25 U/mL Dispase II (Stemcell Technologies, Cat: 07913) and 500 µg/mL Collagenase Type I (Thermo Fisher Scientific, 17018029). The resulting cell suspension was sequentially filtered through 100 µm, 70 µm, and 40 µm cell strainers (Corning Life Sciences) to remove debris. Cells isolated from digested muscle tissue were stained with CD45 Alexa eFluor700 (eBioscience, 56-0451-82, 1:100), CD31 PE (eBioscience, 12-0311-82, 1:100), SCA-1 PE-Cy7 (eBioscience, 25-5981-82, 1:100), and Integrin-α7 (ITGΑ7) FITC (MBL, K0046-4, 1:50). Cells were analyzed based on forward scatter (FSC) and side scatter (SSC) parameters to exclude debris and doublets. CD45-and CD31-positive cells were quantified to assess immune and endothelial cell populations, respectively. After removing CD45/CD31-expressing cells, SCA-1-positive cells were gated to identify fibro-adipogenic progenitor cells (FAPs). Finally, SCA-1-negative, ITGΑ7-positive cells were classified as satellite cells. Fluorescence intensity was measured using a flow cytometer (BD FACSymphony A1™, BD Biosciences, USA). Analysis were conducted using FlowJO software as described ^68^.

#### Muscle immune cell populations composition

Cells isolated from digested muscle tissue were stained with CD45 Alexa eFluor700 (eBioscience, 56-0451-82, 1:100), CD11b PerCP-Cy5.5 (BioLegend, 101228, 1:100), Ly-6G-GR-1 FITC 488 (BioLegend, 108417, 1:100), Ly-6C PE-CF594 (BD Horizon™, AB_2737749, 1:100), and F4/80 APC eFluor 780 (eBioscience™, 47-4801-82, 1:100). Cells were analyzed based on forward scatter (FSC) and side scatter (SSC) parameters to exclude debris and doublets.

Data analysis was performed by sequential gating on single, live CD45⁺ leukocytes, followed by identification of myeloid cells as CD11b⁺, neutrophils as CD11b⁺Ly6G⁺, monocytes as CD11b⁺Ly6G⁻Ly6C⁺, and macrophages as CD11b⁺Ly6G^-^ Ly6C^-^F4/80⁺. Fluorescence intensity was measured using a flow cytometer (BD FACSymphony A1™, BD Biosciences, USA). Analyses were conducted using FlowJO software.

### Fluorescence-Activated Cell Sorting (FACS) of Satellite Cells

After stromal vascular fraction isolation, mononucleated cells were pelleted by centrifugation at 400g for 5 min, resuspended in FACS buffer (DMEM without phenol red, 4.5 g/L glucose, 2% BSA, 2 mM EDTA), and labeled with fluorescence-conjugated antibodies. Cells were stained with CD45 PE (eBioscience, 12-0451-83, 1:100), CD11b PE-Cy7 (eBioscience, 25-0112-82, 1:100), CD31 PE (eBioscience, 12-0311-82, 1:100), TER119 PE (BD Pharmingen^TM^, 553673, 1:100), ITGΑ7 FITC (MBL, K0046-4, 1:50), CXCR4 APC (eBioscience, 17-9991-82, 1:100), and CD34 eFluor405 (eBioscience, 48-0341-82, 1:33). Cells were gated based on FSC and SSC parameters to exclude cellular debris and doublets. Leukocytes and endothelial cells were removed by gating out CD45-, CD11b-, CD31-, and TER119-positive populations. The remaining fraction was further gated for CD34 and ITGΑ7 expression, and double-positive cells were selected based on CXCR4 expression. Fluorescence intensity and sorting were conducted using a FACS sorter (BD FACSAria™, BD Biosciences, USA). Sorted satellite cells were subsequently processed for RNA extraction or CUT&RUN analyses.

### CUT&RUN Assay and Bioinformatics Analysis

CUT&RUN was performed as described ^68, 69^. Briefly, FACS-isolated satellite cells were centrifuged at 500 g for 10 min at 4°C, and the resulting cell pellet was incubated with nuclear extraction buffer (20mM HEPES-KOH pH = 7.9, 10 mM KCl, 0.5 mM Spermidine, 0.1% Triton X-100, 20% Glycerol and protease inhibitor cocktail) for 20 min on ice. Isolated nuclei were then washed and incubated with Concanavalin A-coated beads (BioMag Plus Concanavalin A, Cat: 86057-3) for 10 min at 4°C. Following additional washes, the nuclei-bead complexes were incubated overnight at 4°C with gentle agitation in the presence of primary antibodies targeting AR (Abcam, ab108341, 1:100) and H3K4me2 (Active Motif, 39141, 1:100), and a Rabbit IgG (PeproTech, 500-P000-500UG, 1:100). Nuclei-bead complexes were precipitated and incubated with protein A-micrococcal nuclease (pA-MN, IGBMC) for 1 h at 4°C under constant agitation. Enzymatic cleavage was initiated by adding 3 µL of 100 mM CaCl₂, followed by 30 min of incubation on ice. The reaction was terminated at 37°C for 20 min using stop buffer (200 mM NaCl, 20 mM EDTA, 4 mM EGTA, 50 µg/mL RNase A, 40 µg/mL glycogen, 25 pg/mL yeast spike-in DNA). DNA was extracted from immunoprecipitated samples for sequencing.

Libraries were generated from immunoprecipitated DNA as described ^68^, and sequenced on an Illumina HiSeq 2000 platform. Reads overlapping ENCODE blacklist regions (V2) were removed, and deduplicated unique reads were conserved for further analysis. Reads were mapped to the mm10 reference genome using Bowtie2 (v2.3.4.3) ^70^. To generate genome-wide intensity profiles, uniquely mapped reads were processed as follows: Bigwig files were generated using bamCoverage (deeptools 3.3.0) ^71^, with normalization by RPKM (--normalizeUsing RPKM --binSize 20). Raw bedgraph files were produced with genomeCoverageBed (bedtools v2.26.0) ^72^. Peak calling was performed using SEACR ^73^ and MACS2 ^74^ with IgG-immunoprecipitated samples as controls (-keepall), and peaks with an FDR at least less than 0.05 were retained for further analysis. The IGV genome browser (http://software.broadinstitute.org/software/igv/) was used for genome-wide intensity visualization. Peaks were annotated, and motif analysis was conducted using HOMER. Genomic feature distribution (promoters/TSS, 5’ UTRs, exons, introns, 3’ UTRs, transcription termination sites, and intergenic regions) was determined based on RefSeq and HOMER annotations. For bioinformatics and clustering analysis, Venn diagrams were generated using Venny (2.1.0) (https://bioinfogp.cnb.csic.es/tools/venny/).

Clustering analyses were performed using seqMINER ^75^ and deeptools (-computeMatrix-skip0; - plotHeatmap) with K-Means linear normalization. Genomic intersections and sequence extractions were carried out using bedtools. GetFastaBed was used to extract FASTA sequences from BED files. Intersect Interval and Multiple Intersect were applied to identify overlapping genomic locations. WindowBed was utilized for gene-centric peak distribution analyses. When not specified, all bioinformatics analyses were performed using default parameters, ensuring robust and reproducible results.

### RNA extraction and analyses

Total RNA from muscle tissue, LHCN-M2 and C79 cell lines was extracted using TRIzol reagent (TRIREAGENT, Molecular Research Center, TR-118, 513-841-0900) as described ^76^. RNA concentration and purity were assessed spectrophotometrically using a NanoDrop system (Thermo Fisher Scientific). For RNA isolation from FACS-isolated MuSC, the RNeasy Plus Micro Kit (Qiagen, 74034) was employed according to the manufacturer’s instructions.

For RT-qPCR experiments, 2 µg of total RNA underwent reverse transcription using SuperScript IV (Life Technologies) with oligo(dT) primers, according to the supplier’s protocol. cDNA was diluted hundred times and quantitative PCR (qPCR) was performed with a Lightcycler 480 II (Roche) using the SYBR® Green PCR kit (Roche), in accordance with the recommended protocol (2 µl cDNA, 4.8 µl H_2_O, 5 µl Syber Green 2x mix and 0.2 µl of 100 µM primer mix). The primer sequences for mouse and human are detailed in Tables S5 and S6, respectively. Housekeeping genes *Tbp* (for mouse), and *HSP90AB1* (for human) were used as internal controls. Data were analyzed using ΔΔCt ^77^ method.

For RNA sequencing (RNA-seq) and SMART-seq experiments, RNA integrity was assessed using a Bioanalyzer. Library preparation was performed at the GenomEast platform from IGBMC.

For RNA-seq, cDNA library was prepared, and sequenced by the standard Illumina protocol (HiSeq 4000, single-end, 50 bp), following the manufacturer’s instructions. Image analysis and base calling were performed using RTA 2.7.7 and bcl2fastq 2.17.1.14. Adapter dimer reads were removed using DimerRemover. FastQC 0.11.2 (http://www.bioinformatics.babraham.ac.uk/projects/fastqc/) was used to evaluate the quality of the sequencing. Reads were mapped to the mouse mm10 genome (NCBI Build 38) using STAR v2.7.10b ^78^. Only uniquely aligned reads were retained for further analyses.

Quantification of gene expression was performed using HTSeq 0.11.0 ^79^. For comparison among datasets, the transcripts with more than 100 raw reads were considered.

For SMART-seq, full length cDNA was generated from 250 pg to 2 ng of total RNA using SMART-Seq® mRNA Kit (Takara Bio Europe, Saint Germain en Laye, France), according to manufacturer’s instructions, with 12 cycles of PCR for cDNA amplification by Seq-Amp polymerase. Six hundred pg of pre-amplified cDNA were used as input for Tn5 transposon tagmentation followed by 12 cycles of library amplification using Nextera XT DNA Library Preparation Kit and IDT for Illumina DNA/RNA UD Indexes, Tagmentation (Illumina, San Diego, USA). Following purification with SPRIselect beads (Beckman-Coulter, Villepinte, France), the size and concentration of libraries were assessed using Bioanalyzer 2100 system (Agilent technologies, Les Ulis, France). Libraries were sequenced on an Illumina NextSeq 2000-X sequencer as paired-end 50 base reads. Image analysis and base calling were performed using RTA version 2.7.7 and BCL Convert version 3.8.4. Adapter dimer reads were removed with DimerRemover (-a AGATCGGAAGAGCACACGTCTGAACTCCAGTCAC), and the quality of sequencing data was evaluated using FastQC 0.11.5. Reads were aligned to the mouse mm39 genome using STAR v2.7.10b, retaining only uniquely mapped reads for downstream analyses. Gene expression was quantified using STAR, and for dataset comparisons, only transcripts with more than 50 raw reads were considered.

For RNA sequencing (RNA-seq) and SMART-seq experiments, differentially expressed genes (DEGs) were identified using the Bioconductor libraries DESeq2 ^80^ with a p < 0.05. Identified DEGs were subjected to functional analysis using STRING 12.0 ^81^ and to pathway analysis using WebGestalt ^82^ with the Over-Representation Analysis (ORA) method, applying a false discovery rate (FDR) < 0.05 as the significance threshold. Heatmaps were generated by centering and normalizing expression values using Cluster 3.0 ^83^, with subsequent visualization Morpheus (https://software.broadinstitute.org/morpheus/). Genes were clustered using K-Mean with gene tree construction, Pearson correlation, and average linkage as clustering parameters ^84^.

### Single-cell RNA sequencing

Single-cell RNA sequencing (scRNA-seq) experiments were performed on TA muscle cells, isolated from 3 control (AR^L2/Y^) and 3 AR^(i)sat-/Y^/YFP^Tg/0^ injured TA muscles at 5 dpi. TA muscles were dissociated into single cells, as described above. Cells from each genotype group were pooled and stained with DAPI, and DAPI-negative living cells were isolated using a BD FACS Aria™ Fusion flow cytometer. To assess cell viability and yield, a Trypan blue exclusion assay was performed using a Neubauer Chamber. Only samples with a viability greater than 95% were processed using a Chromium Controller (10X Genomics, Leiden, The Netherlands). 16,500 total cells were loaded per well to yield approximately 10,000 captured cells into nanoliter-scale Gel Beads-in-Emulsion (GEMs). Single cell 3 prime mRNA seq library were generated using Chromium Next GEM Single Cell 3 prime Reagent Kits v3.1 (10X Genomics). Libraries were quantified and controlled for their quality using Bioanalyzer 2100 (Agilent Technologies, Santa Clara CA), and further sequenced on an Illumina NextSeq 550 sequencer as paired-end 28 + 100 base reads. Image analysis and base calling were performed using RTA version 2.7.7 ^85^ and Cell Ranger version 3.0.2 ^86^.

Data were processed using the Cell Ranger 7.0.0 count pipeline (10x Genomics) for read alignment, barcode and unique molecular identifier (UMI) filtering, and gene quantification. A custom reference genome was generated using the mkref function with the GRCm39 assembly of the *Mus musculus* genome and Ensembl release 107 annotations. The output gene-barcode matrix was imported into RStudio (version 2025.05.1) using the Read10X function from the Seurat package (version 5), yielding a sparse matrix of UMI counts per gene per cell. Downstream analyses were conducted in RStudio following standard Seurat v5 workflows ^87^, including quality control, normalization, dimensionality reduction, clustering, and differential gene expression. For quality control, cells with <10% mitochondrial content, <50% ribosomal gene expression, and total RNA features between 200 and 6000 were retained. Principal Component Analysis (PCA, dim=16) was used for dimensionality reduction, and clustering was performed using a resolution parameter of 0.6. Differential gene expression and further downstream analyses were conducted. Satellite cell reclustering was performed with dimension reduction of 14 and cluster resolution of 0.3.

Cell-cell communication analysis was conducted using the CellChat R package within the RStudio environment ^88^. Normalized scRNA-seq data were used, and overexpressed ligands and receptors were identified to infer intercellular signaling networks. Communication probabilities were computed using the trimean method, with a minimum gene expression threshold of 0.1, and genes were considered only if expressed in more than 10% of cells in the described groups. The CellChatDB ligand-receptor interaction database was used as the reference. The analysis pipeline included identification of major signaling inputs and outputs, classification of signaling roles (incoming, outgoing, mediator, and influencer), and visualization via circle plots and hierarchical clustering. Inter-condition comparisons were performed using the compareInteractions and rankNet functions. All analyses were performed using default CellChat parameters, including 500 permutations for significance testing and standard visualization settings, unless otherwise specified. Results were visualized and further interpreted using built-in and custom R scripts. Cell cycle bioinformatic analyses were performed using Scanpy function (scanpy.tl.score_genes_cell_cycle), with marker genes extracted from scRNA-seq of mouse embryonic stem cells generated for DeepCycle as described ^87, 89^.

### Protein analysis

TA muscles were mechanically ground in RIPA buffer [50 mM Tris (pH 7.5), 1% NP-40, 0.5% sodium deoxycholate, 0.1% SDS, 150 mM NaCl, 5 mM EDTA, and a protease inhibitor cocktail (45 mg/ml; Roche, Cat. No. 11 873 580 001)] at 4 °C. Alternatively, adherent cells were harvested in RIPA buffer, and kept on ice for 10 min. TA muscles and cell lysates were centrifuged at 12,000 g for 10 minutes at 4 °C, and the supernatants were collected for further analysis. For western blot analyses, homogenates were separated in 8% polyacrylamide gels, transferred to Hybond nitrocellulose membranes (Amersham Biosciences), and probed with specific antibodies targeting AR (Abcam, ab108341, 1:500), α-TUBULIN (IGBMC, 1Tub2A2, 1:5000), and GAPDH (Cell Signaling, #2118, 1:5000).

Secondary antibodies conjugated to horseradish peroxidase (anit-Mouse-HPR, Cell Signalling, 7076S, 1:10,000; anit-Rabbit-HPR, Cell Signalling, 7074S, 1:10,000;) were detected using chemiluminescence detection systems (GE Healthcare, Amersham, ImageQuant 800 or Imager 600). Protein quantification was assessed by the FIJI/ImageJ distribution software (https://imagej.net/ImageJ) ^90^.

### Statistical analyses

Data processing and analysis were conducted using DESeq2, Microsoft Excel and GraphPad Prism 9. In all graphical representations, error bars indicate the standard error mean (SEM). Comparisons between groups were performed using appropriate statistical tests, selected based on the number of groups, data distribution (normality), variance homogeneity, and multiple comparisons. The specific statistical tests applied are indicated in the figure legends. Statistical significance was defined as follows: ns, p > 0.05 (non-significant); *, p < 0.05; **, p < 0.01; and ***, p < 0.001.

## Data availability

Custom macro for CSA measurements and Sirius Red quantification are available on GitLab and GitHub, respectively (CSA: the BIOP cellpose extension guide; Sirius Red: https://github.com/Hugues-Jacobs/Fibrosis_analysis). All sequencing datasets generated in this study have been deposited in the Gene Expression Omnibus (GEO) under the following accession numbers: GSE303611 for single-cell RNA-seq (token: afmhqcmenpapfyl), GSE303612 for SMART-seq (token: slileymohzanlal), GSE303614 for bulk RNA-seq (token: chqloqugddwjvur), GSE303615 for CUT&RUN analysis at 5 dpi (token: gtovmecctvyprot), GSE303618 for CUT&RUN analysis at 7 dpi (token: ynahwacgxtotdct), and GSE303620 for CUT&RUN analysis in non-injured animals (token: mjufuemerzyhlwn).

## Supporting information

Supplementary Informations

## ACKNOWLEDGMENTS

We thank Dr. J. Laporte (IGBMC) for generously providing the LHCN-M2 and C79 cell lines. We are grateful to Anastasia Bannwarth, Nikola Djordjevic, and Régis Lutzing for their valuable technical support, and Valentine Gilbart, Tao Ye and Céline Keime for excellent bioinformatics assistance. We extend our gratitude to Mohammed Selloum and Aurélie Aubertin for their valuable scientific expertise on biochemical analyses. We are grateful to Prof. S. Kato for providing the AR floxed mice, and Prof. G. Kardon for the *Pax7*-CreER^T2^ line. We acknowledge the support of the IGBMC core facilities, including the animal facility, microscopy and electron microscopy platforms, biochemistry, cell culture service, flow cytometry, and histopathology services, as well as the Mouse Clinical Institute (ICS, Illkirch-Graffenstaden, France) and GenomEast facility, a member of the ‘France Génomique’ consortium (ANR-10-INBS-0009).

This work was supported by the Interdisciplinary Thematic Institute IMCBio as part of the ITI 2021-2028 program of the University of Strasbourg, the Centre National pour la Recherche Scientifique (CNRS), Institut national de la santé et de la recherche médicale (Inserm), from IdEx Unistra (ANR-10-IDEX-0002), SFRI-STRAT’US (ANR 20-SFRI-0012) and EUR IMCBio (ANR-17-EURE-0023) projects, under the framework of the French Investments for the Future Programme. Additional support was provided by Inserm, CNRS, Unistra, IGBMC, AFM-Téléthon (grant application #24376), and the Agence Nationale de la Recherche (ANR-10-BLAN-1108, ANR-24-CE14-0247-01). J.G.R. received funding from the Programme CDFA-07-22 from the Université franco-allemande and Ministère de l’Enseignement Supérieur de la Recherche et de l’Innovation. K.C.G. received funding by the Association pour la Recherche à l’IGBMC (ARI), E.C by MESRI, Q.C by ANR (ANDROMETAMUS, ANR-24-CE14-0247-01), and R.S. by IMCBio. Anastasia Bannwarth received funding from AFM-Téléthon (grant application #24376) for technical assistance.

## AUTHOR CONTRIBUTIONS

D.D., J.G.R., and K.C.G. formulated the initial hypothesis. J.G.R., K.C.G., S.S.C., A.B., Q.C., G. R., E.C., and R.S. carried out functional, molecular, and histological experiments. D.D., J.G.R., N.M., and G.Z. executed bioinformatics analyses. J.G.R., N.M., H.J., and E.G. took responsibility of imaging analyses.

A.F. conducted *in situ* myofiber contraction experiments. J-F.A. and C.F. provided DHT pellets and performed initial *in vivo* tests. J.G.R., K.C.G., D.M., and D.D. took primary responsibility for data analysis and writing the manuscript.

## REFERENCES

1. Charge, S.B. & Rudnicki, M.A. Cellular and molecular regulation of muscle regeneration. Physiol Rev 84, 209–238 (2004).

2. Goel, A.J., Rieder, M.K., Arnold, H.H., Radice, G.L. & Krauss, R.S. Niche Cadherins Control the Quiescence-to-Activation Transition in Muscle Stem Cells. Cell Rep 21, 2236–2250 (2017).

3. Rozo, M., Li, L. & Fan, C.M. Targeting beta1-integrin signaling enhances regeneration in aged and dystrophic muscle in mice. Nat Med 22, 889–896 (2016).

4. Seale, P. et al. Pax7 is required for the specification of myogenic satellite cells. Cell 102, 777–786 (2000).

5. Wang, Y.X. & Rudnicki, M.A. Satellite cells, the engines of muscle repair. Nat Rev Mol Cell Biol 13, 127–133 (2011).

6. Anderson, J.E. Key concepts in muscle regeneration: muscle “cellular ecology” integrates a gestalt of cellular cross-talk, motility, and activity to remodel structure and restore function. Eur J Appl Physiol 122, 273–300 (2022).

7. Zhang, W., Liu, Y. & Zhang, H. Extracellular matrix: an important regulator of cell functions and skeletal muscle development. Cell Biosci 11, 65 (2021).

8. Koike, H., Manabe, I. & Oishi, Y. Mechanisms of cooperative cell-cell interactions in skeletal muscle regeneration. Inflamm Regen 42, 48 (2022).

9. Abou-Khalil, R. et al. Autocrine and paracrine angiopoietin 1/Tie-2 signaling promotes muscle satellite cell self-renewal. Cell Stem Cell 5, 298–309 (2009).

10. Kedlian, V.R. et al. Human skeletal muscle aging atlas. Nat Aging 4, 727–744 (2024).

11. Su, J. et al. A novel atlas of gene expression in human skeletal muscle reveals molecular changes associated with aging. Skelet Muscle 5, 35 (2015).

12. Moiseeva, V. et al. Senescence atlas reveals an aged-like inflamed niche that blunts muscle regeneration. Nature 613, 169–178 (2023).

13. Day, K., Shefer, G., Shearer, A. & Yablonka-Reuveni, Z. The depletion of skeletal muscle satellite cells with age is concomitant with reduced capacity of single progenitors to produce reserve progeny. Dev Biol 340, 330–343 (2010).

14. Petermann-Rocha, F. et al. Global prevalence of sarcopenia and severe sarcopenia: a systematic review and meta-analysis. J Cachexia Sarcopenia Muscle 13, 86–99 (2022).

15. Munoz-Canoves, P., Neves, J. & Sousa-Victor, P. Understanding muscle regenerative decline with aging: new approaches to bring back youthfulness to aged stem cells. FEBS J 287, 406–416 (2020).

16. Rizk, J., Sahu, R. & Duteil, D. An overview on androgen-mediated actions in skeletal muscle and adipose tissue. Steroids 199, 109306 (2023).

17. Conboy, I.M. & Rando, T.A. The regulation of Notch signaling controls satellite cell activation and cell fate determination in postnatal myogenesis. Dev Cell 3, 397–409 (2002).

18. Seo, J.Y., Kim, J.H. & Kong, Y.Y. Unraveling the Paradoxical Action of Androgens on Muscle Stem Cells. Mol Cells 42, 97–103 (2019).

19. Niel, L., Willemsen, K.R., Volante, S.N. & Monks, D.A. Sexual dimorphism and androgen regulation of satellite cell population in differentiating rat levator ani muscle. Dev Neurobiol 68, 115–122 (2008).

20. Diel, P., Baadners, D., Schlupmann, K., Velders, M. & Schwarz, J.P. C2C12 myoblastoma cell differentiation and proliferation is stimulated by androgens and associated with a modulation of myostatin and Pax7 expression. J Mol Endocrinol 40, 231–241 (2008).

21. Chen, Y., Zajac, J.D. & MacLean, H.E. Androgen regulation of satellite cell function. J Endocrinol 186, 21–31 (2005).

22. Klose, A., et al. Castration induces satellite cell activation that contributes to skeletal muscle maintenance. JCSM Rapid Commun 1 (2018).

23. Sakai, H. & Imai, Y. Cell-specific functions of androgen receptor in skeletal muscles. Endocr J 71, 437–445 (2024).

24. Sinha-Hikim, I., Taylor, W.E., Gonzalez-Cadavid, N.F., Zheng, W. & Bhasin, S. Androgen receptor in human skeletal muscle and cultured muscle satellite cells: up-regulation by androgen treatment. J Clin Endocrinol Metab 89, 5245–5255 (2004).

25. Ghaibour, K. et al. Androgen receptor coordinates muscle metabolic and contractile functions. J Cachexia Sarcopenia Muscle 14, 1707–1720 (2023).

26. Chambon, C. et al. Myocytic androgen receptor controls the strength but not the mass of limb muscles. Proc Natl Acad Sci U S A 107, 14327–14332 (2010).

27. Mourikis, P. & Tajbakhsh, S. Distinct contextual roles for Notch signalling in skeletal muscle stem cells. BMC Dev Biol 14, 2 (2014).

28. Pizza, F.X. & Buckley, K.H. Regenerating Myofibers after an Acute Muscle Injury: What Do We Really Know about Them? Int J Mol Sci 24 (2023).

29. Oprescu, S.N., Yue, F., Qiu, J., Brito, L.F. & Kuang, S. Temporal Dynamics and Heterogeneity of Cell Populations during Skeletal Muscle Regeneration. iScience 23, 100993 (2020).

30. McKellar, D.W. et al. Large-scale integration of single-cell transcriptomic data captures transitional progenitor states in mouse skeletal muscle regeneration. Commun Biol 4, 1280 (2021).

31. Yue, F. et al. Lipid droplet dynamics regulate adult muscle stem cell fate. Cell Rep 38, 110267 (2022).

32. Morroni, J. et al. Injury-experienced satellite cells retain long-term enhanced regenerative capacity. Stem Cell Res Ther 14, 246 (2023).

33. Brun, C.E. et al. GLI3 regulates muscle stem cell entry into G(Alert) and self-renewal. Nat Commun 13, 3961 (2022).

34. Barsky, S.T. & Monks, D.A. The role of androgens and global and tissue-specific androgen receptor expression on body composition, exercise adaptation, and performance. Biol Sex Differ 16, 28 (2025).

35. Chinvattanachot, G., Rivas, D. & Duque, G. Mechanisms of muscle cells alterations and regeneration decline during aging. Ageing Res Rev 102, 102589 (2024).

36. Serra, C. et al. Testosterone improves the regeneration of old and young mouse skeletal muscle. J Gerontol A Biol Sci Med Sci 68, 17–26 (2013).

37. Oura, M. et al. Testosterone/androgen receptor antagonizes immobility-induced muscle atrophy through Inhibition of myostatin transcription and inflammation in mice. Sci Rep 15, 10568 (2025).

38. Lai, Y. et al. Multimodal cell atlas of the ageing human skeletal muscle. Nature 629, 154–164 (2024).

39. Tan, M.H., Li, J., Xu, H.E., Melcher, K. & Yong, E.L. Androgen receptor: structure, role in prostate cancer and drug discovery. Acta Pharmacol Sin 36, 3–23 (2015).

40. Gruntmanis, U. The Role of 5α-Reductase Inhibition in Men Receiving Testosterone Replacement Therapy. JAMA 307, 968–970 (2012).

41. Bartsch, G., Rittmaster, R.S. & Klocker, H. Dihydrotestosterone and the concept of 5alpha-reductase inhibition in human benign prostatic hyperplasia. Eur Urol 37, 367–380 (2000).

42. Dubois, V., Laurent, M., Boonen, S., Vanderschueren, D. & Claessens, F. Androgens and skeletal muscle: cellular and molecular action mechanisms underlying the anabolic actions. Cell Mol Life Sci 69, 1651–1667 (2012).

43. Sheppard, R.L., Spangenburg, E.E., Chin, E.R. & Roth, S.M. Androgen receptor polyglutamine repeat length affects receptor activity and C2C12 cell development. Physiol Genomics 43, 1135–1143 (2011).

44. Fu, S., Lin, X., Yin, L. & Wang, X. Androgen receptor regulates the proliferation of myoblasts under appropriate or excessive stretch through IGF-1 receptor mediated p38 and ERK1/2 pathways. Nutr Metab (Lond) 18, 85 (2021).

45. Fu, S., Hu, J., Wang, G., Qian, Z. & Wang, X. Androgen receptor regulates the differentiation of myoblasts under cyclic mechanical stretch and its upstream and downstream signals. Int J Biol Macromol 281, 136257 (2024).

46. Dubois, V. et al. Androgen Deficiency Exacerbates High-Fat Diet-Induced Metabolic Alterations in Male Mice. Endocrinology 157, 648–665 (2016).

47. Rocchi, A. et al. Glycolytic-to-oxidative fiber-type switch and mTOR signaling activation are early-onset features of SBMA muscle modified by high-fat diet. Acta Neuropathol 132, 127–144 (2016).

48. Campbell, M.D., Djukovic, D., Raftery, D. & Marcinek, D.J. Age-related changes of skeletal muscle metabolic response to contraction are also sex-dependent. J Physiol 603, 69–86 (2025).

49. Grevendonk, L. et al. Impact of aging and exercise on skeletal muscle mitochondrial capacity, energy metabolism, and physical function. Nat Commun 12, 4773 (2021).

50. Kobayashi, A., Azuma, K., Ikeda, K. & Inoue, S. Mechanisms Underlying the Regulation of Mitochondrial Respiratory Chain Complexes by Nuclear Steroid Receptors. Int J Mol Sci 21 (2020).

51. Jomova, K. et al. Reactive oxygen species, toxicity, oxidative stress, and antioxidants: chronic diseases and aging. Arch Toxicol 97, 2499–2574 (2023).

52. Lee, J. et al. Androgen Deprivation Therapy-Induced Muscle Loss and Fat Gain Predict Cardiovascular Events in Prostate Cancer Patients. J Cachexia Sarcopenia Muscle 16, e13844 (2025).

53. Argiles, J.M., Campos, N., Lopez-Pedrosa, J.M., Rueda, R. & Rodriguez-Manas, L. Skeletal Muscle Regulates Metabolism via Interorgan Crosstalk: Roles in Health and Disease. J Am Med Dir Assoc 17, 789–796 (2016).

54. Penvose, A., Keenan, J.L., Bray, D., Ramlall, V. & Siggers, T. Comprehensive study of nuclear receptor DNA binding provides a revised framework for understanding receptor specificity. Nat Commun 10, 2514 (2019).

55. Gelman, L. et al. Integrating nuclear receptor mobility in models of gene regulation. Nucl Recept Signal 4, e010 (2006).

56. Hager, G.L., Nagaich, A.K., Johnson, T.A., Walker, D.A. & John, S. Dynamics of nuclear receptor movement and transcription. Biochim Biophys Acta 1677, 46–51 (2004).

57. Stavreva, D.A. et al. Nuclear Receptors Dynamics and Gene Regulation in Response to Ultradian Hormone Pulses. Endocrinology 166, bqaf043.034 (2025).

58. Forouhan, M. et al. AR cooperates with SMAD4 to maintain skeletal muscle homeostasis. Acta Neuropathol 143, 713–731 (2022).

59. Nath, S.R. et al. Androgen receptor polyglutamine expansion drives age-dependent quality control defects and muscle dysfunction. J Clin Invest 128, 3630–3641 (2018).

60. Arnold, F.J., Pluciennik, A. & Merry, D.E. Impaired Nuclear Export of Polyglutamine-Expanded Androgen Receptor in Spinal and Bulbar Muscular Atrophy. Sci Rep 9, 119 (2019).

61. Shiina, H. et al. Premature ovarian failure in androgen receptor-deficient mice. Proc Natl Acad Sci U S A 103, 224–229 (2006).

62. Murphy, M.M., Lawson, J.A., Mathew, S.J., Hutcheson, D.A. & Kardon, G. Satellite cells, connective tissue fibroblasts and their interactions are crucial for muscle regeneration. Development 138, 3625–3637 (2011).

63. Tosic, M. et al. Lsd1 regulates skeletal muscle regeneration and directs the fate of satellite cells. Nat Commun 9, 366 (2018).

64. Chenu, C. et al. Testosterone Prevents Cutaneous Ischemia and Necrosis in Males Through Complementary Estrogenic and Androgenic Actions. Arterioscler Thromb Vasc Biol 37, 909–919 (2017).

65. Rovito, D. et al. Myod1 and GR coordinate myofiber-specific transcriptional enhancers. Nucleic Acids Res 49, 4472–4492 (2021).

66. Simon, A. et al. Transcriptomic characterization of postnatal muscle maturation. Dis Model Mech 18 (2025).

67. Cigrang, M. et al. Pan-inhibition of super-enhancer-driven oncogenic transcription by next-generation synthetic ecteinascidins yields potent anti-cancer activity. Nat Commun 16, 512 (2025).

68. Ghaibour, K. et al. An Efficient Protocol for CUT&RUN Analysis of FACS-Isolated Mouse Satellite Cells. J Vis Exp (2023).

69. Janssens, D.H. et al. Automated in situ chromatin profiling efficiently resolves cell types and gene regulatory programs. Epigenetics Chromatin 11, 74 (2018).

70. Langmead, B. & Salzberg, S.L. Fast gapped-read alignment with Bowtie 2. Nat Methods 9, 357–359 (2012).

71. Ramirez, F., Dundar, F., Diehl, S., Gruning, B.A. & Manke, T. deepTools: a flexible platform for exploring deep-sequencing data. Nucleic Acids Res 42, W187–191 (2014).

72. Quinlan, A.R. & Hall, I.M. BEDTools: a flexible suite of utilities for comparing genomic features. Bioinformatics 26, 841–842 (2010).

73. Meers, M.P., Tenenbaum, D. & Henikoff, S. Peak calling by Sparse Enrichment Analysis for CUT&RUN chromatin profiling. Epigenetics Chromatin 12, 42 (2019).

74. Zhang, Y. et al. Model-based analysis of ChIP-Seq (MACS). Genome Biol 9, R137 (2008).

75. Ye, T. et al. seqMINER: an integrated ChIP-seq data interpretation platform. Nucleic Acids Res 39, e35 (2011).

76. Cai, Q. et al. LSD1 inhibition circumvents glucocorticoid-induced muscle wasting of male mice. Nat Commun 15, 3563 (2024).

77. Livak, K.J. & Schmittgen, T.D. Analysis of relative gene expression data using real-time quantitative PCR and the 2(-Delta Delta C(T)) Method. Methods 25, 402–408 (2001).

78. Dobin, A. et al. STAR: ultrafast universal RNA-seq aligner. Bioinformatics 29, 15–21 (2013).

79. Anders, S., Pyl, P.T. & Huber, W. HTSeq--a Python framework to work with high-throughput sequencing data. Bioinformatics 31, 166–169 (2015).

80. Love, M.I., Huber, W. & Anders, S. Moderated estimation of fold change and dispersion for RNA-seq data with DESeq2. Genome Biol 15, 550 (2014).

81. Szklarczyk, D. et al. The STRING database in 2023: protein-protein association networks and functional enrichment analyses for any sequenced genome of interest. Nucleic Acids Res 51, D638–D646 (2023).

82. Wang, J., Duncan, D., Shi, Z. & Zhang, B. WEB-based GEne SeT AnaLysis Toolkit (WebGestalt): update 2013. Nucleic Acids Res 41, W77–83 (2013).

83. de Hoon, M.J., Imoto, S., Nolan, J. & Miyano, S. Open source clustering software. Bioinformatics 20, 1453–1454 (2004).

84. Lu, Y., Lu, S., Fotouhi, F., Deng, Y. & Brown, S.J. Incremental genetic K-means algorithm and its application in gene expression data analysis. BMC Bioinformatics 5, 172 (2004).

85. Raine, A., Liljedahl, U. & Nordlund, J. Data quality of whole genome bisulfite sequencing on Illumina platforms. PLoS One 13, e0195972 (2018).

86. Lun, A.T.L. et al. EmptyDrops: distinguishing cells from empty droplets in droplet-based single-cell RNA sequencing data. Genome Biol 20, 63 (2019).

87. Satija, R., Farrell, J.A., Gennert, D., Schier, A.F. & Regev, A. Spatial reconstruction of single-cell gene expression data. Nat Biotechnol 33, 495–502 (2015).

88. Jin, S. et al. Inference and analysis of cell-cell communication using CellChat. Nat Commun 12, 1088 (2021).

89. Riba, A. et al. Cell cycle gene regulation dynamics revealed by RNA velocity and deep-learning. Nat Commun 13, 2865 (2022).

90. Schneider, C.A., Rasband, W.S. & Eliceiri, K.W. NIH Image to ImageJ: 25 years of image analysis. Nat Methods 9, 671–675 (2012).

