## Supplementary Informations for "Androgen receptor imprints satellite cells stemness and preserves their reservoir for lifelong regeneration and optimal repair"

**Supplementary Information**

This file includes Supplementary Figures 1-7 and Supplementary Tables 1-6.

### Supplementary Figures

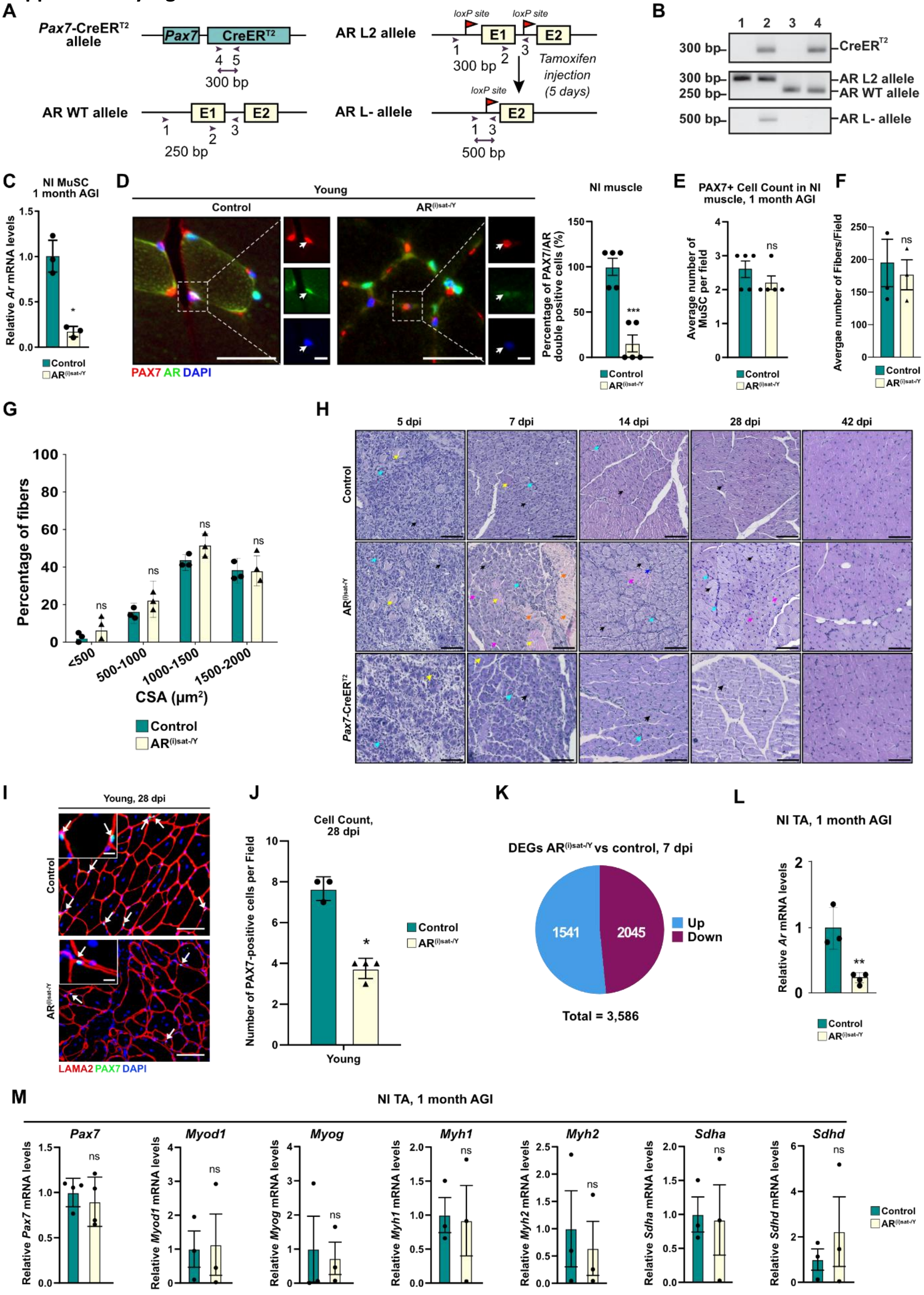

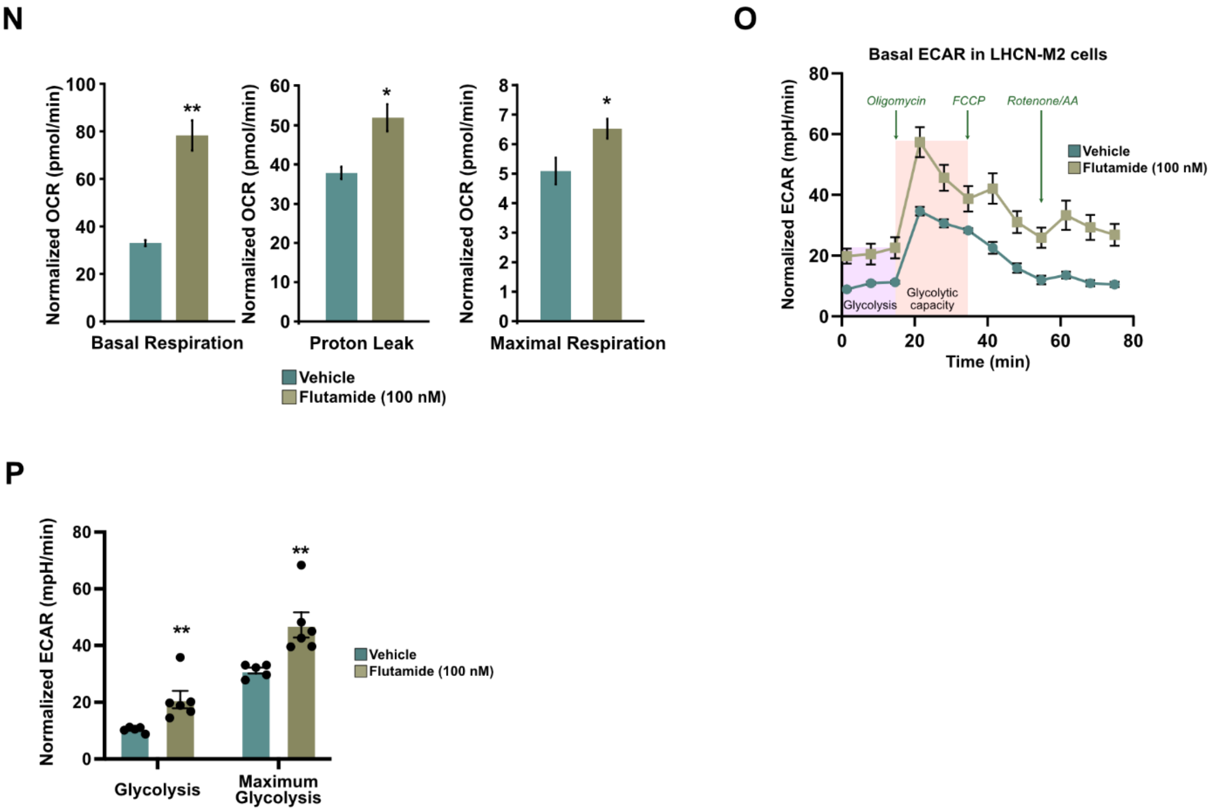


**Supplementary Figure 1: Absence of MuSC-AR impairs muscle regenerative capacities at puberty.**

**(A)** Schematic representation of *Pax7*-CreER$^{T2}$, wild-type (WT), floxed (L2) and CreER$^{T2}$-mediated exon 1 deleted (L-) AR alleles. LoxP sites are shown by arrowheads. Primers used for allele characterization are depicted by arrows and sequences are in Table S4.

**(B)** Representative electrophoresis of PCR-amplified genomic DNA from soleus muscles of control (AR$^{L2/Y}$) (line 1), mutant (AR$^{(i)sat-/Y}$) (line 2), AR$^{+/Y}$ (WT) (line 3), and AR$^{+/Y}$/Pax7-CreER$^{T2}$ (line 4) mice, illustrating distinct amplicon sizes: Cre$^{ERT2}$ (300 bp), AR L2 (300 bp), wild-type (WT; 250 bp), and AR L– (500 bp). The absence of the AR L - band in mice 1, 3, and 4 confirms their control genotype, whereas its presence in mouse 2 indicates successful recombination in the mutant.

**(C)** Relative transcript levels of *Ar* in MuSC FACS-isolated from NI TA muscles of young male mice, 1 month after gene invalidation (AGI). Data are presented as mean ± SEM. Statistical test used was two-tailed Mann-Whitney test; ns = non-significant, * = $p < 0.05$.

**(D-E)** Representative immunofluorescent labeling of AR (in green) and PAX7 (in red), and corresponding quantification of the percentage of PAX7$^{+}$/AR$^{+}$ double-positive cells **(D)**, and average number of MuSC per field **(E)**, in non-injured (NI) TA of young control and AR$^{(i)sat-/Y}$ male mice. White arrows denote MuSC. Nuclei were stained with DAPI. Scale bars, 100 µm for the main, 30 µm for magnified inset. Statistical test used was two-tailed Mann-Whitney test; *** = $p < 0.001$.

**(F)** Quantification of the number of fibers per filed in control and AR$^{(i)sat-/Y}$ male mice in non-injured condition (NI). Scale bars, 250 µm. Statistical test used was two-tailed Mann-Whitney test; ns = non-significant.

**(G)** Distribution of myofibers cross-section area (CSA) in TA muscles of control and AR$^{(i)sat-/Y}$ male in non-injured condition (NI). Data are presented as mean ± SEM. Statistical test used Two-way ANOVA with Sidak’s correction; ns = non-significant.

**(H)** Representative hematoxylin and eosin (H&E) staining of tibialis anterior (TA) muscles of control and $AR^{(i)sat-/Y}$ male mice in non-injured condition (NI), and at indicated time points after injury. Black arrows indicate centrally nucleated fibers. The *Pax7*-$CreER^{T2}$ individuals lacking floxed AR were included to confirm that Cre recombinase expression does not produce any leakage effects. Cyan blue arrows point to immune infiltration. Yellow arrows denote necrotic fibers. Pink arrows label misshaped regenerating myofibers. Red arrows designate adipocytes. Orange arrows mark fibrosis. Scale bars, 250 µm.

**(I-J)** Representative immunofluorescent labeling of laminin 2-alpha (LAMA2, in red) and PAX7 (in green) **(I)**, and corresponding quantification of the number of PAX7-positive cells **(J),** in 28-days injured TA of young control and $AR^{(i)sat-/Y}$ male mice. White arrows denote MuSC. Nuclei were stained with DAPI. Scale bars, 250 µm for the main, 10 µm for magnified inset. Statistical test used was two-tailed Mann-Whitney test; * = $p < 0.05$.

**(K)** Pie chart depicting the number of down- and up-regulated genes in 7-days-injured TA muscles of $AR^{(i)sat-/Y}$ male mice when compared to control littermates.

**(L-M)** Relative transcript levels of indicated genes in NI TA muscles of young male mice, 1 month after gene invalidation (AGI). Data are presented as mean ± SEM. Statistical test used was two-tailed Mann-Whitney test; ns = non-significant, ** = $p < 0.01$.

**(N)** Quantification of mitochondrial respiration derived bioenergetic parameters, Basal respiration (OCR after subtraction of non-mitochondrial respiration), Proton leak (OCR following oligomycin addition), and Maximal respiration (OCR following FCCP treatment) in vehicle and 100 nM flutamide-treated LHCN-M2 cells. Data are normalized to total nuclear counts. Statistical test used was two-tailed Mann-Whitney test; * = $p < 0.05$; ** = $p < 0.01$.

**(O-P)** Basal extracellular acidification rate (ECAR) profile **(O)**, and quantification of glycolysis and glycolytic capacity **(P)** in vehicle and 100 nM flutamide-treated LHCN-M2 cells, assessed using the Seahorse Mito Stress Test, following sequential injection of oligomycin, carbonyl cyanide-p-trifluoromethoxyphenylhydrazone (FCCP), and rotenone/AA. Data are normalized to total nuclear counts, and presented as mean ± SEM. Statistical test used was two-tailed Mann-Whitney test; ** = $p < 0.01$.

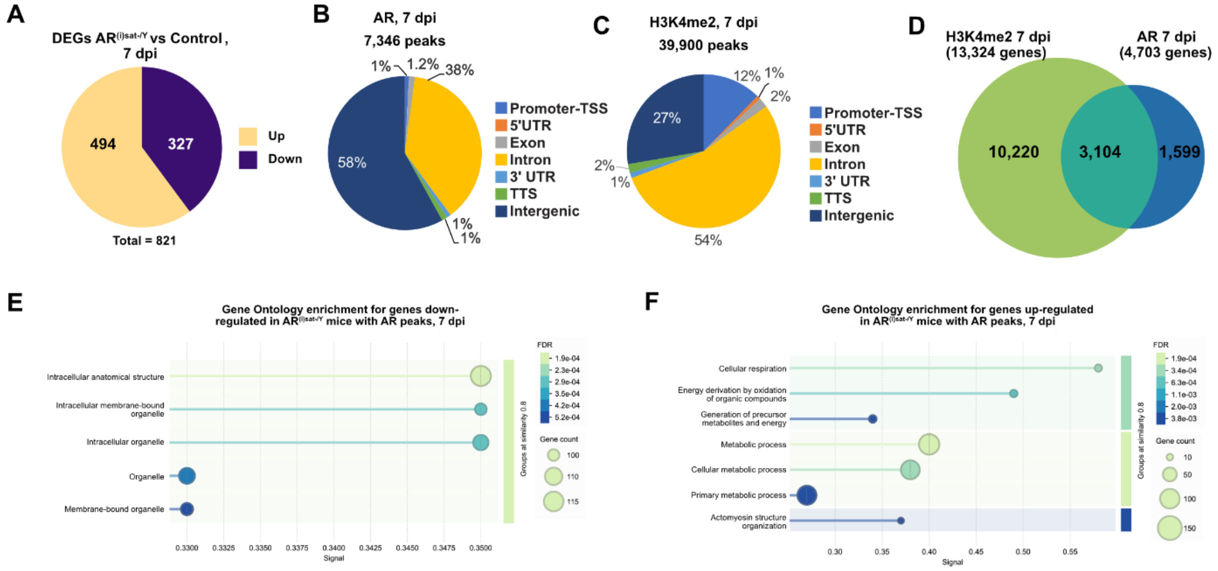


**Supplementary Figure 2: Characterization of AR cistrome and transcriptome in MuSC at puberty**

**(A)** Pie chart depicting the number of down- and up-regulated genes in young $AR^{(i)sat-/Y}$ versus control MuSC FACS-isolated from 7-day-injured mouse TA.

**(B-C)** Pie chart depicting the genomic location of AR binding sites **(B)**, and the peak distribution of H3K4me2 in the MuSC genome of 7-days-injured young mouse TA **(C)**.

**(D)** Overlap between genes bound by AR and active promoters marked by H3K4me2 peaks in the MuSC genome of 7-days-injured young mouse TA.

**(E-F)** Gene ontology enrichment analysis of genes associated with AR peaks that are down- **(E)** or upregulated **(F)** in male mice, in 7-days injured TA.

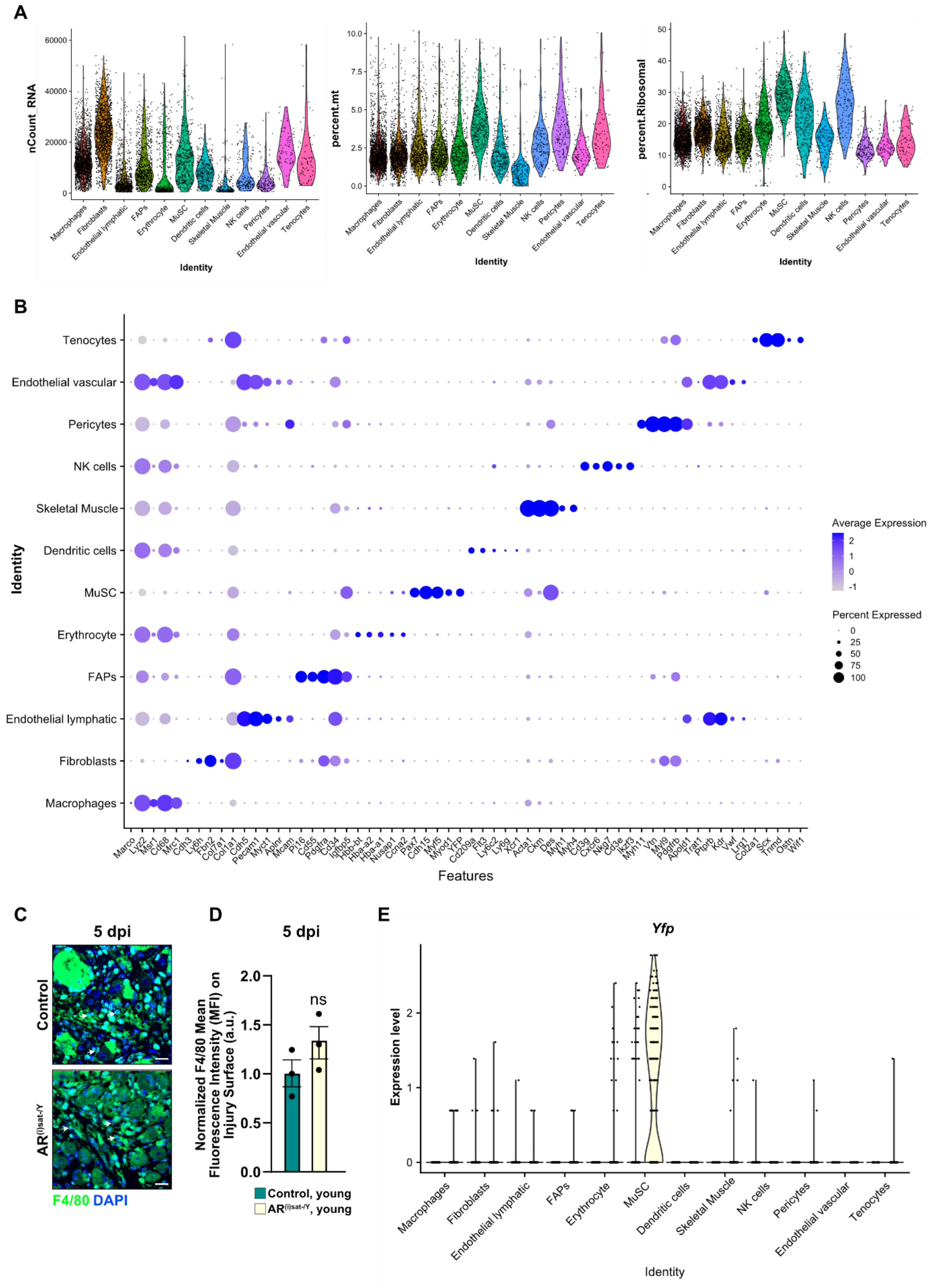
A
nCount_RNA
percent.mt
percent.Ribosomal
Identity
B
Identity
Features
Average Expression
Percent Expressed
C
5 dpi
Control
F4/80 DAPI
D
5 dpi
Normalized F4/80 Mean Fluorescence Intensity (MFI) on Injury Surface (a.u.)
ns
Control, young
E
Yfp
Expression level
Identity

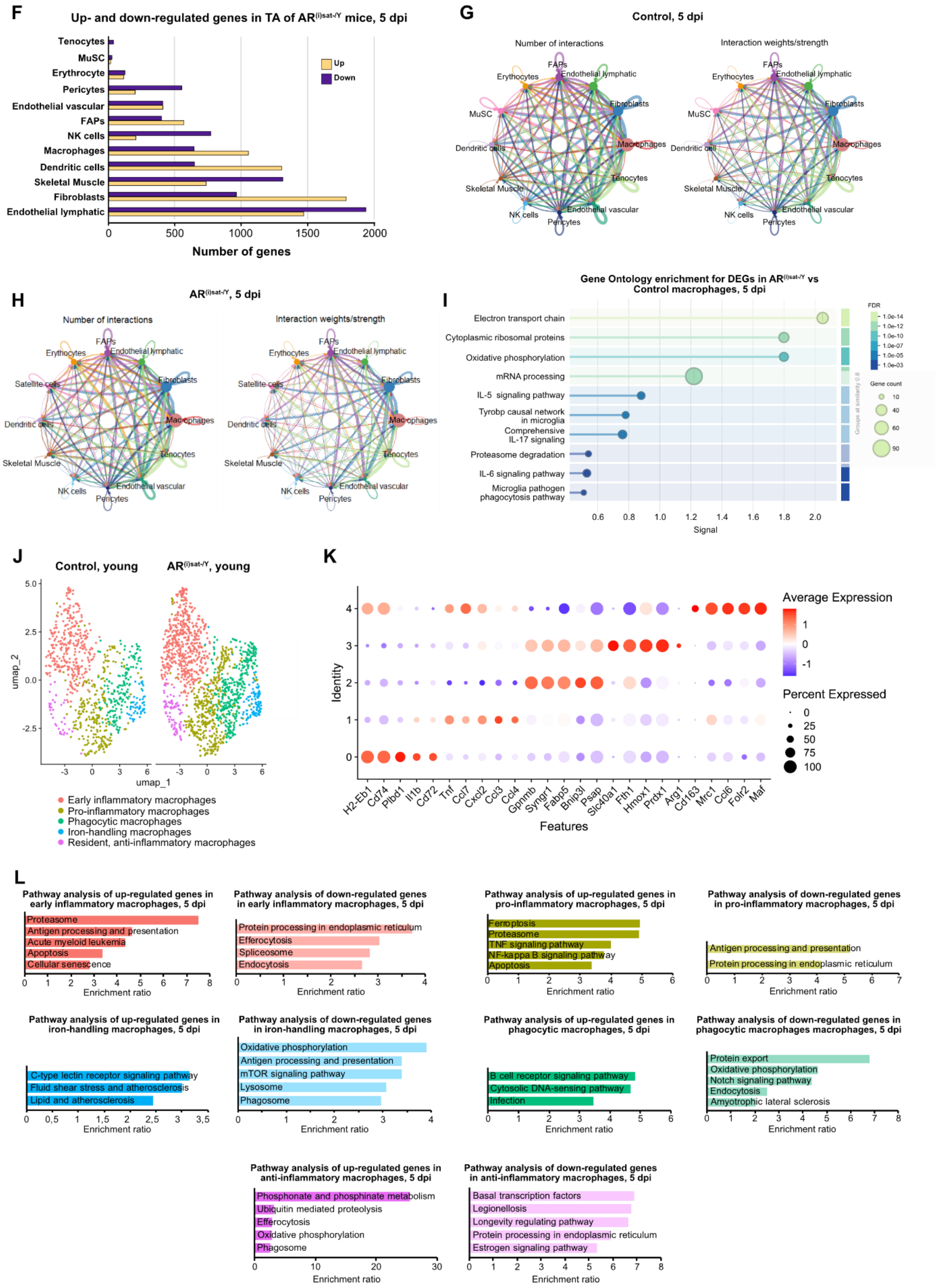
F
Up- and down-regulated genes in TA of AR(i)sat-/Y mice, 5 dpi
Tenocytes
MuSC
Erythrocyte
Pericytes
Endothelial vascular
FAPs
NK cells
Macrophages
Dendritic cells
Skeletal Muscle
Fibroblasts
Endothelial lymphatic
Up
Down
0
500
1000
1500
2000
Number of genes
G
Control, 5 dpi
Number of interactions
Interaction weights/strength
H
AR(i)sat-/Y, 5 dpi
I
Gene Ontology enrichment for DEGs in AR(i)sat-/Y vs Control macrophages, 5 dpi
Electron transport chain
Cytoplasmic ribosomal proteins
Oxidative phosphorylation
mRNA processing
IL-5 signaling pathway
Tyrobp causal network in microglia
Comprehensive IL-17 signaling
Proteasome degradation
IL-6 signaling pathway
Microglia pathogen phagocytosis pathway
Signal
FDR
Gene count
J
Control, young
AR(i)sat-/Y, young
umap_1
umap_2
Early inflammatory macrophages
Pro-inflammatory macrophages
Phagocytic macrophages
Iron-handling macrophages
Resident, anti-inflammatory macrophages
K
Average Expression
Percent Expressed
Identity
Features
H2-Eb1
Cd74
Plbd1
Il1b
Cd72
Tnf
Ccl7
Cxcl2
Ccl3
Ccl4
Gpnmb
Syngr1
Fabp5
Bnip3l
Psap
Slc40a1
Fth1
Hmox1
Prdx1
Arg1
Cd163
Mrc1
Ccl6
Folr2
Maf
L
Pathway analysis of up-regulated genes in early inflammatory macrophages, 5 dpi
Proteasome
Antigen processing and presentation
Acute myeloid leukemia
Apoptosis
Cellular senescence
Pathway analysis of down-regulated genes in early inflammatory macrophages, 5 dpi
Protein processing in endoplasmic reticulum
Efferocytosis
Spliceosome
Endocytosis
Pathway analysis of up-regulated genes in pro-inflammatory macrophages, 5 dpi
Ferroptosis
Proteasome
TNF signaling pathway
NF-kappa B signaling pathway
Apoptosis
Pathway analysis of down-regulated genes in pro-inflammatory macrophages, 5 dpi
Antigen processing and presentation
Protein processing in endoplasmic reticulum
Pathway analysis of up-regulated genes in iron-handling macrophages, 5 dpi
C-type lectin receptor signaling pathway
Fluid shear stress and atherosclerosis
Lipid and atherosclerosis
Pathway analysis of down-regulated genes in iron-handling macrophages, 5 dpi
Oxidative phosphorylation
Antigen processing and presentation
mTOR signaling pathway
Lysosome
Phagosome
Pathway analysis of up-regulated genes in phagocytic macrophages, 5 dpi
B cell receptor signaling pathway
Cytosolic DNA-sensing pathway
Infection
Pathway analysis of down-regulated genes in phagocytic macrophages macrophages, 5 dpi
Protein export
Oxidative phosphorylation
Notch signaling pathway
Endocytosis
Amyotrophic lateral sclerosis
Pathway analysis of up-regulated genes in anti-inflammatory macrophages, 5 dpi
Phosphonate and phosphinate metabolism
Ubiquitin mediated proteolysis
Efferocytosis
Oxidative phosphorylation
Phagosome
Pathway analysis of down-regulated genes in anti-inflammatory macrophages, 5 dpi
Basal transcription factors
Legionellosis
Longevity regulating pathway
Protein processing in endoplasmic reticulum
Estrogen signaling pathway
Enrichment ratio

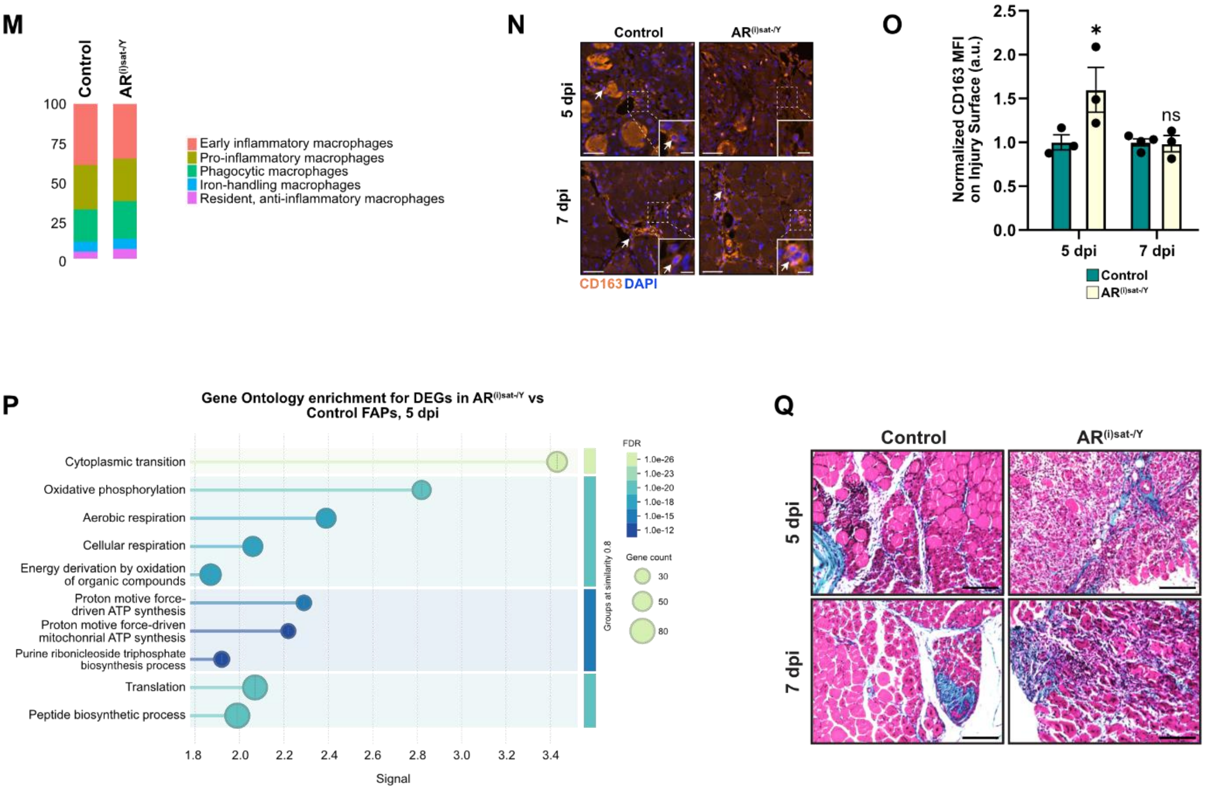


**Supplementary Figure 3: MuSC microenvironment is altered in the absence of AR**

**(A)** Violin plots representing the normalized read counts of RNA, the percentage of mitochondrial genes and the percentage of ribosomal genes of the indicated cell populations in 5-days-injured TA of young control and $AR^{(i)sat-/Y}$ mice.

**(B)** Dot plot heatmap of the indicated cell population marker genes in 5-days-injured TA of control and $AR^{(i)sat-/Y}$ mice. The dot size corresponds to the fraction of cells in each cluster that have higher than average expression of the indicated gene. Scale: grey/indigo is low expression, blue is high gene expression.

**(C-D)** Representative immunofluorescent labeling of F4/80 (in green) **(C)**, and corresponding quantification of the mean fluorescence intensity (MFI) of F4/80 signal **(D)**, in 5-days-injured TA of control and $AR^{(i)sat-/Y}$ young male mice. White arrows point to PAX7-positive cells. Nuclei were stained with DAPI. Scale bars, 50 µm. Data are presented as mean ± SEM. Statistical test used is two-tailed Mann-Whitney test; ns, non-significant.

**(E)** Violin plots representing the normalized read counts of *Yfp* in MuSC, selected from scRNA-seq analysis performed in 5- days-injured TA of control and $AR^{(i)sat-/Y}$ mice.

**(F)** Number of up- and down-regulated genes in cell populations of 5-days-injured TA of $AR^{(i)sat-/Y}$ mice, relative control littermates.

**(G-H)** Circle plots showing the number and strength of ligand-receptor interactions between pairwise cell populations among the 12 major cell populations in 5-days-injured TA of control **(G)**, and $AR^{(i)sat-/Y}$ mice **(H)**. The strength of ligand-receptor interactions between cell population pairs was visualized with edge width that was proportional to the number of L-R pairs.

**(I)** Gene Ontology enrichment of macrophages differentially expressed genes from scRNA-seq analysis performed in 5-days-injured TA of control and $AR^{(i)sat-/Y}$ mice.

**(J)** UMAP visualization of macrophages reclusters from 5-days-injured control and $AR^{(i)sat-/Y}$ TA of young mice, colored by cluster identity.

**(K)** Dot plot heatmap of macrophages marker genes in 5-days-injured TA of control and $AR^{(i)sat-/Y}$ mice. The dot size corresponds to the fraction of cells in each cluster that have higher than average expression of the indicated gene. Scale: blue is low expression, red is high gene expression.

**(L)** Pathway analysis performed on down- and up-regulated genes identified in macrophages sub-populations of 5-day-injured TA of young control and $AR^{(i)sat-/Y}$ mice.

**(M)** Percentage of macrophages sub-populations in 5-days-injured TA of young control and $AR^{(i)sat-/Y}$ mice, analyzed from sc-RNA seq data.

**(N-O)** Representative immunofluorescent labeling of CD163 (in orange) **(N)**, and corresponding quantification of the mean fluorescence intensity (MFI) of CD163 signal **(O)**, in 5- and 7-days-injured TA of control and $AR^{(i)sat-/Y}$ young male mice. Nuclei were stained with DAPI. White arrows point to CD163-pisitive nacrophages. Data are presented as mean ± SEM. Statistical test used is Ordinary One-way ANOVA relative to 5 dpi control; ns, non-significant; * = $p < 0.05$. Scale bars, 100 µm for the main, 10 µm for magnified inset **(K)**.

**(P)** Pathway analysis performed on differentially expressed genes identified in FAPs in scRNA-seq analysis performed in 5-days-injured TA of young control and $AR^{(i)sat-/Y}$ mice.

**(Q)** Representative Masson Trichrome Staining in 5- and 7-days-injured TA of control and $AR^{(i)sat-/Y}$ mice. Connective tissue is stained in blue, nuclei are stained in dark red/purple, and cytoplasm is stained in red/pink. Scale bars, 300 µm.

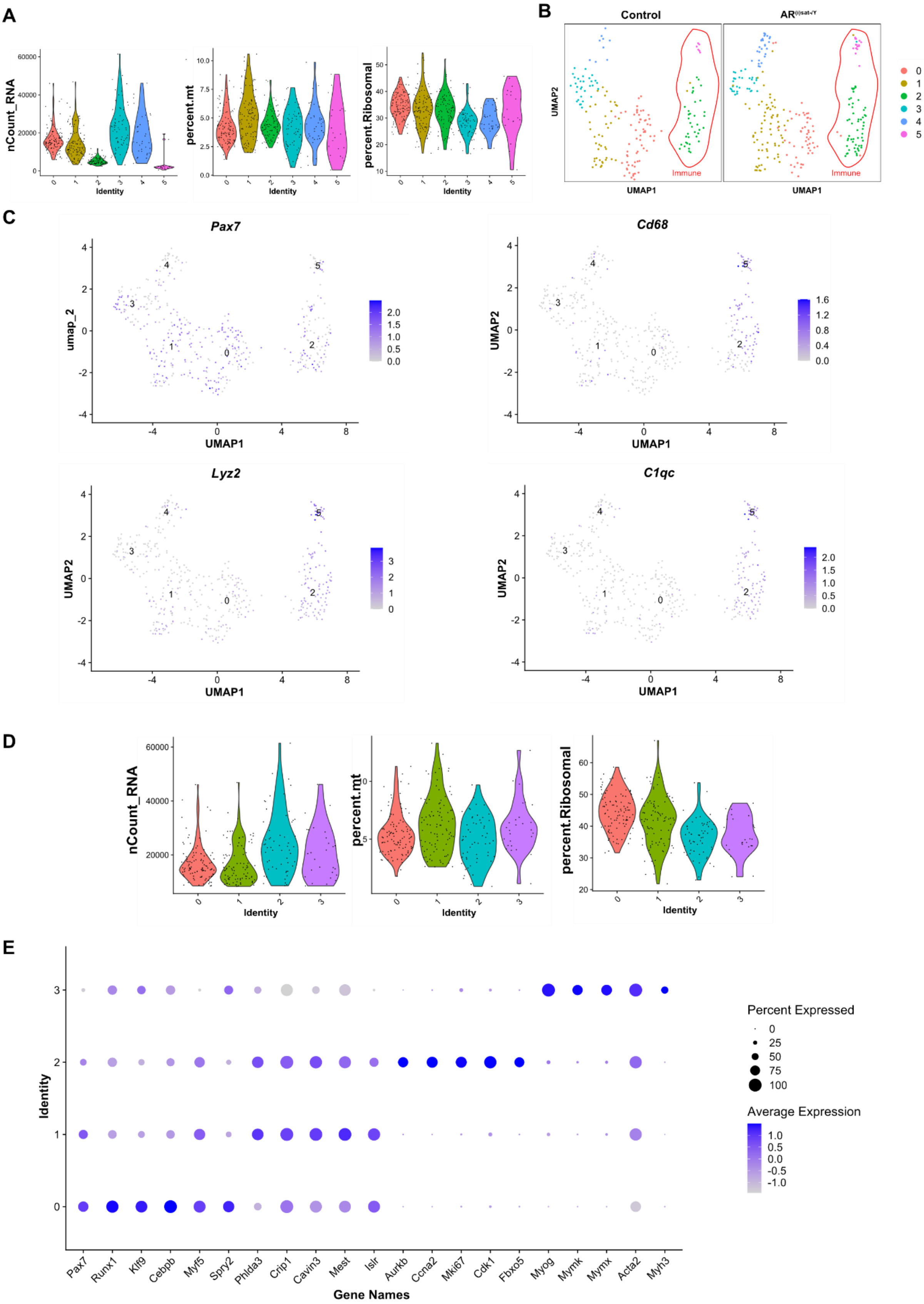
A
nCount_RNA
percent.mt
percent.Ribosomal
Identity
B
Control
UMAP2
UMAP1
Immune
C
Pax7
Cd68
Lyz2
C1qc
umap_2
UMAP2
UMAP1
D
nCount_RNA
percent.mt
percent.Ribosomal
Identity
E
Identity
Pax7
Runx1
Klf9
Cebpb
Myf5
Spry2
Phlda3
Crip1
Cavin3
Mest
Islr
Aurkb
Ccna2
Mki67
Cdk1
Fbxo5
Myog
Mymk
Mymx
Acta2
Myh3
Gene Names
Percent Expressed
Average Expression

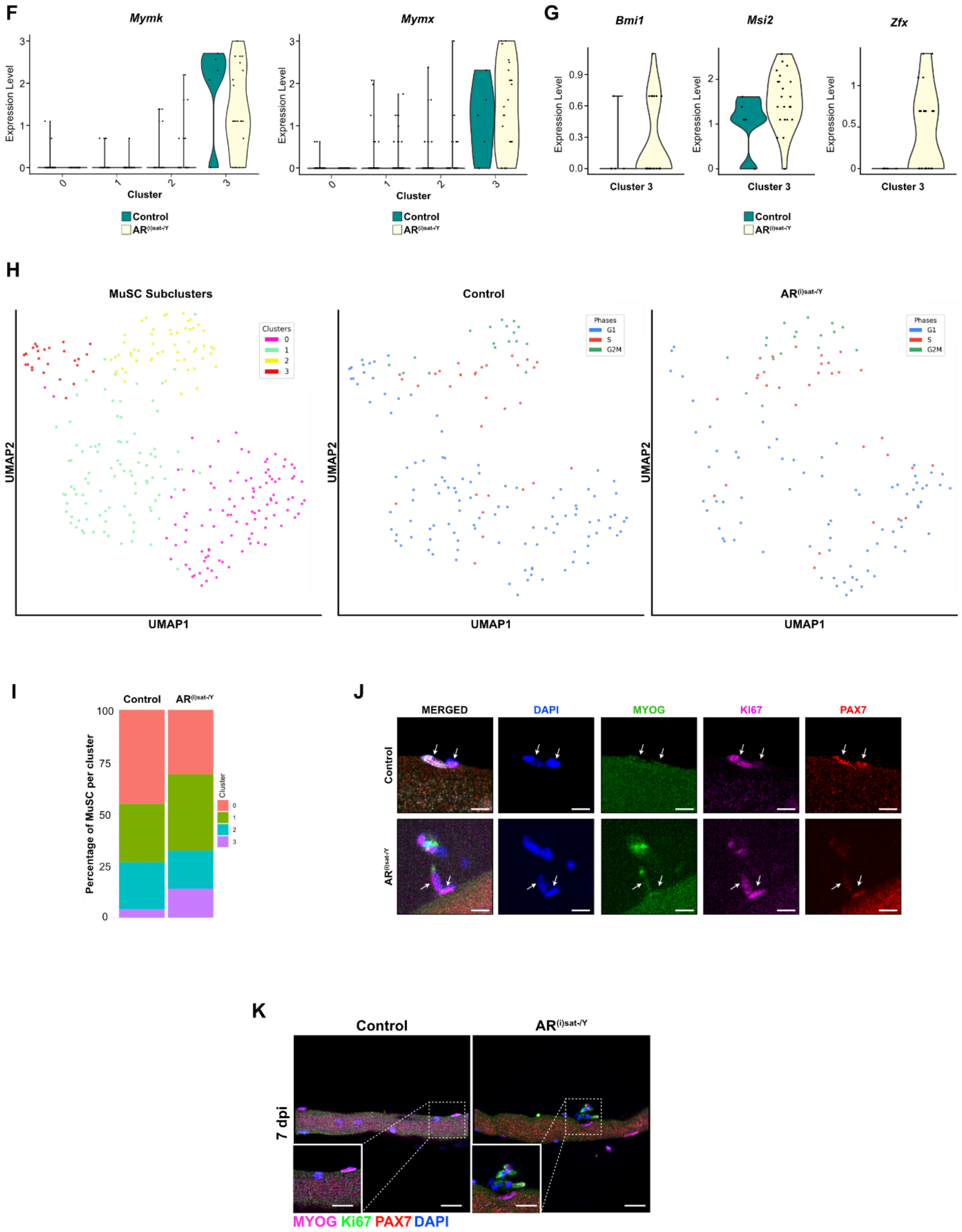


**Supplementary Figure 4: AR deficiency in MuSC favors symmetric differentiation over asymmetric division or symmetric self-renewal**

**(A)** Violin plots representing the normalized read counts of RNA, the percentage of mitochondrial genes and the percentage of ribosomal genes of MuSC clusters (clusters 0-5) in 5-days-injured TA of control and AR$^{(i)sat-/Y}$ mice.

**(B)** UMAP visualization of MuSC clusters (clusters 0-5) from 5-days-injured control and AR$^{(i)sat-/Y}$ TA of mice, colored by cluster identity.

**(C)** Feature plots representing *Pax7*, *Cd68*, *Lyz2*, and *C1qc* genes expression levels and pattern MuSC clusters (clusters 0-5) from 5-days-injured control and AR$^{(i)sat-/Y}$ TA of mice. Scale: grey/indigo is low expression, blue is high gene expression.

**(D)** Violin plots representing the expression level of RNA, the percentage of mitochondrial genes and the percentage of ribosomal genes of MuSC reclusters (cluster 0-3) in 5- days-injured TA of control and AR$^{(i)sat-/Y}$ mice.

**(E)** Dot plot heatmap of the MuSC reclusters (clusters 0-3) marker genes in 5-days-injured control and AR$^{(i)sat-/Y}$ TA of mice. The dot size corresponds to the fraction of cells in each cluster that have higher than average expression of the indicated gene. Scale: grey/indigo is low expression, blue is high gene expression.

**(F)** Violin plots representing the expression level of *Mymk* and *Mymx* in MuSC reclusters (clusters 0-3), selected from scRNA-seq analysis performed in 5- days-injured TA of control and AR$^{(i)sat-/Y}$ mice.

**(G)** Violin plots representing the expression of indicated genes, in MuSC cluster 3, selected from scRNA-seq analysis performed in 5- days-injured TA of control and AR$^{(i)sat-/Y}$ mice.

**(H)** UMAP visualization of MuSC reclusters (clusters 0-3), illustrating their distribution in 5-days-injured control and AR$^{(i)sat-/Y}$ TA of mice. Cells are color-coded by cluster identity or by cell cycle phase (G1, S, G2/M) to highlight phase-specific repartition within each genotype.

**(I)** Percentage of MuSC reclusters (clusters 0-3) in 5-days-injured control and AR$^{(i)sat-/Y}$ TA of mice.

**(J-K)** Representative immunofluorescence detection of MYOG (green), KI67 (magenta), and PAX7 (red) in isolated muscle fibers from TA muscles of control and AR$^{(i)sat-/Y}$ young mice at 7 dpi. Nuclei were stained with DAPI. Scale bars, 5 µm **(J)**; 25 µm for the main, 10 µm for magnified inset **(K)**.

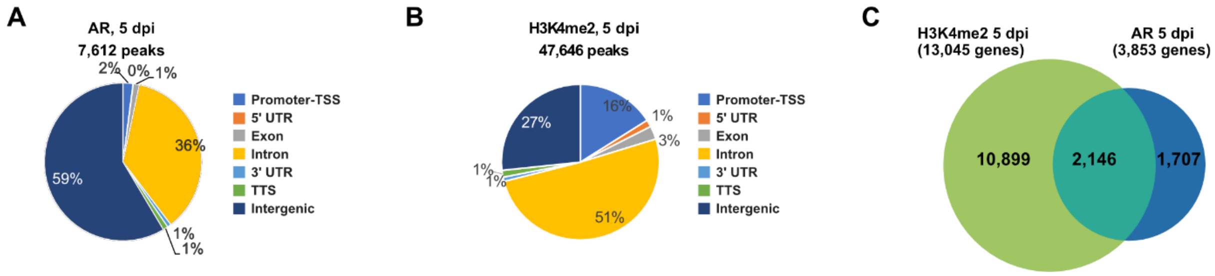


**Supplementary Figure 5: MuSC-AR exhibits time-dependent binding to target genes following injury**
**(A-B)** Pie chart depicting the genomic location of AR binding sites **(A)**, and the peak distribution of H3K4me2 in the MuSC genome **(B)** in 5-day-injured TA of young male mice.

**(C)** Overlap between genes bound by AR and active promoters marked by H3K4me2 peaks in the in the MuSC genome of 5-day-injured TA of male mice.

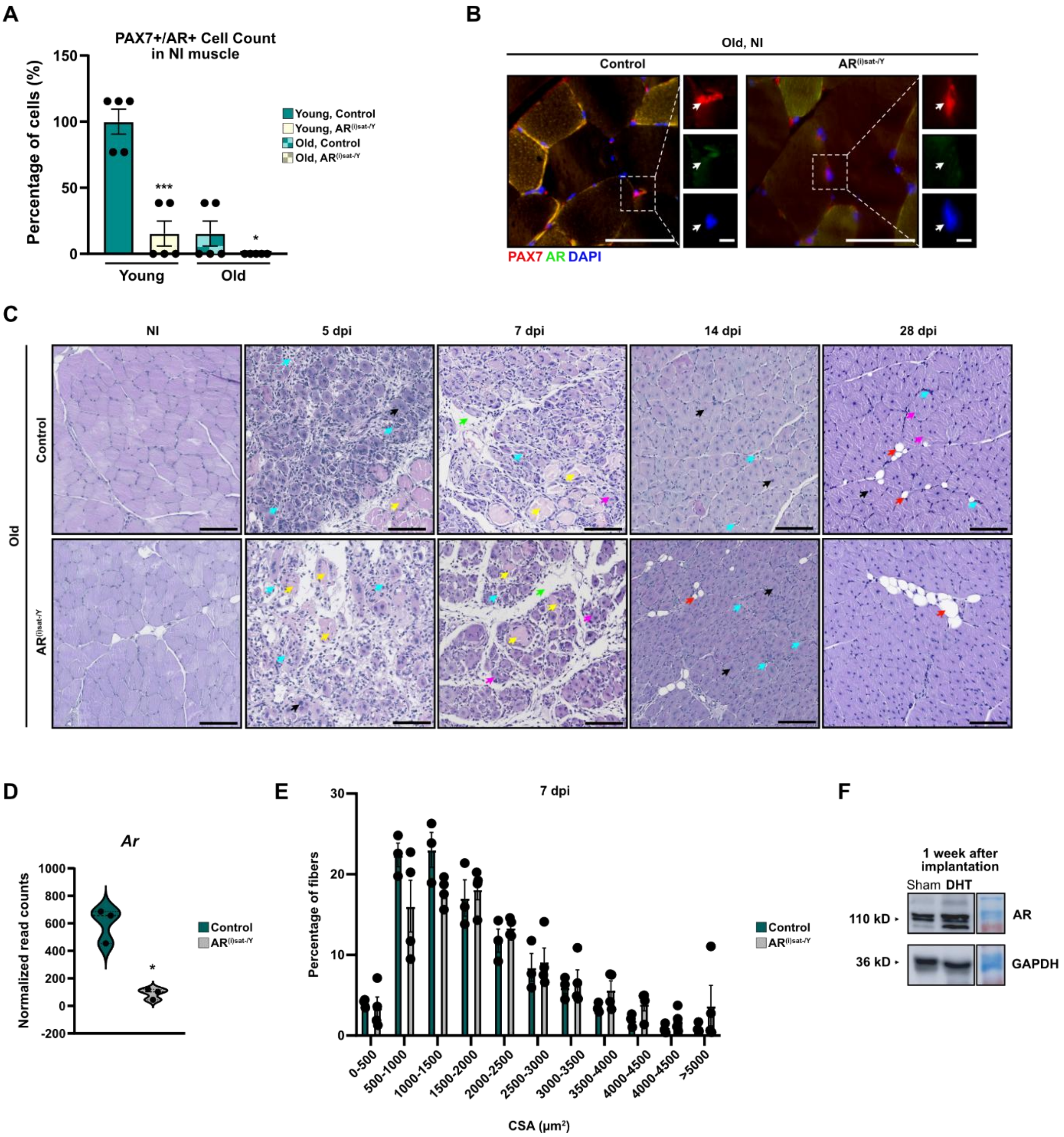


**Supplementary Figure 6: Aging effects on MuSC-AR mediated signalling**

**(A)** Percentage of PAX7/AR-double positive cells in non-injured (NI) muscles of young and old, control and mutant male mice. Data are presented as mean ± SEM. Statistical test used is Ordinary One-way ANOVA; * = $p < 0.05$; *** = $p < 0.001$.

**(B)** Representative immunofluorescent labeling of AR (in green) and PAX7 (in red) in non-injured TA of control and $AR^{(i)sat-/Y}$ old male mice. White arrows indicate double PAX7- and AR-positive cells. Yellow arrows point to AR-positive cells. White arrows denote MuSC. Nuclei were stained with DAPI. Scale bars, 100 µm for the main, 30 µm for magnified inset.

**(C)** Representative hematoxylin and eosin (H&E) staining of TA muscles of control and $AR^{(i)sat-/Y}$ old male mice in non-injured condition (NI), and at indicated time points after injury. Black arrows indicate centrally nucleated fibers. Cyan blue arrows point to immune infiltration. Yellow arrows denote necrotic fibers. Green arrows show interstitial space. Pink arrows label misshaped regenerating myofibers. Red arrows designate adipocytes. Orange arrows mark fibrosis. Scale bars, 250 µm.

**(D)** Violin plots representing the normalized read counts of *Ar*, selected from SMART-seq analysis performed control MuSC FACS-isolated from 7-day-injured TA of old and young mice. Statistical test used is Wald test; * = $p < 0.05$.

**(E)** Distribution of myofibers cross-section area (CSA) in TA muscles of old control and $AR^{(i)sat-/Y}$ male mice 7 dpi. Data are presented as mean ± SEM. Statistical test used Two-way ANOVA with Sidak's correction.

**(F)** Representative Western blot analysis of AR protein in Gastrocnemius skeletal muscles from old male mice under different conditions: sham-operated (Sham) and DHT-supplemented (DHT). Samples were analyzed at 1 week after DHT supplementation. GAPDH was used as a loading control.

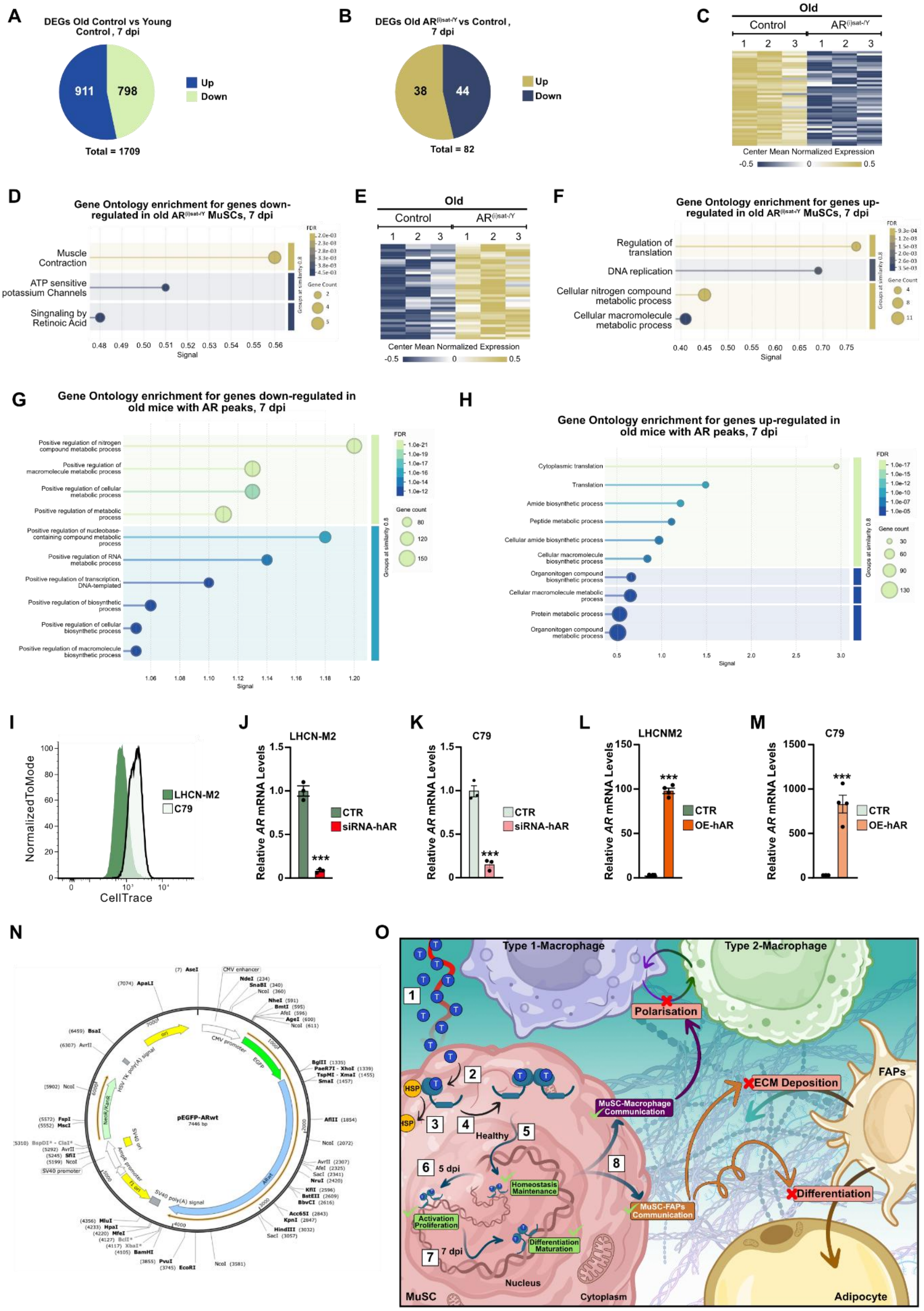


**Supplementary Figure 7:** Aging specific AR-mediated control of skeletal muscle repair

**(A)** Pie chart depicting the number of down- and up-regulated genes in control MuSC FACS-isolated from 7-day-injured TA of old versus young control mice.

**(B)** Pie chart depicting the number of down- and up-regulated genes in AR$^{(i)sat-/Y}$ versus control MuSC FACS-isolated from 7-day-injured TA of old mice.

**(C-D)** Heatmap depicting the mean centered normalized expression of down-regulated genes from SMART-seq analysis performed on AR$^{(i)sat-/Y}$ MuSC FACS-isolated from 7-day-injured TA of old mice **(C)**, and corresponding gene ontology enrichment **(D)**.

**(E-F)** Heatmap depicting the mean centered normalized expression of up-regulated genes from bulk RNA-seq analysis performed on AR$^{(i)sat-/Y}$ MuSC FACS-isolated from 7-day-injured TA of old mice **(E)**, and corresponding gene ontology enrichment **(F)**.

**(G-H)** Gene ontology enrichment analysis of genes associated with AR peaks that are down- **(G)** or upregulated **(H)** in old mice, in 7-days injured TA.

**(I)** Representative normalized CellTrace flow cytometry profiles of LHCN-M2 and C79 cells, 48 hours after CellTrace dye incubation.

**(J-M)** Relative *Ar* mRNA expression levels quantified in control and siRNA-hAR-transfected LHCN-M2 **(J)** and C79 cells **(K)**, as well as in plasmid-hAR-transfected LHCN-M2 **(L)** and C79 cells **(M)**, 24 hours post-transfection. Data are presented as mean ± SEM. Statistical test used was two-tailed Mann-Whitney test; *** = $p < 0.001$.

**(N)** Schematic representation of the pEGFP-ARwt expression plasmid (7,446 bp). Circular map of the pEGFP-ARwt vector encoding wild-type human androgen receptor (ARwt) fused to enhanced green fluorescent protein (EGFP). EGFP expression is driven by the CMV promoter and enhancer. The ARwt coding sequence is inserted downstream of EGFP, generating a fusion construct. The plasmid contains a neomycin/kanamycin resistance cassette (NeoR/KanR) under the control of the SV40 promoter for selection in mammalian and bacterial cells, respectively. Additional elements include the HSV-TK and SV40 polyadenylation signals, the pUC origin of replication (ori) for bacterial propagation, the f1 origin, and the AmpR promoter. Unique restriction enzyme recognition sites and their nucleotide positions are indicated around the plasmid backbone.

**(O)** Schematic overview illustrating the role of AR signaling in MuSC during skeletal muscle regeneration at puberty. Following its release into the bloodstream, testosterone passively diffuses into muscle tissue, where it enters cells, including MuSC, to exert its genomic effects (1). In the cytoplasm, AR remains inactive due to its association with heat-shock proteins (HSPs) (2). Upon testosterone binding, AR dissociates from HSPs (3), forms a homodimer with the ligand (4), and translocates to the nucleus (5). There, it binds to specific DNA sequences known as androgen response elements (AREs) to regulate gene transcription. In homeostatic absence of injury, AR maintains MuSC identity and quiescence by promoting the expression of genes involved in MuSC maintenance such as *Pax3* and *Pax7* (5). Upon injury, AR activity becomes spatially and temporally dynamic. For instance, at 5 days post-injury (5 dpi), AR repositions on the genome to target genes associated with MuSC activation and proliferation including *Myf5*, supporting the initial commitment phase of regeneration (6). By 7 dpi, AR shifts its chromatin binding profile toward loci governing differentiation and fusion by binding genes as *Myod1* and *Mymk*, ensuring the proper maturation of myogenic progenitors for effective tissue repair (7). This finely tuned AR signaling orchestrates a balance between proliferation and differentiation, thereby preserving the MuSC pool for future regenerative needs. Beyond its MuSC-intrinsic regulatory functions, AR signaling in MuSC orchestrates a balanced dialogue with other microenvironment-resident cell types, fostering a regenerative environment characterized by coordinated immune cell recruitment, controlled extracellular matrix remodeling, and the proper restraint of adipogenic conversion required for efficient muscle regeneration (8). AR: the androgen receptor; AREs: androgen response elements; T: testosterone; HSP: heat-shock protein; dpi: days post-injury; MuSC: muscle stem cell; FAP: fibro-adipogenic progenitor; 5 dpi: 5 days post-injury; 7 dpi: 7 days post-injury.

**Supplementary Tables**

**Supplementary Table 1: Cell Counts per Population from scRNA-seq analysis at 5 dpi**

| Cell Population | Number of cells in control | Number of cells in mutants | Total number |
|---|---|---|---|
| Macrophages | 802 | 1444 | 2246 |
| Fibroblasts | 521 | 1048 | 1569 |
| Endothelial lymphatic | 393 | 481 | 874 |
| FAPs | 340 | 505 | 845 |
| Erythocytes | 148 | 387 | 535 |
| MuSC | 189 | 256 | 445 |
| Dendritic cells | 145 | 216 | 361 |
| Myocytes | 38 | 152 | 190 |
| NK cells | 86 | 89 | 175 |
| Pericytes | 61 | 108 | 169 |
| Endothelial vascular | 35 | 61 | 96 |
| Tenocytes | 29 | 57 | 86 |
| Total | 2787 | 4804 | 7591 |

**Supplementary Table 2: List of the markers from the different populations obtained from scRNA-seq analysis at 5 dpi**

**Supplementary Table 3: List of the markers of various MuSC subclusters obtained from scRNA-seq analysis at 5 dpi**

**Supplementary Table 4: Primers for mouse genotyping**

| Gene | Forward | Reverse |
|---|---|---|
| *CreER$^{T2}$* | 5’ TTCCCGCAGAACCTGAAGATGTTCG 3’ | 5’ GGGTGTTATAAGCAATCCCCAGAAATGC 3’ |
| *ARL2/WT* | 5’ CTGGTTGCTAAGGGACTTCGG 3’ | 5’ GCCCACACAAACAGTCAGCCCA 3’ |
| *ARL-* | 5’ TGTTTTCACTGTCACTGCAGC 3’ | 5’ GCCCACACAAACAGTCAGCCCA 3’ |

**Supplementary Table 5: Primers for RT-qPCR analysis in mice**

| Gene | Forward | Reverse |
|---|---|---|
| *Pax7* | 5’ TAAGAGAGAGAACCCCGGGA 3’ | 5’ GATGCCATCGATGCTGTGTT 3’ |
| *Ar* | 5’ CTGCCTCCGAAGTGTGGTAT 3’ | 5’ GCCAGAAGCTTCATCTCCAC 3’ |
| *Tbp* | 5’ GTCCCGTGGCTCTCTTATTCTC 3’ | 5’ ATATAATCCCAAGCGATTTGC 3’ |
| *Myod1* | 5’ AGCACTACAGTGGCGACTCA 3’ | 5’ GCTCCACTATGCTGGACAGG 3’ |
| *Myog* | 5’ CACCCTGCTCAACCCCAAC 3’ | 5’ CAGCCCCACTTAAAAGCCC 3’ |
| *Myh1* | 5’ CCAAGGGCCTGAATGAGGAG 3’ | 5’ GCAAAGGCTCCAGGTCTGAG 3’ |
| *Myh2* | 5’ AAGCGAAGAGTAAGGCTGTC 3’ | 5’ GTGATTGCTTGCAAAGGAAC 3’ |
| *Sdha* | 5’ ACACAGACCTGGTGGAGACC 3’ | 5’ GGATGGGCTTGGAGTAATCA 3’ |
| *Sdhd* | 5’ GATCCCTGCTGGGTACTTGA 3’ | 5’ AAGTAGCAAAGCCCAGCAAA 3’ |

**Supplementary Table 6: Primers for RT-qPCR analysis in human**

| Gene | Forward | Reverse |
|---|---|---|
| *AR* | 5’ CTGCCTCCGAAGTGTGGTAT 3’ | 5’ GCCAGAAGCTTCATCTCCAC 3’ |
| *HSP90AB1* | 5’ CTCTGTCAGAGTATGTTTCTCGC 3’ | 5’ GTTTCCGCACTCGCTCCACAAA 3’ |
